# The 3D genomics of lampbrush chromosomes highlights the role of active transcription in chromatin organization

**DOI:** 10.1101/2025.01.27.635017

**Authors:** T. Lagunov, M. Gridina, A. Nurislamov, T. Kulikova, A. Maslova, V. Konstantinov, A. Popov, A. Krasikova, V. Fishman

**Affiliations:** Institute of Cytology and Genetics, Novosibirsk, Russia; Novosibirsk State University, Novosibirsk, Russia; Laboratory of Cell Nucleus Structure and Dynamics, Saint-Petersburg State University, Saint-Petersburg, Russia; Artificial Intelligence Research Institute, Moscow, Russia; Sirius University of Science and Technology, Sirius, Russia

**Keywords:** Chromatin architecture, chromatin domain, chromomere, cohesive cohesin, cohesion, contact domain, hypertranscription, lampbrush chromosomes, loop extrusion, oocyte nucleus, polymer simulation, single-cell Hi-C, transcription loops

## Abstract

Lampbrush chromosomes are giant meiotic bivalents in growing oocyte nuclei that have served as a classic model system for studying chromatin organization and RNA synthesis for over a century. Despite their importance, the molecular mechanisms underlying lampbrush chromosomes formation and their distinctive chromomere-loop architecture have remained poorly understood. Moreover, the influence of hypertranscription on chromatin organization during oogenesis remains enigmatic. Here, we provide the first comprehensive genomic, cytological, and biophysical analysis of lampbrush chromosome organization by integrating single-cell Hi-C, RNA-seq, NOMe-seq, FISH mapping, and chromatin simulations. Single-nucleus Hi-C analysis revealed CTCF-independent contact domains with stable boundaries defined by transcription units in a convergent orientation. Contact domains identified through Hi-C analysis correspond to insulated chromomeres in lampbrush chromosomes. Small transcriptionally inactive contact domains surrounded by transcription units in the diverged orientation form “chromatin knots”, which are often detached from the chromosome axis. Transcription loops frequently manifest as a “cross” pattern with reduced contacts within chromatin domains. Integrative analysis of the whole-genome data uncovers the mechanisms underlying lampbrush chromosome structure, revealing how hypertranscription modulates chromatin stiffness and repositions SMC complexes to establish the distinctive chromomere-loop organisation. Biophysical modeling through polymer simulation reproduces key features of lampbrush chromosomes, including transcription loop formation, chromomere compaction, and insulation patterns. These findings offer a unifying framework for understanding the remarkable chromatin architecture of lampbrush chromosomes and their transcription-dependent organization.

**Highlights:**

- First integration of single-cell Hi-C, RNA-seq, NOMe-seq and microscopy methods uncovers molecular mechanisms underlying lampbrush chromosome architecture.
- Hi-C reveals contact patterns corresponding to lampbrush chromomeres and transcription loops, validated through BAC-based FISH mapping.
- Lampbrush chromosomes are segmented into contact domains formed via a CTCF-independent mechanism, with boundaries coinciding with convergently oriented gene pairs.
- Hypertranscription shapes lampbrush chromosome through multiple mechanisms, increasing stiffness and decreasing compaction of transcribed units, generating outward pressure, pushing transcription loops away from the chromosome axis, and repositioning SMC complexes to form transcription-dependent domains with stable boundaries.
- Hi-C and RNA-seq data analysis as well as polymer simulations demonstrate that cohesive cohesin functions as a transcription-anchored barrier essential for domain insulation in lampbrush chromosomes.

**Graphical Abstract:** 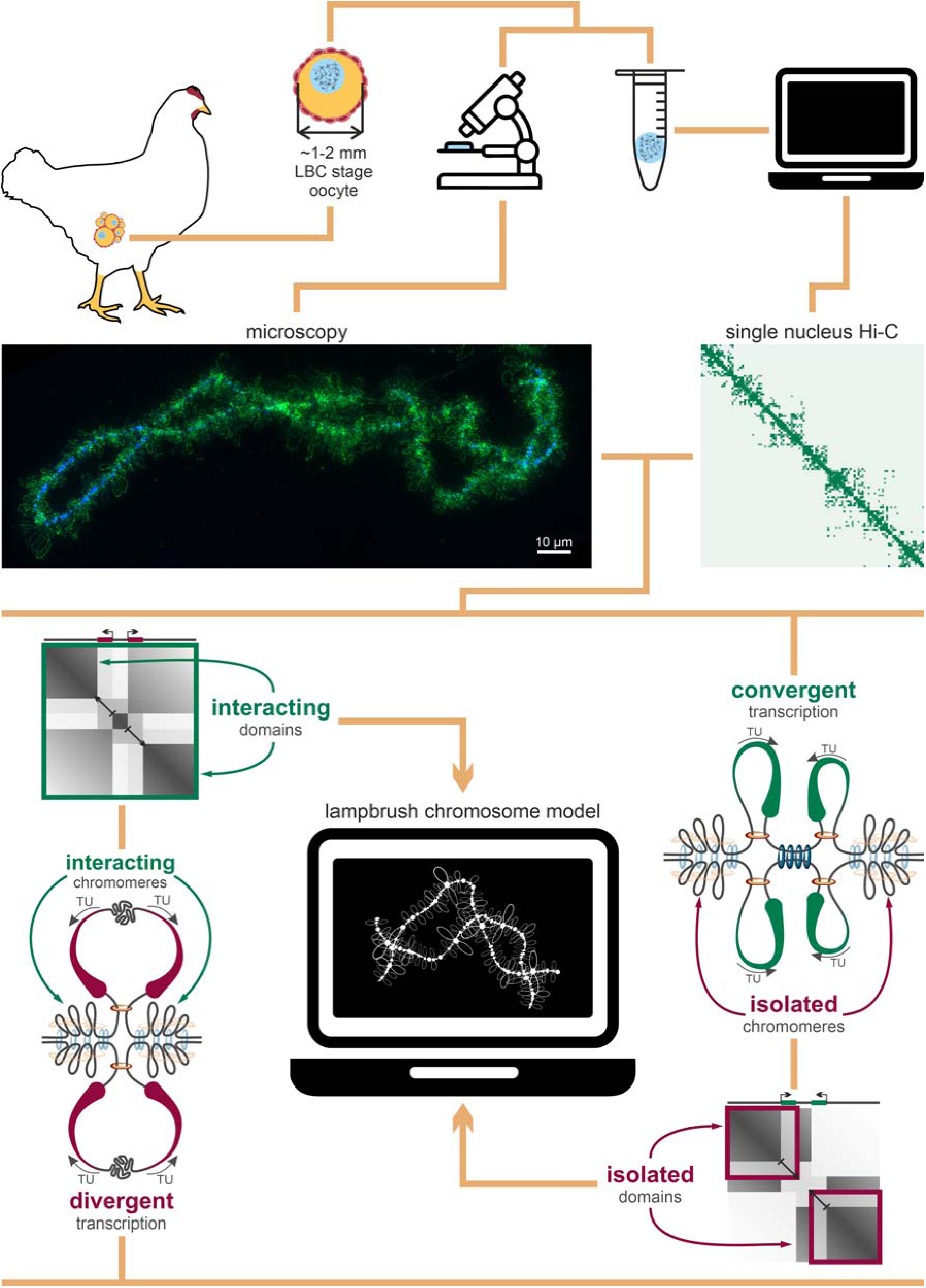

## Main

Chromosomes are folded non-randomly in the nuclear space. Their spatial organization is essential for several genomic processes (1–3), such as regulation of gene expression (4–7), DNA replication (8–10) and repair (11), and mechanical properties of the nucleus (12, 13). Consistent with this pivotal role in genome regulation, spatial organization of chromatin was shown to be conserved across large evolutionary distances of hundreds of million years (14–17).

There are several essential mechanisms responsible for chromatin organization in animal cells. First, DNA is a polymer organized in the nuclear space according to the principles of polymer physics. This implies that frequency of contacts between loci decreases with increase of genomic distance, following power law. For the majority of the studied cell types, scaling of contacts with genomic distance is concordant with the “crumpled” states of the chromatin (18), characterized by largely unknotted conformations with a fractal dimension of ∼3 (19). Second, multivalent homotypic interactions between proteins cause segregation of the chromatin into compartments with distinct epigenetic states (20–24). Moreover, active and inactive chromatin is directed towards different nuclear landmarks (2). Inactive chromatin is often associated with nuclear lamina and nucleolar periphery, while active chromatin is in proximity to the interchromatin space comprising nuclear speckles (25–27). Third, ATP-dependent extrusion of DNA by SMC-complexes cohesin and condensin results in chromosome compaction and increased probability of long-range interactions (1, 28, 29). The extrusion process can be modulated by several factors. In the majority of vertebrate cell types, CTCF protein blocks cohesin-mediated extrusion, therefore CTCF binding sites demarcate compact self-interacting genomic regions, often referred to as topologically associating domains (TADs) (1, 30–33). Interactions of cohesin with other proteins, including replication-specific MCM (34) or transcription-associated RNA-polymerase II complexes (35–38) also may modulate extrusion, although details of these mechanisms are less studied. Moreover, interaction of cohesin with MCM and RNA-polymerase have less impact on genomic contacts compared to cohesin-CTCF interactions, and their effect becomes pronounced only when artificially modulated (37) or in analysis of specific cell cycle stages (34).

Although chromatin contacts in the majority of vertebrate cells can be explained by above mentioned mechanisms with relatively high accuracy (39, 40), specific cell types or cell cycle stages show distinct arrangement of chromatin (2). This includes early stages of development, prior to or immediately after zygotic genome activation, and germ cells (41). Importantly, chromatin architecture is uniquely reorganized during the oocyte-to-zygote transition. In mammals, features of genomic organization at diplotene stage include compartments, TADs, and loops (41, 42). Although these chromatin domains can be detected when averaged over the genome, the presence of each type of domain at a locus varies between cells (42). TAD boundaries in murine oocytes are located concordant with somatic cells, demarcated by CTCF, and associated with cohesin-mediated loop extrusion (42). Compartments in murine diplotene oocytes manifest as plaid-pattern of Hi-C contacts and demonstrate strong association with distribution of H3K27me3 mark (41). Compared to somatic cells, in oocytes interactions within compartments are restricted to the regions located less than 10 Mb away from each other, presenting a “local compartmentalization” pattern (41). Similar depletion of long-range interactions and local compartmentalization was also observed during meiotic division of sperm cells (43, 44). Strength of both TADs and compartments gradually decreases with oocyte transition from early to late diplotene stage, and these structures disappear at metaphase-II (MII) stage (41, 42).

Previous whole-genomic studies focused on chromatin organization in oocytes did not include representatives of vertebrate taxa in which chromatin undergoes lampbrush chromosome state. Lampbrush chromosomes (LBCs) represent a specific form of nuclear chromatin that emerges at the diplotene stage of oogenesis in the majority of vertebrate species, including amphibians, fish, reptiles, birds, monotremes, and certain insects. In the 3D space of the giant oocyte nucleus, lampbrush chromosomes do not contact each other, are not tethered to the nuclear periphery, and do not have a preferred radial position (45, 46). Lampbrush chromosomes consist of separate chromatin globules connected by a thin thread and protruding lateral loops, together forming chromomere-loop complexes (47–49). The unusually enormous size of LBCs allows for chromomere-loop structure visualization using conventional light microscopy, making diplotene oocytes a preferred model for microscopy-based studies of chromatin architecture.

The phenomenon of LBC organization is attributed to hypertranscriptional activity of the oocyte nucleus (50, 51). Hypertranscription results in formation of lateral loops tightly covered by elongating RNA polymerases and nascent transcripts (49, 52, 53) and containing single or multiple genes essential for maternal products biosynthesis (54–57). Another structural unit of LBC are lampbrush chromomeres – compact insulated chromatin globules present along LBC axes (48). Chromomeres do not clearly correspond to any of the known spatial chromatin domains of the interphase nucleus. Genomic loci of neighbouring somatic TADs can be detected as a part of a single chromomere, while genomic regions belonging to one somatic TAD can be detected as a part of two adjacent chromomeres (55, 58). There is, however, partial correspondence between gene-poor heterochromatic regions of constitutive B-compartments identified in somatic cells and clusters of large DAPI-positive chromomeres, which are also enriched for inactive chromatin marks (58–62). Overall, chromatin organization in LBCs drastically differs from somatic cells.

Hypertranscription being a hallmark of LBCs is also a widespread phenomenon common in the other cell types, including developmental hypertranscription and hypertranscription in adult stem/progenitor cells (63, 64). Our work aims to investigate the influence of hypertranscription on chromatin organization using LBCs as a model system.

Following this aim, we present the first Hi-C analysis of lampbrush chromosomes, supplemented by transcriptomic and epigenomic profiling. We show that chromatin contacts in LBCs display unique features, including restriction of long-range and inter-chromosomal interactions and formation of CTCF-independent chromatin domains and loops. We show strong correlation between location and orientation of transcription units and formation of chromatin domain boundaries. Based on our observations, validated by FISH and other complementary analysis, we develop a physical model of LBCs explaining LBC architecture via interplay between SMC and RNA-polymerase complexes. We show that the polymer model can qualitatively explain observed Hi-C patterns genome-wide and accurately reproduce spatial configuration of individual loci. Finally, the developed model highlights the importance of the transcription-mediated ribonucleoprotein matrix accumulation, nucleosome removal, and cohesive cohesin repositioning as factors determining spatial structure of LBC.

## Methods

### Oocytes isolation

Chicken oocytes, with diameters ranging from 1 to 2 mm as lampbrush chromosome and from 4 to 7 mm as post-lampbrush chromosome stages, were dissected from the ovary of healthy chicken and placed in individual drops of cooled "5:1" medium (83 mM KCl, 17 mM NaCl, 6.5 mM Na2HPO4, 3.5 mM KH2PO4, 1 mM MgCl2, 1 mM DTT, pH 7.2). All animal experiments were approved by the bioethics committee of the Institute of Cytology and Genetics SB RAS (Protocol №66, October 9th, 2020). International guidelines were followed during experimental procedures (“Guide for the Care and Use of Laboratory Animals” (65)). Nuclei were isolated as described earlier (45), washed with "5:1" medium, and transferred to a dish containing 2% formaldehyde in PBS for fixation at room temperature for 15 minutes. The formaldehyde was quenched by adding glycine to a final concentration of 0.125 M for 15 minutes at room temperature. The integrity of the nuclei was assessed using an Olympus stereomicroscope, and the Hi-C protocol was initiated immediately.

### Hi-C libraries preparation

Individual fixed nuclei were transferred to microplate wells containing 9 μL of ice-cold lysis buffer (10 mM Tris-HCl pH 8.0, 10 mM NaCl, 0.2% Igepal) for 20 minutes. The nuclei were then washed with NEB3.1 + 0.5% SDS and transferred to 0.2 mL PCR tubes with 3 μL of NEB3.1 + 0.5% SDS. 10 μL freshly prepared 1.2% Low Melting Point Agarose were added to each tube under a stereomicroscope. After solidification, 10 μL of NEB3.1 + 0.6% SDS was added, and the samples were incubated at 37 °C for 1 hour. SDS was quenched using 10 μL of 6% Triton X-100, and chromatin fragmentation was performed with 25 U of *Dpn*II at 37 °C overnight.

The digested chromatin ends were labeled using 5 U of the Klenow fragment in the presence of biotin-15-dCTP at 22 °C for 4 hours, followed by overnight ligation at 16 °C. Crosslinks were reversed by incubation at 65 °C overnight, and the Low Melting Point Agarose was digested with 0.4 U Agarase for 1 hour at 42 °C. The purified DNA was fragmented with *Alu*I for 1 hour at 37 °C, and next-generation sequencing (NGS) libraries were prepared using the KAPA HyperPrep kit with modifications. The volumes for the end repair, adapter ligation, and PCR reactions were reduced by 5-fold, 5-fold, and 2-fold, respectively, and adapters were diluted to 300 nM. A biotin pull-down was performed with Dynabeads MyOne Streptavidin C1 beads after the adapter ligation step, followed by 25 cycles of PCR. Genomic libraries were sequenced using BGI DNBseq G400 machine in 2×150 paired-end mode.

### Hi-C data analysis and quality control

The illumina adapters from raw sequences were cut with cutadapt ver-4.1 (66). Data were aligned with using juicer ver-1.5.6 (67) script against chicken telomere-to-telomere genome assembly (ASM2420605v2, (68)). The read pairs without duplicates were filtered by mapping quality >= 30. Quality statistics were computed using juicer output with slight modifications in cis/trans ratio estimation described previously (69). We performed deep sequencing only for samples with duplicates percentage below 30% (estimated in test run with less than 200k read pairs sequenced) and fraction of intra-fragment reads percentage 20% (see Supplementary Note 1 for additional details).

We used juicer output to generate mcool files with cooler ver-0.10.2 (70). Due to the low number of contacts in the dataset (∼500’000 per cell), we didn’t use the cooler balance function. Instead, we set weights for bins with zero coverage as “nan” and weights of bins with non-zero coverage as 1. Aggregated plots were made with coolpuppy ver-1.1.0 (71) with “maxshift” of 2 MB and “local” flag. The P(s) curves were calculated using cooltools (72) gaussian smooth sigma value equal to 0.1.

The raw reads from (42) were aligned against GRCm39 genome assembly and processed the same way (except cooler balance was used for mcool maps). The raw reads from (15) were aligned against ASM2420605v2 genome assembly and processed the same way (except cooler balance was used for mcool maps).

### Domain boundaries annotation

To annotate domain boundaries for both chicken oocytes, mouse oocytes and chicken somatic cells the lavaburst ver-0.2.0 was used (https://github.com/nezar-compbio/lavaburst). This algorithm divides the genome into domains in such a way as to maximize a global domain scoring function. The algorithm depends on *gamma* hyperparameter which should be found empirically. We found the value of gamma separately for each single oocyte the same way as in the (42): 1) the 40 values of gamma in range from 6 to 24 were scanned; 2) the value of gamma was multiplied by chromosome size and divided on (3250 * resolution); 3) the average domain size was calculated for each gamma (taking into account only domains more than 200 Kb); 4) the gamma value that makes the average domain size closest to 500 Kb for mouse NSN oocytes was used. For chicken oocytes the preferable average domain size was 1 Mb (motivated by the visual assessment of Hi-C maps) and the gamma range was from 0.5 to 10.5. Both analyses (for mouse and chicken oocytes) were performed on resolution 20 kb.

To assess the stability of domain-boundary positions at single-cell resolution, we computed an F1 score for boundary concordance. Boundaries on the same chromosome were counted as true positives (TP) if their coordinates differed by ≤80 kb; boundaries without a counterpart within this window contributed to false positives (FP) and false negatives (FN). The F1 score was calculated as F1 = 2*TP / (2*TP + FP + FN). For cross-species comparison (chicken vs mouse), we normalized the F1 by a permutation baseline: for each single oocyte, domain intervals were randomly re-positioned along each chromosome while preserving the total length of inter-domain gaps, and 10 such permutations were generated per oocyte. For each oocyte pair, the permutation baseline was defined as the mean F1 across the corresponding permutation replicates. The normalized pairwise F1 was then obtained by dividing the observed pairwise F1 by this baseline.

The domains were divided into LUDs and SSDs by N50 threshold: all domains were sorted by their size and the domain size that covers the half of the summed domain length was set as threshold.

### Hi-C map Insulation score track

Insulation function from cooltools package ver-0.5.1 (72) has been used for calculating the Insulation scores on merged oocytes. Resolution size is 16 Kb and window size is 2 Mb.

### Cross Pattern Score track

“Cross Pattern Score” (CPS) for genome position n was calculated using the equation:

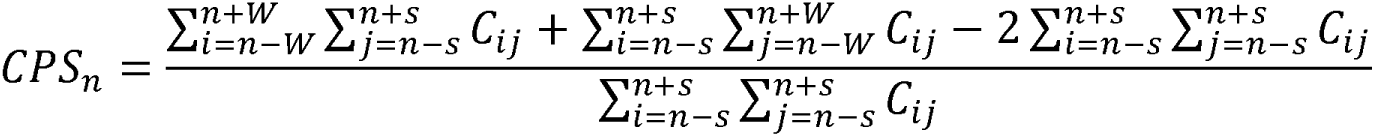

Where s = 35 kb, W = 4 Mb, and Cij is the number of contacts between loci i and j. The resolution of the calculations was 1 kb. A schematic of the score computation is provided in the Supplementary Figure 1.

### Aggregated Hi-C maps analysis

The aggregated Hi-C maps were generated using the coolpuppy package (https://zenodo.org/badge/latestdoi/147190130). For normalization, the coolpuppy program utilized random shifting, a built-in feature designed to adjust positional biases in the data. Insulation scores for the aggregated maps were computed with this package by calculating the relative mean signal in the first and third quadrants compared to the second and fourth quadrants. Higher insulation scores indicate higher levels of spatial separation in this context.

### Saddle strength analysis

To quantify A/B compartment interaction strength, we performed a saddle plot analysis based on the first principal eigenvector of the Hi-C contact matrix derived from chicken embryonic fibroblasts. This eigenvector was computed at 100 Kb resolution using the *cooltools* package (version 0.7.1) and served as a reference compartment annotation for all subsequent analyses. Eigenvector values were divided into 40 percentile-based bins. To minimize the influence of extreme regions with ambiguous compartment identity, bins corresponding to the highest and lowest mean eigenvector values were excluded from further analysis. Saddle plots were generated by calculating the observed-over-expected contact frequencies between each pair of bins. Compartment strength was defined as the ratio of mean intra-compartment interactions (AA + BB) to inter-compartment interactions (AB + BA). All computations were performed using functions implemented in *cooltools* (ver-0.7.1).

### Chicken liver ChIP-seq data preparation

The CTCF ChIP-seq data was taken from (73) as sequenced reads. The reads were aligned at ASM2420605v2 chicken genome assembly (68) and bamCoverage ver-3.5.4 were used to get bigwig files. The ChIP signal bigwig was divided on input signal bigwig for each bin and logarithm from results were taken.

### NOMe-seq experiments and data analysis

Nuclei were isolated and washed as described above. Individual nuclei were transferred immediately to 3 μL of GC Reaction Buffer (NEB) with 0.2% Igepal for 20 minutes at RT. Reaction of methylation was performed in 1x GC Reaction Buffer supplemented with 160 µM S-adenosylmethionine with 6U GpC Methyltransferase at 37 °C 1 hour. Proteines were digested by proteinase K at 65 °C for 2 hours. Libraries for whole genome bisulfite sequencing were prepared using Pico Methyl-Seq Library Prep Kit (Zymo Research) according to the user manual. For control experiments we also supplemented chicken oocytes nuclei with human K562 cells.

Genomic libraries were sequenced using the BGI DNBseq G400 machine in 2×150 paired-end mode.

The illumina adapters from raw sequences were cut with cutadapt ver-4.1 (66). To process NOMe-seq reads, we used Bismark version 0.23.0 (74, 75) in conjunction with Bowtie2 version 2.4.4 (76–79), setting the -t parameter to 10. Reads were aligned to the ASM2420605v2 genome assembly (68), and any reads with MAPQ < 10 were excluded. Methylation data from the aligned reads were then converted to the *.bedgraph* format using the bismark-methylation-extractor.

The GpC methylation profile was obtained from the NOMe-seq experiment described above. The CpG methylation profile was obtained from the NOMe-seq experiment and merged with the profile from the Methyl-seq experiment from (80).

### RNA-seq experiments and data analysis

Nuclei were isolated and washed as described above. Individual nuclei were frozen immediately. Total RNA was isolated from the nuclei using TRIzol reagent (ThermoFisher Scientific) according to previously described protocol (57). The RNA samples were sent to BGI company to generate transcriptomic libraries using experimental workflow including the following steps: 1) oligoT-enrichment; 2) RNA fragmentation; 3) first-strand synthesis with random primers; 4) second strand synthesis with dUTP instead of dTTP; 5) end-repair, 3’-adenylation and adaptors ligation; 6) Uracil-DNA-Glycosylase treatment followed by PCR amplification and circularization; 7) Sequencing on DNBSEQ in PE100 paired-end mode.

Together with poly(A)-enriched stranded RNA-seq data generated in this study, the total RNA stranded RNA-seq data from (57) were used. The Illumina adapters from raw sequences were cut with cutadapt ver-4.1 (66). All data were aligned with hisat2 ver-2.2.1 (81) on ASM2420605v2 genome assembly (68). Bam files were prepared using samtools ver-1.15.1 (82) and bigwig files were made using bamCoverage ver-3.5.4 (https://github.com/BGI-shenzhen/BamCoverage). Each bam file was split on subfiles by strand (the direction of transcription) and duplicates were marked with “samtools markdup”. The unique reads were merged from all experiments (still grouped by strand) and *de novo* LBC transcript structure recovery was made with stringtie ver-2.2.2 (83). The forward and reverse transcription files were separately used with stringtie and called as “FORW” and “REV” respectively. Maximum gap allowed between read mappings was set as 300 bp; minimum assembled transcript length = 1000 bp; minimum junction coverage = 5; minimum anchor length for junctions = 10 bp; minimum isoform fraction = 0.05; minimum reads per bp coverage to consider for single-exon transcript = 3.5; minimum reads per bp coverage to consider for multi-exon transcript = 1.2.

For the expression analysis, stranded RNA-seq data of the oocyte nuclei from this work and (57) was compared with the RNA-seq data of the chicken embryonic fibroblasts from (84). All raw reads were aligned on GRCg6a genome assembly. The TPMCalculator ver-0.0.3 was used to calculate TPM values for genes of interest for ncbiRefSeq.2020-04-01, ensGene.v101 and refGene annotations (see Supplementary Table S4).

### Transcriptional activity analysis

The FPKM value was calculated for each gene and used as a measure of transcriptional activity in analysis.

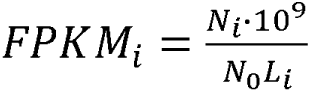

where *i* denotes the transcript index, *N_i_* - number of reads mapped to transcript *i*, *N*_0_ - total number of mapped reads, *L_i_* - length of transcript *i*. To estimate the amount of nascent RNA connected to the elongating RNA polymerases, the values of intron FPKM and exon FPKM were calculated separately for each transcription unit annotated. The minimum transcript FPKM value was used for the further analysis.

For a block of consecutive co-directional genes (a tandem, *t*), transcriptional activity was summarized by a length-weighted mean:

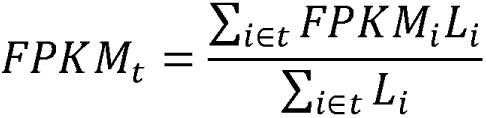

Genes and their tandems were then stratified into four classes by median splits on their length and transcriptional activity: long strong, short strong, long weak and short weak.

### Mitotic metaphase chromosomes preparation

Mitotic metaphase chromosomes were prepared from chicken embryonic fibroblast (CEF) culture. Cell cycle was arrested by 0.1 μg/ml colcemid (Biological Industries) for 5 hours, then the cells were fixed in cold 3:1 ethanol:acetic acid mixture. The CEF suspension was dropped onto the slides which had been heated in a water bath to 60°C and then air dried.

### Lampbrush chromosomes preparation

Lampbrush chromosome spreads were prepared according to a well-described procedure (85). The experimental procedure was approved by the Ethics Committee for Animal Research of St. Petersburg State University (protocol #131–04-6 dated 25 March 2019). Nuclei were manually isolated from oocytes (1-2 mm), one by one, using tungsten needles under a stereomicroscope Leica S9D or M165C (Leica Microsystems) in the isolation medium “5:1” (83 mM KCl, 17 mM NaCl, 6.5 mM Na2HPO4, 3.5 mM KH2PO4, 1 mM MgCl2, 1 mM dithiothreitol, pH 7.2). The isolated nuclei were then washed in the hypotonic solution (“5:1” medium diluted 4 times with the addition of 0.1% formaldehyde) and transferred to the isolation chamber mounted on a microscope slide and filled with the hypotonic solution. Nuclear envelope was broken manually with a pair of thin tungsten needles to release the nuclear content and slides were left to disperse for 15-20 minutes at +4°C. Slides were then centrifuged at 1600 g for 30 minutes at +4°C in a cytology centrifuge (Hettich Universal 320R), fixed in 2% formaldehyde in PBS for 30 minutes and dehydrated in a series of ethanol solutions of increasing concentrations. Specimens were mounted in an antifade solution containing 1 mg/ml DAPI (Sigma), 55% glycerol, 10 mM Tris-HCl, 2% DABCO (Sigma) and were imaged prior to FISH procedure.

### Preparation of DNA-probes

To obtain DNA probes for FISH mapping of the selected scHi-C chromatin domains, 29 BAC clones were selected: 14 BAC clones in a region (55-50 Mb) on chromosome 1; 2 and 7 BAC clones in two regions (50-53 Mb and 70-73 Mb) on chromosome 4, respectively; and 6 BAC clones in the region 3-7 Mb of chromosome 13 from the chicken BAC-library CHORI-261 (https://bacpacresources.org/chicken261.htm) (Supplementary Table S3). BAC DNA was isolated from night culture by standard alkaline lysis. DNA was labeled by nick-translation (86) with dUTP-bio (DNA-Synthesis), dUTP-dig (Jena Bioscience) and dUTP-Atto637 (Jena Bioscience) using DNA polymerase I (ABclonal) and DNAse I (New England BioLabs). Labeled DNA probes in various combinations (up to 10 probes in a single mixture) were precipitated with salmon sperm DNA and, in case of DNA probes to the region of chromosome 1, with the C0t-5 fraction of chicken genomic DNA, and dissolved to final concentration of 20 ng/μl for each DNA probe in the hybridization buffer containing 50% formamide, 10% dextran sulfate and 2⍰SSC.

### Fluorescent in situ hybridization

Chromosomal position of each BAC clone-based DNA probe was first verified by DNA/DNA FISH on CEF metaphase chromosomes: preparations were pretreated with RNAse A (Thermo Scientific) and pepsin (Sigma-Aldrich). For lampbrush chromosomes, the DNA/DNA+RNA FISH and RNA FISH procedures were used, omitting the RNAse treatment (56, 87). Chromosome preparations were denatured in 70% formamide in 2⍰SSC at 70°C for 10 minutes, and dehydrated in a series of cold (−20°C) ethanol solutions of increasing concentration. Hybridization mixtures were denatured at 90°C and cooled on ice and applied to the slides with the denatured chromosomes. Hybridization was carried out overnight in a humidity chamber at 37°C, followed by a post-hybridization wash in 0.2⍰SSC at 60°C. Biotin was detected with Alexa488-conjugated streptavidin (Jackson ImmunoResearch Laboratories) and biotinylated anti-streptavidin antibody (Vectorlabs), and digoxigenin was detected with Cy3-conjugated mouse anti-digoxigenin antibody (Invitrogen) and Cy3-conjugated anti-mouse IgG (Jackson ImmunoResearch Laboratories).

### Immunostaining

Lampbrush chromosome preparations were immunostained with mouse monoclonal antibodies against phosphorylated serine 5 of CTD repeat YSPTSPS of RNA polymerase II [H14] (Abcam, ab24759), followed by the detection with Alexa-488-conjugated goat anti-mouse IgG (Molecular Probes). Immunofluorescence staining procedure was described previously (88, 89).

### Microscopy and image processing

Chromosome preparations were analyzed with the fluorescence microscope Leica DM4000 (Leica Microsystems, GmbH) equipped with a monochrome CCD camera (1,3 Mp) and corresponding fluorescence filter cubes. LAS AF software (Leica Microsystems, GmbH) was used for image acquisition, and Adobe Photoshop CS2 (Adobe) and ImageJ (http://rsbweb.nih.gov/ij/) was used for image processing and layout. Schematic drawings were created in CorelDRAW X5 and represent an approximation of the FISH mapping of DNA probes to lampbrush chromatin domains seen in at least 4 micrographs for each DNA probe.

### Polymer Modelling

We used polymer modeling based on the Polychrom tool (90) to gain insights into the roles of individual RNA polymerases in the folding of lampbrush chromosomes. The simulation was performed in the coarse-grained model approximation, where clusters of nucleosomes and individual polymerases acted as beads.

### 1D

The modeled object is a one-dimensional array of N beads of sizes ranging from 45 to 225 bp, with dynamically loaded-unloaded RNA polymerases and active extruders (loop extrusion factors, LEFs) and non-removable cohesins without the loop extruding activity (loaded with the density of 45 Kb).

The number of base pairs in a bead differs for TUs where RNA polymerases are located and for transcriptionally inactive regions with a dense fit of nucleosomes. The size of 45 bp per bead was used to represent one RNA polymerase (∼15 nm, (91)). The regions between the TUs are condensed by the nucleosomes ∼5 times (because ∼140 bp entangle it and ∼70 bp are between nucleosomes). Our estimation is also close to the estimation from (92). Transcription units are placed according to the annotation obtained from RNA-seq data.

RNA polymerases are represented as objects with the following properties: they are loaded at the beginning of TUs with a probability proportional to the activity of the TU (estimated from FPKM metric), from where they begin movement at a speed of *vp* = 100 bp/sec (93) (the speed of LEFs is *vL* = 625 bp/sec for the naked DNA [(94)]) along the transcript. Nascent RNA fiber growth from each RNA polymerase at length *vp*dt* for the time step of *dt* seconds. The introns are excised from the pre-mRNA attached to the RNA polymerases. If the bead in the direction of movement is occupied by a LEF or cohesive cohesin (i.e. SMC), then RNA polymerase pushes the SMC (37) and all subsequent SMCs in the direction of movement until they collide with another RNA polymerase.

The mechanism of LEF action is as follows. It is randomly loaded onto a pair of adjacent beads that are not occupied by another LEF or RNA polymerase. Each time one of the two LEF retaining components can reattach to the next bead in the corresponding direction releasing the occupied one. The direction of movement changes every timestep (effectively two-way movement). Each part of the LEF can move independently of the other. If the next bead is occupied by another LEF then no movement occurs. If the next bead is occupied by RNA polymerase then the LEF can bypass the RNA polymerase with the stable probability *Pbypass*. The cohesive cohesin can be pushed by LEF if the next bead is not occupied. Extruding continues until the LEF is removed from the chromatin. The probability of removal is (*1 / lifetime)* for each timestep.

Some parameters for these LEFs were determined by using P(s) data from the experiment Hi-C data on lampbrush-stage oocyte nuclei (see Supplementary Methods 1. Modeling): *lifetime* = 1 Mb (∼60% of LEFs will pass this distance on naked DNA), *average_loop_size* = 90 Kb (determines the density of loaded condensins on chromatin), *Pbypass =* 0.01 (as in (37) article).

### 3D

The next step after forming 1D snapshots of the SMCs and RNA polymerase locations on the chromatin is to transfer the interactions into 3D. For this purpose all beads are grouped into bead clusters of 10 which include nucleosomes and transcription units. The sizes of the clusters vary from 450 to 2250 bp. New beads are placed into the starting conformation form of two parallel 2D fractals, corresponding to the two sister chromatids. The distances between beads in 3D varied according to the fractal dimension of the site (Supplementary Methods 1. Modeling).

With the launch of a three-dimensional simulation, at each moment in time, a spring of a fixed size (∼40 nm (95)) is stretched between each pair of beads connected by a given LEF, and the beads move according to a random Brownian motion.

RNA tails from 1D simulation are converted into increased bead sizes by the Marco and Siggia model (96). The repulsive potential between beads is realized as *0.5*k*(r1+r2)^2*.

A more detailed analysis of the model and parameters used is presented in the Supplementary Methods 1. Modeling.

In order to reproduce the patterns observed in LBC under microscopy, the original three-dimensional regions of the chromosome were flattened to 1 µm (see Supplementary Methods 1. Modeling).

LBC chromatin simulations were additionally validated using obtained FISH-mapping data in individual chromosomal regions (see Supplementary Note 3).

The 3D simulated structures were visualized using *plotly* (97). Online Supplementary files can be found via link (https://genedev.bionet.nsc.ru/ftp/by_User/tlagunov/LBCs/htmls/) or be downloaded from git (https://github.com/genomech/LBC_model).

### Simulated Hi-C

The resulting 3D polymers were used to construct contact maps similar to Hi-C maps. The beads of different sizes are correctly converted to Kb resolution that allow us to compare simulated and experimental contact maps of the specific chromatin slices. Since the contact map collected from one polymer conformation is essentially one conformation of an oocyte, the sum of contacts from one "*Dpn*II site" in a bead was normalized to 8 as in the experiment (see Supplementary Methods 1. Modeling for the details). Simulated Hi-C maps were then stored in mcool format and the analyses that are the same with experiment Hi-C maps were performed.

## Results

### Isolation and quality control of chicken diplotene oocytes

To investigate chromatin organization in avian lampbrush chromosomes, we selected chicken oocytes at the lampbrush stage based on follicle size. Individual nuclei were isolated and processed using a previously established single-cell Hi-C protocol (98) (Figure 1A). Multiple quality control measures were conducted to validate the integrity and reliability of the resulting data.

**Figure 1.**
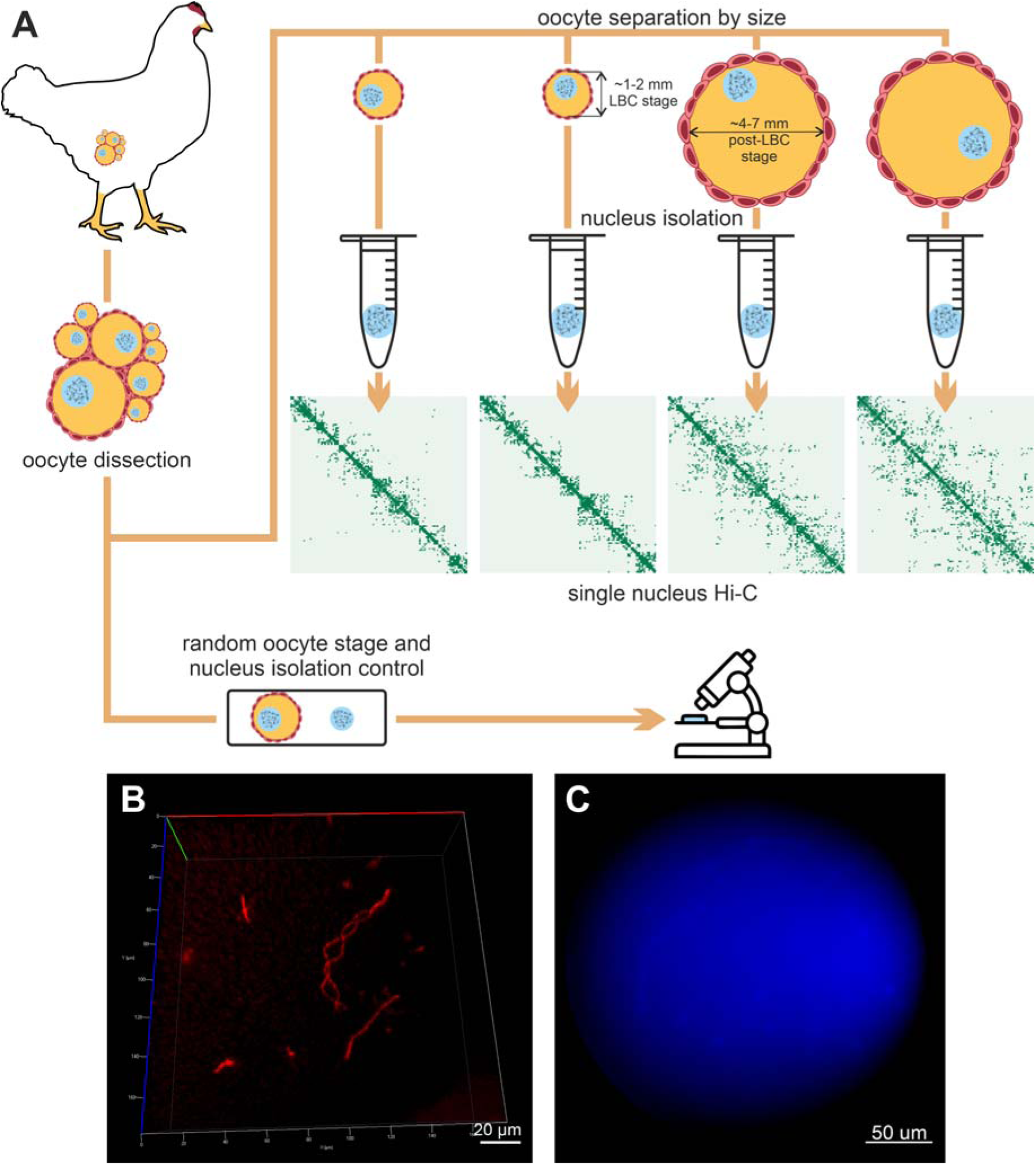
Experimental workflow preserves structural integrity of chicken lampbrush chromosomes. A) Experimental workflow, including oocyte dissection, oocyte separation by size (lampbrush chromosome stage oocyte: 1–2 mm diameter; post-lampbrush chromosome stage oocyte: 4–7 mm diameter), microsurgical isolation of nuclei, and preparation of single-nucleus Hi-C libraries. Selected oocytes and nuclei were randomly chosen for quality control.B) Lampbrush chromosomes preserve their 3D-structure after formaldehyde fixation. The entire nucleus is visualized after fixation and staining with SYTO 61. C) Isolated DAPI-stained chicken oocyte nucleus (no evidence of contamination with other cell nuclei).

To confirm that formaldehyde fixation preserves the lampbrush chromosome structure, we stained fixed nuclei with SYTO 61 dye and visualized them using confocal microscopy. The analysis demonstrated the characteristic lampbrush morphology of the captured chromosomes (Figure 1B, Supplementary Figure S2A), consistent with previous findings (45).

Next, we ensured that oocyte genomic material was not contaminated by somatic cells. Chicken follicles include various cell types, such as erythrocytes, which are significantly smaller than oocyte nuclei. To exclude contamination, nine randomly selected oocyte nuclei were stained with DAPI post-isolation and visually inspected. Microscopy analysis revealed no DAPI signals outside oocyte chromosomes (Figure 1C). As a control, erythrocytes were manually added to oocyte samples, confirming that erythrocyte nuclei are detectable during visual inspection (Supplementary Figure S2B).

To further validate the absence of contamination, we leveraged the fact that a single oocyte nucleus contains no more than four copies of each genomic fragment. Consequently, no more than eight unique interactions should be detected per *Dpn*II restriction fragment (two ends per fragment, (99)). Genomic fragments exhibiting more than eight interactions would indicate contamination or Hi-C artifacts. Our analysis showed that the vast majority of fragments exhibit fewer than nine interactions. Comparison with the bulk Hi-C data downsampled to the same sequencing depth shows that the portion of fragments with fewer than nine interactions in chicken oocytes is lower than expected based on total Hi-C contacts count (Supplementary Figure S2C). Finally, the Hi-C contact profiles of chicken oocyte nuclei display a characteristic pattern distinct from that of typical somatic cells (described below). Taken together, these results confirm that the isolated oocyte nuclei were free of contamination from other chicken cell types.

Single-nucleus Hi-C experiments were performed on a total of 96 oocyte nuclei. Initial library quality was evaluated through shallow sequencing (∼200,000 reads per sample), using metrics such as the fraction of unmapped reads, duplicate reads, intra-chromosomal reads, and intra-fragment reads (see Methods). Based on these criteria, 25 single-cell oocyte libraries were selected for deeper sequencing (∼3–40 million read pairs per nucleus) and downstream analysis (Supplementary Table S1).

The resulting Hi-C maps cover of >80% bins with at least one interaction at 4 Kb resolution, with a few exceptions where one or more chromosomes were completely absent but the remaining genome showed consistent coverage (Supplementary Figure S2D). Deeply sequenced Hi-C maps displayed a high degree of similarity across individual oocyte nuclei (Figure 3A-B), confirming the high quality of the obtained data.

### Lampbrush chromosomes form elongated structures with minimal inter-chromosomal interactions

Lampbrush chromosomes at the diplotene stage of meiosis are known to form elongated structures that are spatially separated within the giant oocyte nucleus (45). This spatial arrangement is also preserved after fixation, as shown in Figure 1B and Supplementary Figure S2A. Hi-C data analysis further confirms these unique features of LBCs.

Our results show a strikingly low frequency of inter-chromosomal (trans) contacts, with only 0.2–1% of spatial interactions occurring between non-homologous chromosomes (Figure 2A, Supplementary Table S1). This is markedly lower than the ∼10% reported for chicken fibroblasts ((15), Figure 2A) and the 10–30% typical of somatic cells in various species ((14, 100), Figure 2A). These data underscore the exceptional rarity of inter-chromosomal interactions in lampbrush-stage oocytes (Figure 2A, Supplementary Table S1).

**Figure 2.**
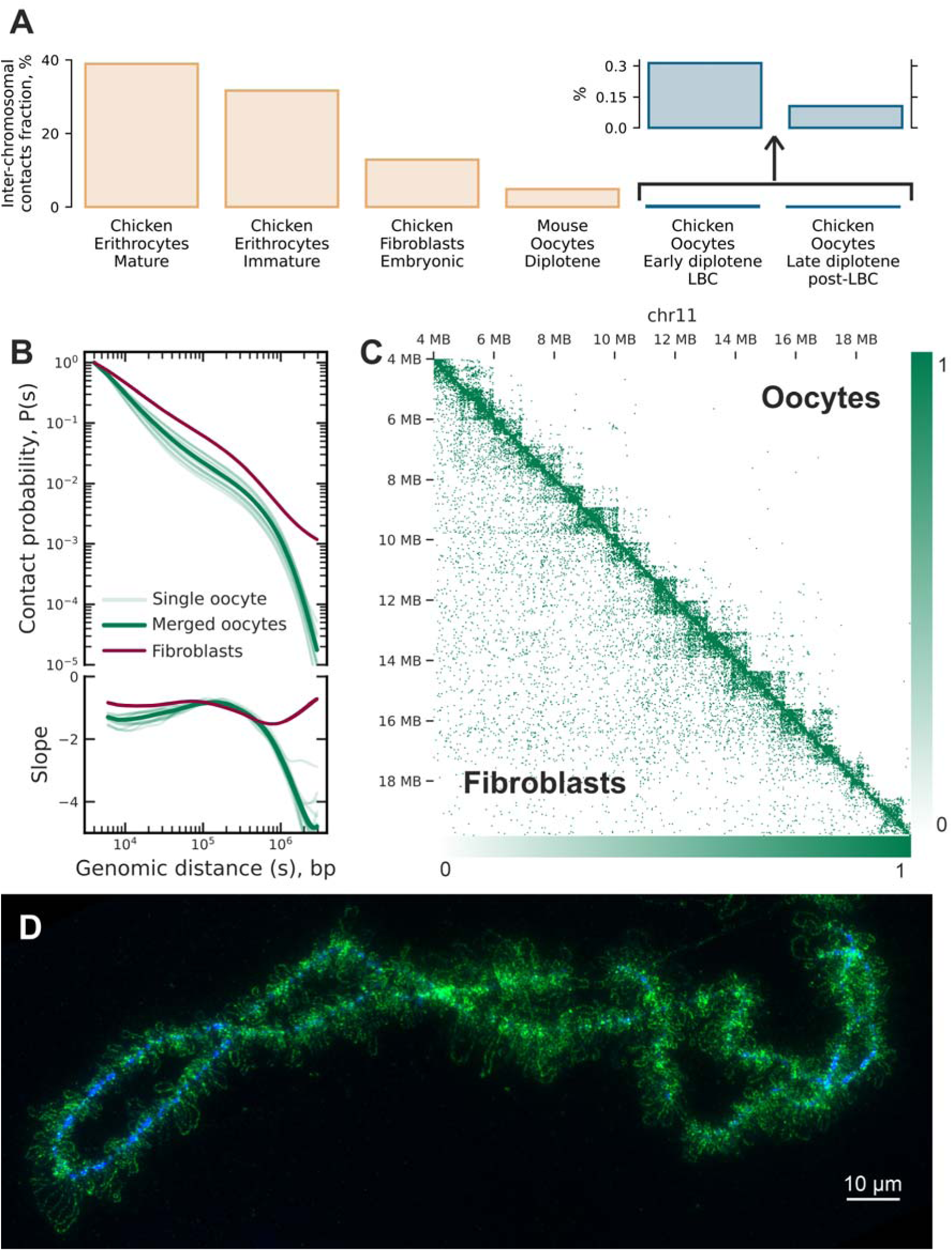
Chromatin organization of lampbrush chromosomes at different genomic distances. A) Fraction of inter-chromosomal contacts across different cell types and species (see Methods). Chicken oocytes at the lampbrush chromosome stage exhibit a markedly reduced proportion of inter-chromosomal contacts, consistent with cytological observations of spatial chromatin organisation. B) Contact probability versus the genomic distance for single and merged oocyte data (lampbrush chromosome stage, LBC) and chicken embryonic fibroblasts (see Methods). The genomic distances are cut to display only representative contacts of oocytes. See extended version in Supplementary Figure S3A. C) Hi-C contact maps comparing pseudo-bulk chicken oocytes and bulk fibroblast cells for chromosome 11. The fibroblast map was downsampled to match oocyte data (both maps sum to 19,946 contacts within the depicted region). Each bin represents 32 Kb. See extended version in Supplementary Figure S3B. D) Chromomere-loop structure of isolated chicken lampbrush chromosome 1. Active RNA polymerase II is labeled with antibodies (green), and chromatin is counterstained with DAPI (blue).

Intra-chromosomal (cis) interactions in lampbrush chromosomes exhibit an abrupt drop in contact frequency beyond ∼1 Mb (Figure 2 B-C, and Supplementary Figure S3A), consistent with the elongated, rope-like morphology of these chromosomes (Figure 2D). The depletion of both long-range and trans-chromosomal interactions in LBC Hi-C maps results in the absence of the plaid-pattern of contacts typically observed in interphase nucleus Hi-C data, which is associated with chromatin compartmentalization ((101), Supplementary Figure S3B and Supplementary Figure S4; see Methods).

### Chromatin of lampbrush chromosomes forms domains with stable boundaries shared across individual oocytes

In contrast to the absence of a plaid-pattern, chromatin domains are clearly visible on LBC Hi-C maps at scales of 0.5–2 Mb (Figure 2C). To understand whether the domain organization observed in chicken LBCs represents a general feature of meiotic chromatin or a unique, potentially LBC-specific mechanism, we compared our results with mouse diplotene oocytes. Available Hi-C maps for individual nuclei of mouse diplotene oocytes, obtained by a similar approach (42), provide an opportunity to compare contact chromatin maps in organisms with different types of oogenesis.

In mouse diplotene oocytes at the non-surrounded nucleolus (NSN) stage, chromatin interaction analysis reveals TADs with highly variable boundary locations across individual cells (42). In contrast, single-nucleus Hi-C maps from chicken lampbrush-stage oocytes demonstrate striking consistency in domain boundary positions (Figure 3A-C), with a mean normilized F1 score (see Methods) of 2.3 compared to 1.3 for mouse NSN oocytes (Figure 3A). This stability in domain boundaries across chicken LBCs suggests fundamental differences in the mechanisms underlying chromatin domain formation between these two systems.

**Figure 3.**
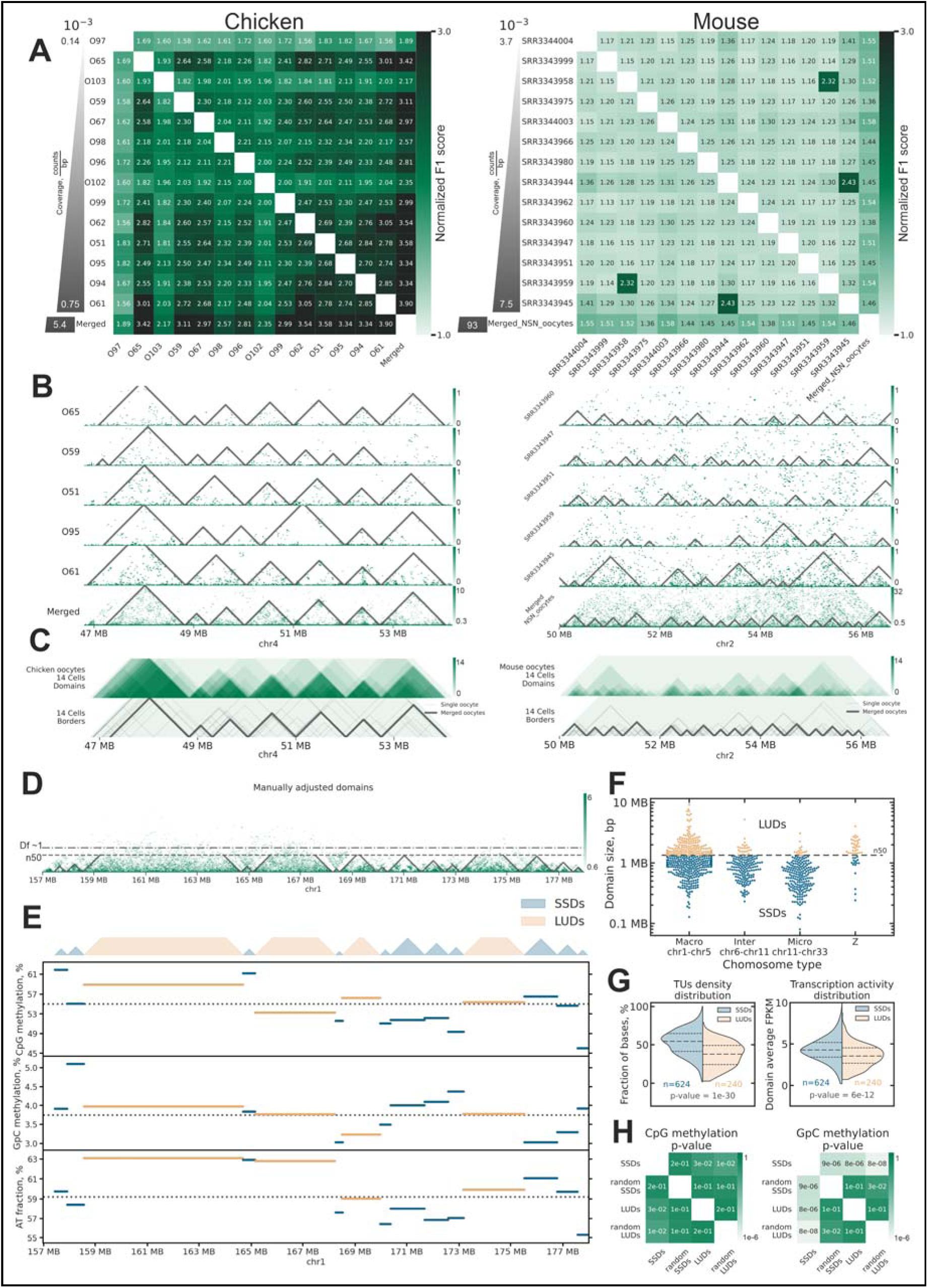
Chromatin domains in chicken lampbrush chromosomes. A) Pairwise normalized F1 scores for domain boundary concordance across individual oocytes. Left panel: chicken lampbrush-stage oocytes; right panel: top 14 mouse NSN oocytes by coverage (42). Oocytes are sorted by the number of Hi-C contacts per bp. Higher normalized F1 scores indicate greater concordance in boundary positions. P-value between chicken and mouse normalized F1 score distributions < 10^(−30). B) Domain annotations for individual cells and merged Hi-C data for chicken (left) and mouse (right) oocytes. Note the higher concordance of domain boundaries in chicken oocytes compared to mouse oocytes. Resolution 20 kb. C) Aggregated Hi-C contacts and domain annotations for 14 chicken oocytes (left) and top 14 mouse oocytes by coverage (right). D) Merged Hi-C data from chicken oocytes with domain annotations overlaid. Domains were manually refined after automatic annotation (see Methods). The dashed line indicates the N50 domain size (∼1.3 Mb). The dash-and-dot line indicates genomic distance where a sharp drop in the frequency of contacts was observed (distance with intra-domain fractal dimension 1, stick-like polymer, see Supplementary Figure S7A). E) Each horizontal segment represents a single chromatin domain, with the Y-position of the line indicating the domain’s average characteristic (CpG methylation, GpC methylation, or AT fraction, see Methods). The dashed horizontal line represents the genome-wide average for each characteristic. This visualization highlights the distinct properties of Large Unstructured Domains (LUDs) and Small Squared Domains (SSDs). The genome-wide CpG and GpC methylation distributions can be found in Supplementary Figure S5C-D. The genome-wide domain AT fraction distribution can be found in Supplementary Figure S7B-C. F) Distribution of LUDs and SSDs lengths on chicken chromosomes. G) The transcription units (TUs) density and activity distribution show that the SSDs contain significantly more TUs, and TUs with higher activity, than LUDs. H) CpG and GpC methylation levels statistically compared between SSDs and LUDs (see Methods). As a baseline, domains were randomly redistributed across the genome while preserving the number of intervals and their length distributions (indicated as ‘random’). SSDs exhibit significantly higher GpC methylation levels, reflecting increased chromatin accessibility. The genome-wide CpG and GpC methylation distributions can be found in Supplementary Figure S5C-D. Panels F–H collectively demonstrate that Large Unstructured Domains (LUDs) are enriched on macrochromosomes and the Z chromosome and are characterized by low gene density and transcriptional quiescence, whereas Small Structured Domains (SSDs) are gene-dense, transcriptionally active, and broadly distributed across the genome.

Contact domains in LBCs can be classified into two types based on their length (Figure 3D,E): 1) <u>L</u>arge <u>U</u>nstructured <u>D</u>omains (LUDs), defined as domains longer than 1.3 Mb (N50 of domains length, see Methods) and 2) <u>S</u>mall <u>S</u>quared <u>D</u>omains (SSDs), shorter than 1.3 Mb. Visually, LUDs have more dispersed contacts that are connecting loci up to ∼2 Mb from each other (Supplementary Figure S5A). Almost all LUDs are located on macro (chr1-chr5) and Z chromosomes (Figure 3F).

To characterize chromatin states in LUDs and SSDs, we performed a single cell NOMe-seq experiment. NOMe-seq measures open chromatin via deposition of exogenous GpC methylation, enabling simultaneous analysis of chromatin accessibility and CpG methylation levels in individual nuclei. We augmented NOME-seq data with CpG methylation from (80) (see Methods). We also quantified transcription levels, reanalyzing RNA-seq data from the previous chicken oocyte nucleus transcriptome study (57) and supplementing it with additional replicas generated in this work. Analysis of these data shows that LUDs have lower median gene density (Figure 3G, Supplementary Figure S5B) and lower normalized expression levels than SSDs (Figure 3G). We found that SSDs have significantly higher levels of chromatin openness (measured in NOME-seq experiment via GpC methylation), indicating that shorter domains contain more accessible chromatin (Supplementary Figure S5C), while there is almost no difference in CpG methylation levels between SSDs and LUDs (Figure 3H, Supplementary Figure S5D), as well as only a weak correlation between promoter methylation and gene expression levels (Supplementary Figure S6). We note that the length threshold separating LUDs and SSDs is somewhat arbitrary, as the properties of both gradually change with the domain length (Supplementary Figure 5E).

Our findings indicate that Large Unstructured Domains correspond to gene-poor, transcriptionally inactive chromatin regions predominantly found on macrochromosomes and the Z chromosome. In contrast, Small Squared Domains are associated with gene-rich, transcriptionally active chromatin regions distributed across all chromosomes.

Based on this characteristic, we propose that SSDs and LUDs in single-nucleus Hi-C maps correspond to chromomeres in LBCs. Additionally, given their properties, we speculate that LUDs represent more compact, DAPI-positive chromomeres, while SSDs correspond to less compact, DAPI-negative chromomeres. Supporting this hypothesis, statistical analysis of the AT fraction in domains—reflecting the genomic proportion susceptible to DAPI staining—shows a weak but significant correlation between AT fraction and domain length (Spearman’s R = 0.55, compared to R = 0.28 for randomly shuffled domains, Supplementary Figure S7B-C).

To further validate the correlation between domain types and DAPI staining in chromomeres, we analyzed sequence markers with known positions in cytological maps of chicken LBCs, primarily derived from DNA- and RNA-FISH experiments using specific BAC clones (Figure 4A-B). The distribution of LUDs and SSDs correlated with the arrangement of DAPI-positive and DAPI-negative chromomeres at certain regions of chromosomes. For instance, we observed multiple LUDs between markers W030P07 and W030B21, coinciding with a region enriched in DAPI-positive chromomeres (Figure 4A-B). However, the number of cytologically visible chromomeres and contact domains on Hi-C maps does not always match exactly, probably due to the limited resolution of microscopy analysis and/or errors in domain calls. Together with sparse availability of sequence markers distinguishing chromomeres and resolution of cytological chromomere-loop maps this limited a genome-wide analysis of the relationship between SSDs, LUDs, and chromomeres.

**Figure 4.**
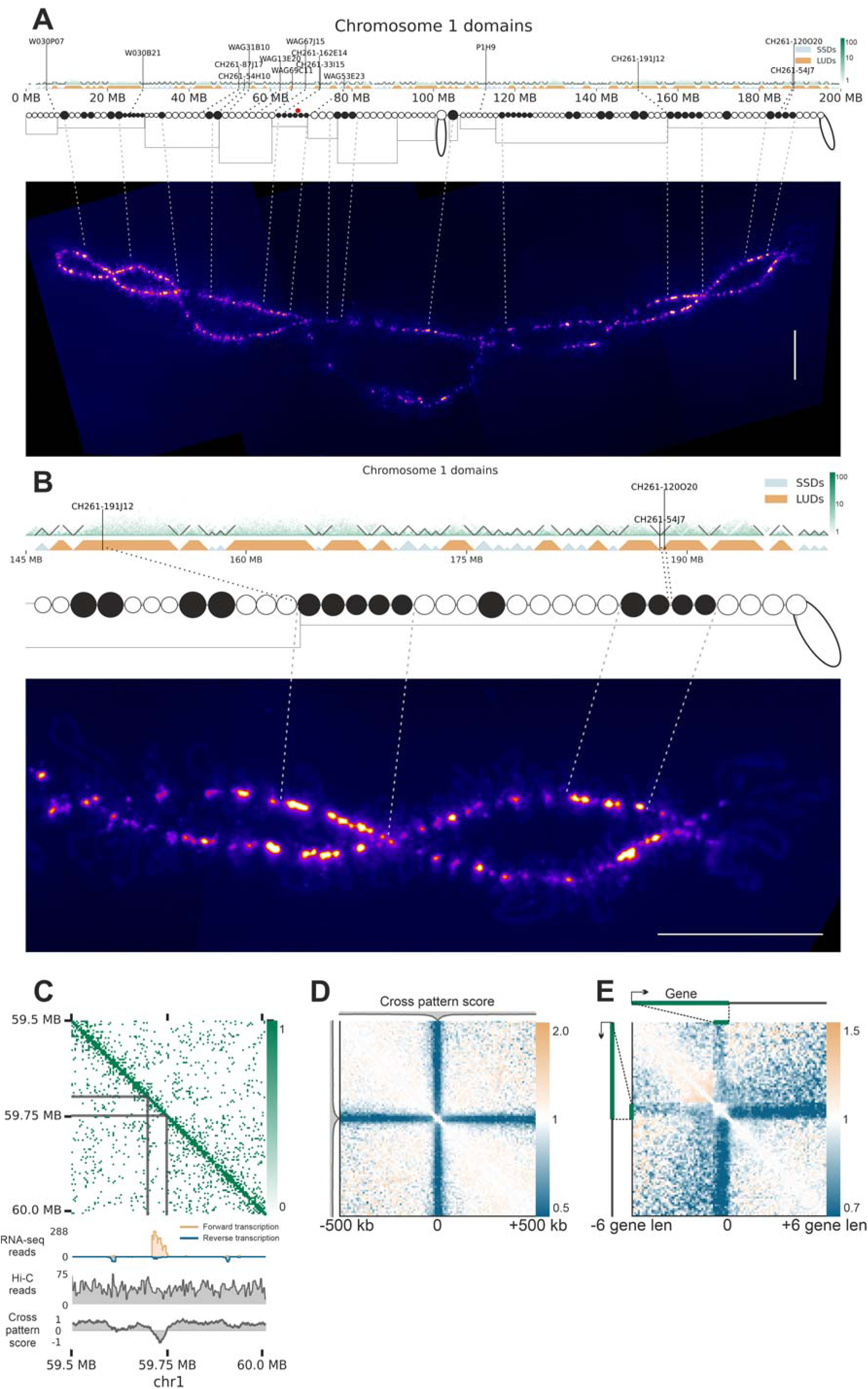
Concordance between microscopically visible chromatin structures and Hi-C map patterns along chicken lampbrush chromosomes. A) Comparison of Hi-C contact domains with chromatin domains identified through microscopy visualization of lampbrush chromosomes. From top to bottom: Hi-C map for chicken lampbrush

### Hi-C patterns of transcription units and compact chromatin domains align with transcription loops and chromomeres identified by microscopy analysis

Transcription loops are a hallmark of lampbrush chromosomes. These loops represent stretched fibers of the actively transcribed chromatin emanating from the LBC axis. Given their spatial separation from each other and the LBC axis, we hypothesize that loci corresponding to transcription loops lack interactions with the rest of the genome.

Visual inspection of Hi-C maps reveals loci with a distinct "cross" pattern, where interactions are limited to nearest neighbors (Figures 4C). These loci compensate reduced long-range interactions by increasing frequency of proximal interactions, thus maintaining consistent coverage levels (Figure 4C). To systematically identify and visualize these features, we introduce a quantitative Cross Pattern Score that captures the relative enrichment of short- versus long-range contacts at each locus (see Methods). Although we observe some variability on “cross” pattern manifestation across loci, aggregated Hi-C maps centered on the regions with low Cross Pattern Score (predominant short-range contacts) display characteristic “cross” pattern (Figure 4D), demonstrating that “cross” features are reproducible genome-wide.

"Cross" patterns frequently occur within chromatin domains and almost always coincide with actively transcribed genes. Supporting this observation, aggregated Hi-C maps centered on long, highly expressed genes (top 50% by length and TPM) show a pronounced "cross" pattern (Figure 4E). These findings suggest that "cross" patterns in Hi-C maps represent the contact signatures of transcription loops in LBCs.

To align lampbrush chromatin contact patterns identified by Hi-C analysis with chromatin structures observable via microscopy, we conducted FISH mapping using probes specific to chromatin domain boundaries, internal domain regions, and loci forming “cross” patterns. A total of 29 BAC clones were hybridized in various combinations on lampbrush chromosomes 1, 4, and 13 (Supplementary Table S3). The large size of lampbrush chromosomes enables detailed microscopy analysis, allowing us to distinguish chromomeres and transcription loops, as well as determine the orientation of transcription units.

The locus analyzed on chromosome 1 contains six Hi-C domains and was covered by 14 BAC clone based probes. Regions 4, 6, 7, 9, and 12 coincide with "cross" patterns with reduced long-range interactions observed in Hi-C maps and correspond to actively transcribed genes (Figure 5A; Supplementary Figure S8). Note that "cross" patterns demonstrate locus-specific heterogeneity in their structure depending on the length and orientation of transcription units. Based on the annotated transcription units’ location and orientation, we anticipated transcription loops with forward-oriented transcription for regions 4, 9, and 12, and reverse-oriented transcription for regions 6 and 7. FISH mapping of LBC1 using BAC clone-based probes with a DNA+RNA hybridization protocol confirmed these predictions. Long lateral loops with a prominent RNP matrix were observed for BACs 4, 9, and 12, with transcription co-directional to the chromosome (Figure 5B, subpanels 2a, 3a-b; Supplementary Figure S8 subpanel c).

**Figure 5.**
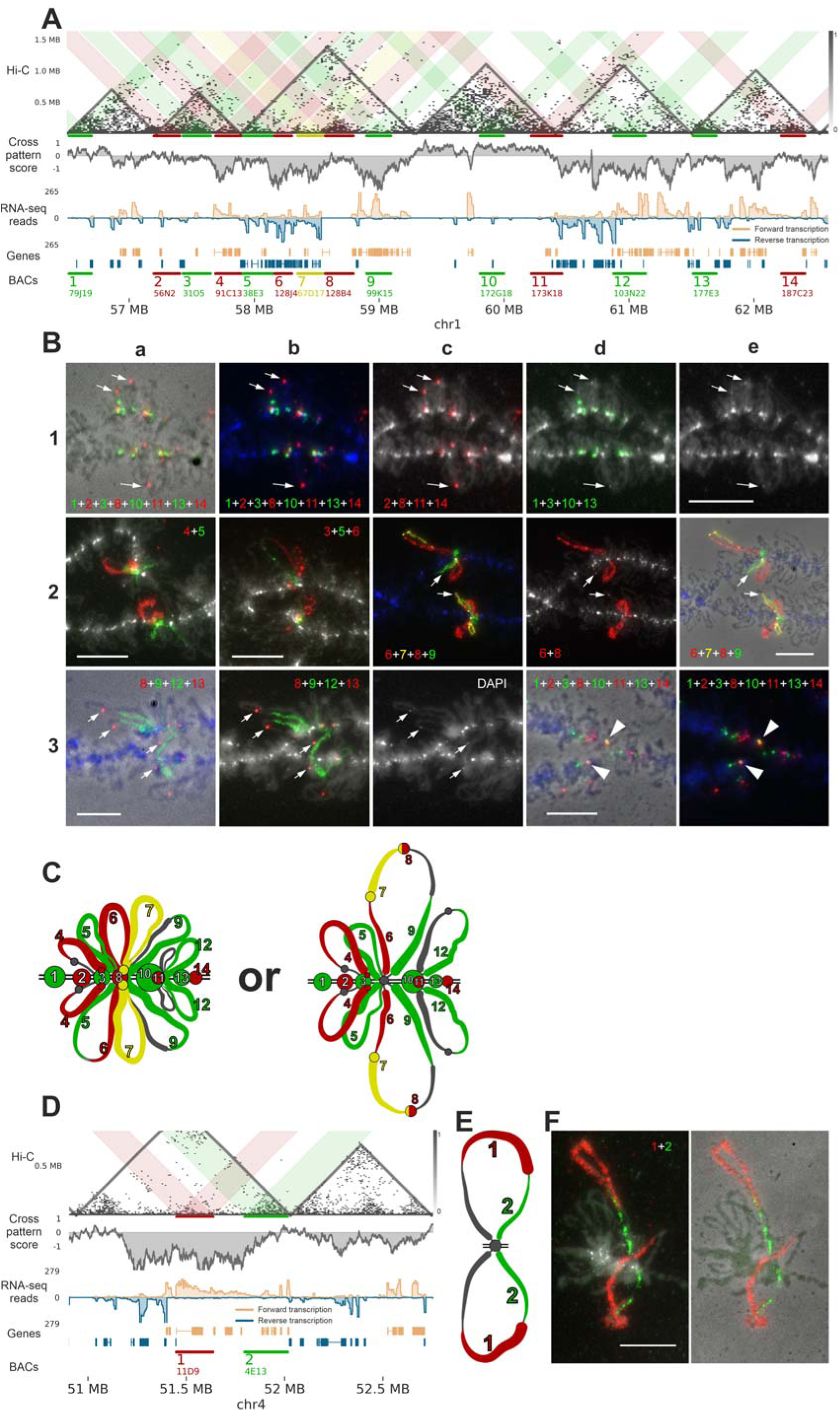
Comparison of Hi-C map patterns with microscopically identified chromatin domains and loops in lampbrush chromosomes 1 and 4. A) Hi-C map and associated data for a fragment of chromosome 1. From top to bottom: combined Hi-C map of the oocyte nuclei, with colored highlights indicating interactions in regions covered by BAC

Similarly, genomic regions 6, 7, and 9, covered by three different BACs, correspond to transcription units forming one or two pairs of lateral loops (Figure 5B, subpanels 2c–e; Supplementary Figure S8 subpanels b, c). BAC 5 doesn’t correspond to a region with an obvious “cross” pattern according to the Hi-C data, however it covers the transcriptionally active region (in reverse direction), thus it can form a transcription loop (Figure 5B subpanels 2a-b). In our FISH experiments, BAC 5 marks a reverse oriented transcription loop anchored in the chromomere (Figure 5A,C). Another combinations of DNA-probes mapped on the chromosome 1 are represented on Supplementary Figure S8. Additionally, RNA-FISH experiments without chromosomal DNA denaturation for several transcribed regions confirmed that BAC clone-based probes hybridize with nascent transcripts on the lateral loops (Supplementary Figure S8 subpanel c). These findings confirm that "cross" patterns observed on Hi-C maps of avian LBCs correspond to the microscopically visible lampbrush lateral loops containing actively transcribed genes.

The intra-domain regions 1, 2, 3, 8, 10, and 14 correspond to five of the six contact domains identified on LBC Hi-C maps, with BACs 2 and 3 covering the same domain (Figure 5 A). Consequently, we anticipated observing five insulated chromomeres along the chromosome axis in microscopy images, one of which would be associated with two BACs. FISH experiments confirmed this, with BAC clone-based probes mapping to five distinct chromomeres (or chromatin knots in case of BAC8, see below) (Figure 5B, subpanels 1a–e, 3d-e; Figure 5C; Supplementary Figure S8 subpanels e-g, f). The chromomeres covered by BACs 2 and 3 are closely spaced but remain distinguishable (Figure 5B, subpanels 1c–d; Supplementary Figure S8 subpanel a).

Interestingly, the compact chromatin domain covered by BAC 8, which lacks transcription, exhibits variable positioning. This chromatin domain can either anchor directly to the chromosome axis (Figure 5B panels 2c-e; Supplementary Figure S8 subpanel d) or remain detached from it (indicated by white arrows in Figure 5B panels 1a-e, 2c-e and 3a-c), forming a compact "knot" on a lateral loop emanating from the chromosomal axis. Four signals for BAC8 correspond to four chromatin knots on each of the sister chromatids (two sister chromatids on each half-bivalent). These chromatin knots, located within transcriptionally active lateral loops, have been previously described as small genomic regions devoid of transcribed genes (Figure 6 in (57)).

**Figure 6.**
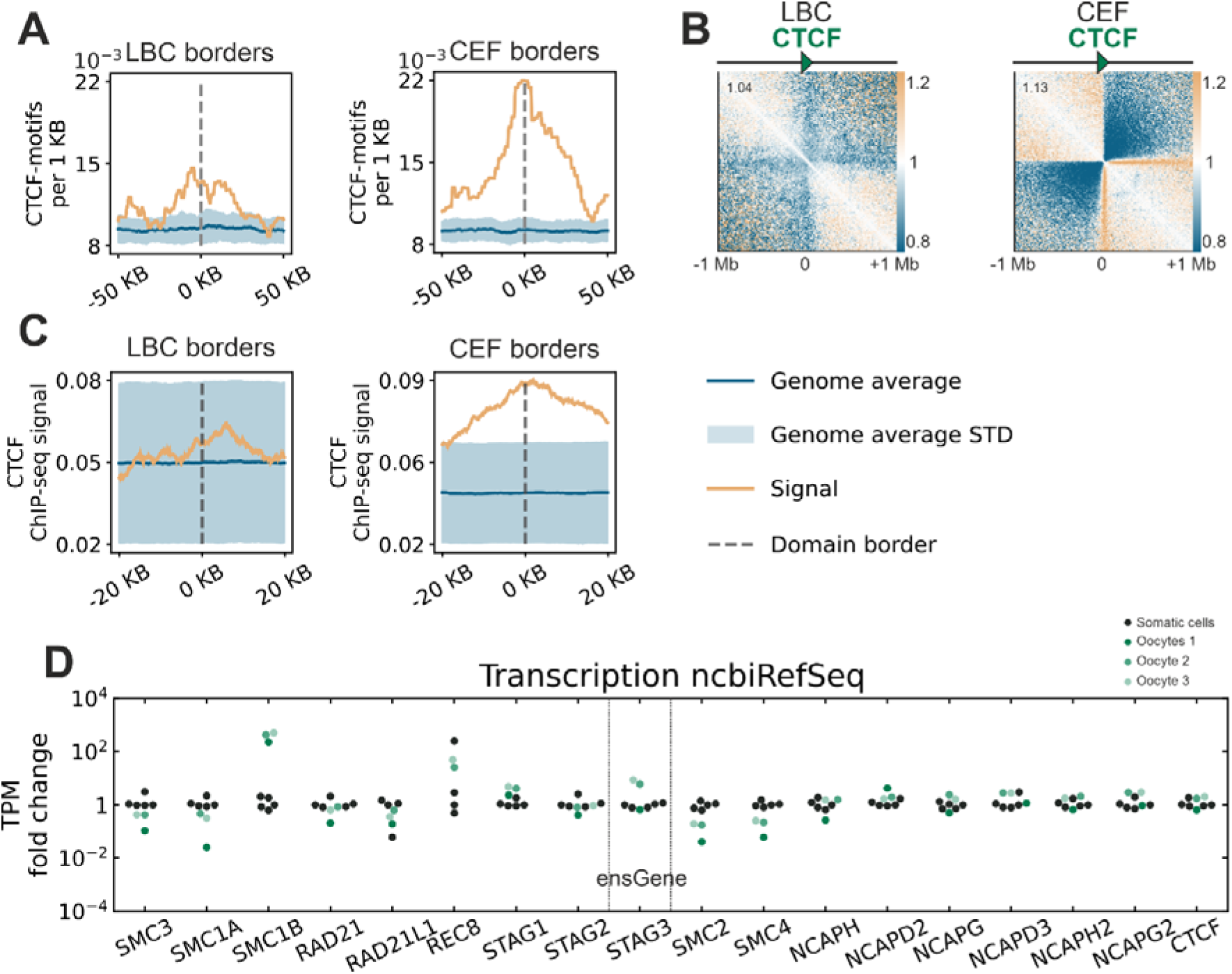
The mechanism of chromatin contact domain formation in lampbrush chromosomes is CTCF independent, despite transcriptional activity of the *CTCF* gene. A) Frequency distribution of CTCF-motifs at domain boundaries in lampbrush chromosome (LBC) and chicken embryonic fibroblast (CEF) maps. Both LBC and CEF boundaries show enrichment relative to the genome average, with peak enrichment factors of ∼1.4 and ∼2.3, respectively. B) Aggregated Hi-C contact maps centered on CTCF-motifs with uniform orientation. While strong insulation is evident around CTCF-motifs in CEF, only weak insulation is observed in LBC contact maps. Resolution: 10Jkb, fibroblast data subsampled to match number of Hi-C contacts in merged LBC. С) Input-normalized CTCF ChIP-seq signal from chicken liver (SRR4068194, see Methods) at oocyte contact domain boundaries and fibroblast contact domain boundaries. The absence of CTCF enrichment across LBC domain boundaries contrasts with the strong signal observed in fibroblast boundaries. D) Comparison of transcript levels for cohesin subunits, condensin subunits, and *CTCF* (TPM fold-change relative to the somatic cell median). Oocytes exhibit reduced transcript levels for *SMC3*, *SMC1A, SMC2,* and *SMC4* genes, higher transcript levels for *SMC1B* and *REC8* genes, and similar *CTCF* transcript levels compared to fibroblasts. Gene annotations were sourced from ncbiRefSeq, except for *STAG3*, which used ensGene annotations (see Supplementary Table S4). Green dots represent transcript levels from three chicken oocyte RNA-seq datasets, and black dots represent five replicates of chicken embryonic fibroblasts (84). Oocytes 1: Total RNA stranded RNA-seq data (57). Oocytes 2 and 3: Poly(A)-enriched stranded RNA-seq replicates generated in this study.

Together, these results demonstrate that: (i) chromatin contact domains identified through Hi-C analysis correspond to insulated chromomeres in lampbrush chromosomes, and (ii) small regions between actively transcribed genes can form microdomains (knots) that are optionally tethered to the chromosomal axis.

Region 11 exhibits partial transcriptional activity and is positioned near the boundaries of chromatin domains (Figure 5A). Based on this, we anticipated that BAC 11 would cover a portion of chromatin spanning two adjacent chromomeres, one of which is covered by BAC 10. FISH mapping confirmed this prediction, showing partial colocalization of BACs 10 and 11 within a prominent DAPI-positive chromomere (Figure 5B, subpanels 1a–d and 3d-e; Supplementary Figure S8 subpanel a, f). The DAPI-positive chromomere partially covered by BACs 10 and 11 corresponds to the most prominent contact domain within analysed genomic locus.

Similarly, BAC 13, located near the boundary of contact domain, produced a signal spatially close to that of BAC 14, which maps to the same domain. While the signals from BACs 13 and 14 are closely positioned, they remain distinct and do not intermix (Supplementary Figure S8 subpanel e).

These results confirm that boundaries between domains identified by Hi-C are concordant with LBC chromomere boundaries (see additional evidence supporting this statement in the Supplementary Note 2).

On lampbrush chromosome 4, we mapped nine BAC clone-based probes across two regions of interest. In the region A (51–52 Mb), Hi-C data revealed the formation of a contact domain with interactions at the top of the domain triangle and a wide “cross” pattern within domain, indicative of chromatin loop formation (Figure 5D). Nuclear RNA-seq data showed that several genes within region A are transcribed in varying orientations. FISH mapping confirmed that this genomic region corresponds to a long transcription loop with multiple RNP matrix gradients (Figure 5E,F), consistent with the transcriptional activity detected. Detailed results for region B are provided in Supplementary Note 2 and Supplementary Figure S9.

We integrated the FISH data to reconstruct the chromatin organization of the analyzed region on chromosome 1 (Figure 5C), accounting for two alternative states of chromatin knots discussed above. Schematic drawing represents an approximation of the FISH mapping data for different combinations of DNA probes to lampbrush chromatin domains seen in at least 4 micrographs (half-bivalent is shown). This reconstruction demonstrates a strong concordance between microscopically visible chromatin structures—such as chromomeres and transcription loops—and the contact domains and “cross” patterns observed in Hi-C maps. The reconstruction of the region A on chromosome 4 also confirmed the correspondence between the transcription loops and the “cross” patterns (Figure 5D,E).

Additionally, similar analyses of two other regions on chromosomes 4 and 13, which included BACs mapped to Hi-C domain boundaries, intra-domain fragments, and “cross” patterns, yielded consistent results (Supplementary Figure S9; Supplementary Note 2). These findings validate the Hi-C maps of chicken oocyte nuclei and confirm the interpretation of Hi-C patterns as representations of lampbrush chromosome chromomeres and lateral loops.

### Chromatin domains in lampbrush chromosomes are formed by a CTCF-independent mechanism

In the majority of vertebrate cell types, including chicken fibroblasts (15), TAD boundaries are demarcated by CTCF binding sites (1, 101). The moderate conservation of CTCF binding site distribution across cell types, combined with a strong correlation between CTCF binding and TAD boundaries, allows CTCF enrichment to be detected even when comparing Hi-C maps to CTCF ChIP-seq data from different cell types (15) and cell-type agnostic motif enrichment.

We show that LBC domain boundaries display almost no enrichment of CTCF motifs (Figure 6A). Concordantly, almost no insulation was detected when LBC Hi-C contacts were aggregated across CTCF motifs (Figure 6B). To provide a ground for comparison, we performed the same analysis for chicken fibroblasts data downsampled to the same sequencing depth. In fibroblasts, we detect strong insulation across CTCF motifs and enrichment of motifs at TAD boundaries (Figure 6A-B). We next replicated this analysis using CTCF ChIP-seq signal from chicken liver, which confirms the absence of CTCF-mediated insulation in LBC (Figure 6C). In contrast, strong enrichment of chicken liver CTCF ChIP-seq signal was observed at fibroblast TAD boundaries (FigureL6C), validating the approach and reinforcing the conclusion that chromatin contact domains in lampbrush chromosomes are formed independently of CTCF binding.

To understand which chromatin architecture proteins can contribute to the formation of contact domain boundaries in oocytes, we explored the chicken oocyte nucleus transcriptome data (see Methods). Despite the absence of CTCF binding enrichment in the domain boundaries, in oocytes *CTCF* gene has the comparable transcript levels with fibroblasts. Genes for the other architectural proteins, such as *RAD21, STAG1*, *STAG2* (cohesin) and *NCAP-H, NCAP-D2, NCAP-G, NCAP-D3, NCAP-H2, NCAP-G2* (condensin) subunits also show comparable transcript levels in fibroblasts and oocytes (Figure 6D, Supplementary Table S4). Consistent with this, the presence of SMC1A, SMC3, Rad21, STAG1, and STAG2 on avian LBC axes in a punctate pattern has previously been shown by immunostaining, with an enrichment of cohesin subunits in the interchromomere regions and in chromomere cores (88). Among the condensin subunits, XCAPD2 was found to accumulate in chromomeres of Xenopus LBCs (103). However, the main cohesin subunits (SMC3, SMC1A) and the main condensin subunits (SMC2, SMC4) have about ten times weaker transcript levels in the LBC stage oocyte (see Figure 6D, Supplementary Table S4). In contrast, the meiosis specific cohesin subunit SMC1B in oocytes has an almost thousand times higher level of transcripts than in fibroblasts (Figure 6D, Supplementary Table S4). Thus, although components of extrusion complexes are present at LBC-stage, their activity might be modulated by quantity of proteins and usage of cell type specific subunits, potentially explaining why extrusion is not blocked by CTCF binding sites as in typical somatic cells. Moreover, the synaptonemal complex protein SYCP3 has been found at the axes of avian LBCs (88), which may affect the formation of structural chromatin loops.

### Transcriptional activity and orientation of transcription units define lampbrush chromosome chromatin domains

To investigate the genomic features underlying the formation of LBC domains, we compared domain borders with position-dependent genomic elements. CpG methylation levels and chromatin accessibility at domain boundaries were similar to the genomic average (Supplementary Figure S10A, see Methods). However, boundaries showed enriched GC-content relative to the background, suggesting a correlation with the presence of genes.

Aggregation of contacts around exons, introns, and repetitive elements revealed no insulatory potential for these features, arguing against their role in forming domain boundaries at the lampbrush stage (Supplementary Figure S10B). In contrast, we observed a significant depletion of contacts across 3′-UTRs and, to a lesser extent, across 5′-UTRs (Figure 7A). This suggests that the distribution and orientation of transcription units may play a key role in LBC domain formation.

**Figure 7.**
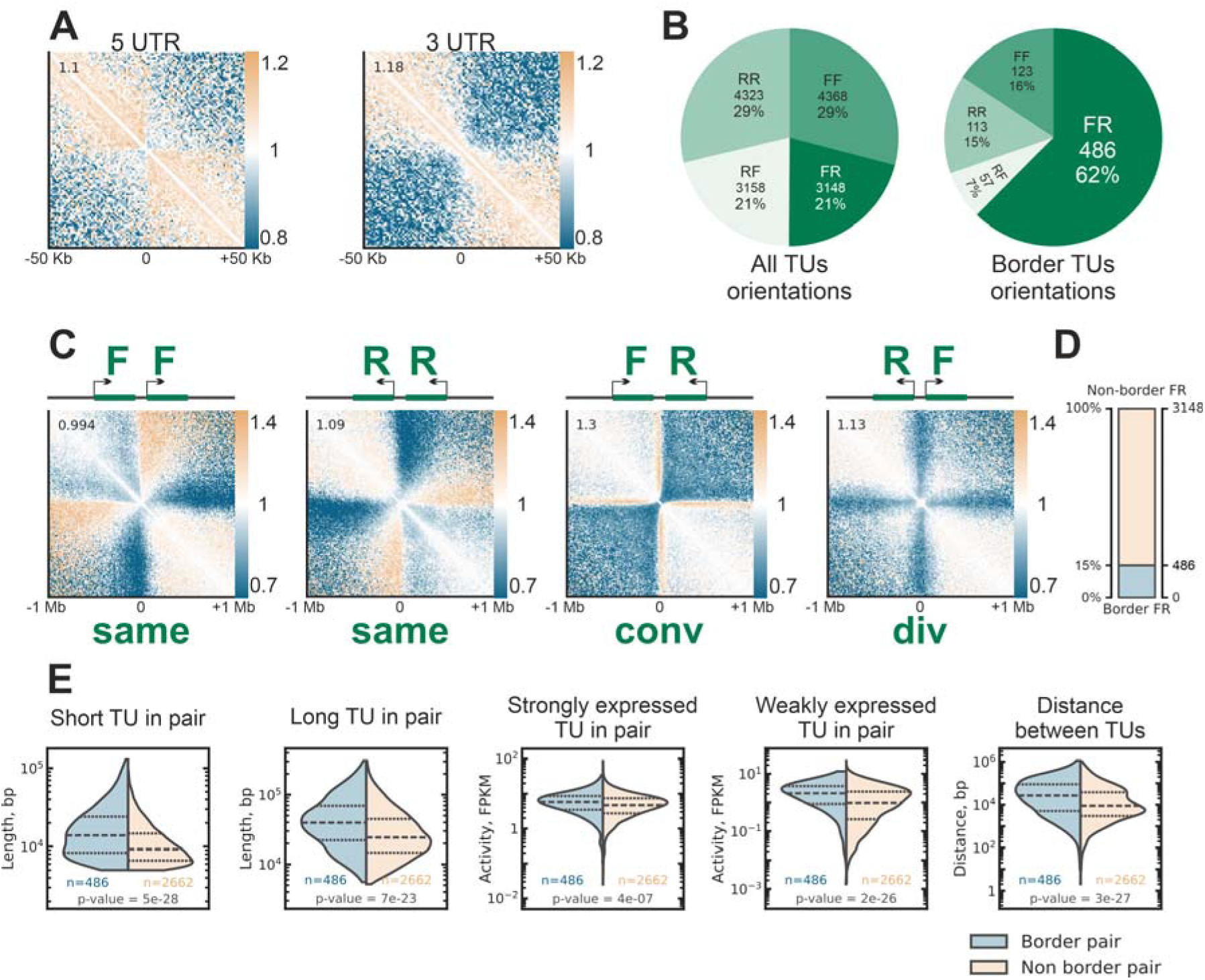
Aggregated 3D chromatin interaction profiles reveal genomic features underlying lampbrush chromosome chromatin domains. A) Aggregated contact maps around 5′ and 3′ UTRs show depletion of interactions across these regions. Insulation scores are displayed in the upper-left corner of each aggregated map triangle (see Methods). Resolution: 10 Kb B) Pie charts displaying transcription unit (TU) orientations for all TU pairs and those forming domain boundaries. A significant enrichment of Forward-Reverse (FR, convergent) orientation is observed at domain boundaries. C) Aggregated contact maps centered between TUs in different orientations: Forward-Forward (FF, co-directional), Reverse-Reverse (RR, co-directional), Forward-Reverse (FR, convergent) and Reverse-Forward (RF, divergent). Insulation score, shown in the upper-left corner of each map, is highest for convergent (FR) TU pairs. RF pairs typically form a "cross" pattern, while FR, FF, and RR pairs display a stripe-like enrichment of contacts between TUs and upstream loci, with FF and RR stripes widening with distance from the diagonal. Insulation scores are displayed in the upper-left corner of each aggregated map triangle (see Methods). Resolution: 10 Kb D) Statistics of FR-oriented TU pairs. Despite their prevalence at domain boundaries, only ∼15-20% of FR-oriented TU pairs form boundaries. E) Comparison of TU features between border-forming and non-border FR-oriented TUs. Border-forming TUs are, on average, longer, more highly expressed, and separated by greater intergenic distances. However, the length of the shorter TU and the activity of the weaker TU in the pair show higher difference between border-forming and non-border FR-oriented TUs than the features of the longer or more strongly expressed TU. p-values obtained using Mann–Whitney U test.

To explore the potential link between transcription and LBC domain boundaries, we aligned RNA-seq data from poly(A)-enriched and total stranded RNA libraries ((57), see Methods) with Hi-C maps and visually inspected these profiles. This analysis highlighted transcriptional activity as a critical determinant of LBC chromatin domain organization.

Visual comparison of transcriptomic data with chromatin domain boundaries revealed multiple transcription units absent from the current gene annotations in (68) (Supplementary Figure S11). This discrepancy reflects the dependence of chicken genome annotation on cell type and database, particularly for non-coding RNAs (104). To address this, we performed *de novo* transcriptome assembly using oocyte RNA-seq data, generating a refined set of transcription units (TUs) active during the LBC stage (see Methods).

Analysis of this updated TU set confirmed a strong correlation between chromatin insulation and TU orientation (Figure 7B-C). Genomic regions flanked by TUs in a convergent (Forward-Reverse, FR) orientation were significantly enriched at chromatin domain boundaries (Figure 7B). Furthermore, FR-oriented TU pairs exhibited the highest insulation scores compared to other TU pair orientations (Figure 7C).

Manual validation of domain boundaries revealed that nearly half of non-FR TU pairs initially identified at domain boundaries were artifacts of automatic domain annotation or transcriptome assembly (26 of 47 randomly selected for inspection boundaries; Supplementary Figure S17). Overall, convergent transcription accounts for at least 79% of domain boundaries, with 62% supported by automatic annotation and an additional 17% identified through manual correction.

Given the central role of transcription in defining LBC chromatin domains, we designate the domains detected in Hi-C maps as **T**ranscription-**D**ependent **D**omains (**TDDs**).

Although the majority of TDD boundaries are formed by TU pairs in a convergent (FR) orientation, only ∼15% (∼20% after manual curation) of FR-oriented TU pairs create boundaries, while the remainder are located within domains (Figure 7D). To investigate the properties of TU pairs associated with boundary formation, we compared transcript features of "border" versus "non-border" TU pairs in a convergent orientation (Figure 7E). No single feature was sufficient to distinguish "border" TU pairs from "non-border" pairs. However, we observed that TU pairs forming boundaries tend to have several distinguishing characteristics: they are, on average, longer, separated by a larger intergenic interval, and exhibit higher activity of the weaker TU of the pair. The activity of the stronger TU in the pair shows minimal difference between boundary-forming and non-boundary-forming pairs. These observations suggest that TDD boundary formation is governed by a complex interaction of TU features rather than a single determinant.

### Fusion of transcription-dependent chromatin domains during oocyte growth and decreased transcriptional activity

Growth of chicken oocyte coincides with reduced transcriptional activity and gradual contraction of both chromosomes and lateral loops (105). To study the dynamics of this process, we isolated nuclei from the late-diplotene stage oocytes (hereinafter referred to as post-LBC) and performed a single-nucleus Hi-C experiment (Figure 1A, Supplementary Table S1,2). Overall, Hi-C patterns detected in the mid-diplotene (LBC) stage persist at the post-LBC stage. However, there were quantitative differences. Concordant with cytological evidence of LBC contraction, we observe an increase of long-range interactions at the post-LBC stage (Figure 8A,B). Moreover, the significant decrease of the loop sizes of the chicken post-LBC chromosomes (105) agrees well with the almost complete absence of the “cross” patterns (Figure 8C). This observation confirms our suggestions about the critical role of the TU loop repulsion in the “cross” pattern formation.

**Figure 8.**
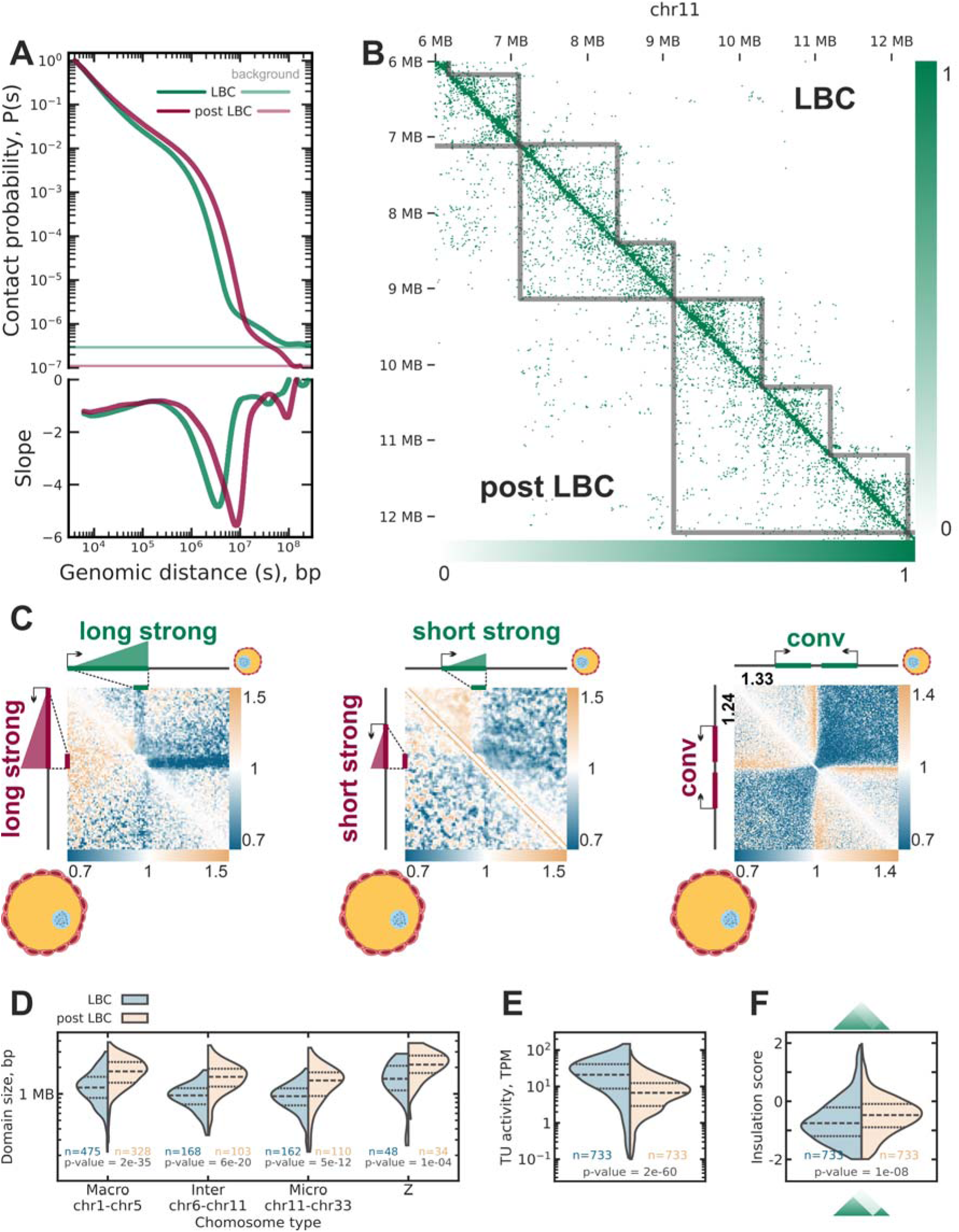
Comparison of oocyte chromatin at the lampbrush and post-lampbrush stages. A) Contact probability as a function of genomic distance for lampbrush (LBC) and post-lampbrush chromosome (post-LBC) stages. Post-LBC chromatin exhibits an increase in distant contacts, consistent with microscopy observations (105). B) Pseudo-bulk Hi-C interaction maps for LBC (8719 contacts) and post-LBC (4949 contacts) chromatin. Grey lines indicate automatically adjusted domain borders for each stage. Neighboring domains fuse in the post-LBC stage, as demonstrated in the shown examples. C) Aggregated observed-over-expected Hi-C contact maps comparing lampbrush chromosome (LBC) stage (upper triangle) and post-LBC stage (lower triangle) chromatin. The characteristic cross-shaped contact enrichment associated with “strong” single genes (see Methods) is markedly reduced in the post-LBC stage, whereas convergently oriented TUs continue to delineate insulating domain boundaries. The first two panels depict genes rescaled to span 1/12 of the figure width, with flanking regions extending ±6 TU lengths from gene ends (2□kb resolution). The third panel shows aggregated maps for convergently oriented gene pairs within a ±500□kb window around their midpoint (5□kb resolution). Insulation scores are indicated in the upper-left corner (see Methods). D) Distribution of domain sizes across different chromosome types. The post-LBC stage shows an increase in mean domain size alongside a reduction in the total number of domains. E) Transcriptional activity of TUs located at domain borders at LBC and post-LBC stages. F) Distribution of insulation scores (see Methods) for domain borders at LBC and post-LBC stages. Lower scores indicate stronger insulation.

Despite the absence of “cross” patterns, domain patterns can be identified on the Hi-C maps of late-diplotene oocytes, and the majority of the boundaries coincide with TDD boundaries of LBC oocytes. However, post-LBC Hi-C maps contain fewer boundaries, resulting in increase of domain lengths (Figure 8D). Visual inspection shows that about half of the neighboring LBC TDDs were fused at the post-LBC stage (44 borders out of 97 examined on chromosome 3, Figure 8B). Genome-wide quantification shows that interaction frequency between neighboring LBC domains in post-LBC oocytes are also increased (2% of contacts between neighboring domains in normal oocytes vs 6% of contacts for late oocytes). Additionally, analysis of the border of TU pairs shows that the activity of weakly expressed TUs decreases, as well as does the insulation (Figure 8E-F). Interestingly, despite the absence of the “cross” pattern of actively transcribed TUs, the insulation by the convergently oriented TUs is still present (Figure 8C).

### Cohesive cohesin barrier function is required for transcription-mediated chromatin domain insulation

Two mechanistic models can explain why strong reduction in global transcription did not substantially diminish insulation attributed to convergent transcription. 1) Even low transcriptional activity in lampbrush chromosomes suffices to produce insulation via interactions between loop-extruding SMC complexes and RNA polymerase. 2) Active transcription redistributes cohesive cohesin towards gene termini, where it acts as a barrier to extrusion. Following transcription shutdown, cohesive cohesin presumably remains in the previously occupied locations, providing insulation even in the absence of hypertranscription. To discriminate between these models, we grouped genes with the same orientation (making gene “tandems”) by length and transcriptional activity (Figure 9A, Methods) and constructed averaged contact maps of chicken oocytes at LBC and post-LBC stages. Model 1 predicts a marked stage-dependent loss of insulation for “weak” tandem pairs but preserved insulation for “strong” tandem pairs. Model 2, in contrast, predicts stronger insulation for long tandem pairs and weak insulation for short tandem gene pairs, with at most a modest overall decrease when transcription is reduced. This pattern arises because longer tandems accumulate more cohesive cohesin at gene termini, thereby forming a more effective barrier to loop extrusion. Under model 2, a global reduction in transcription is expected to attenuate insulation only modestly within each group, since even infrequent transcriptional passages that advance RNA polymerase to gene ends are sufficient to maintain cohesin in a barrier-competent (“locked”) state.

**Figure 9.**
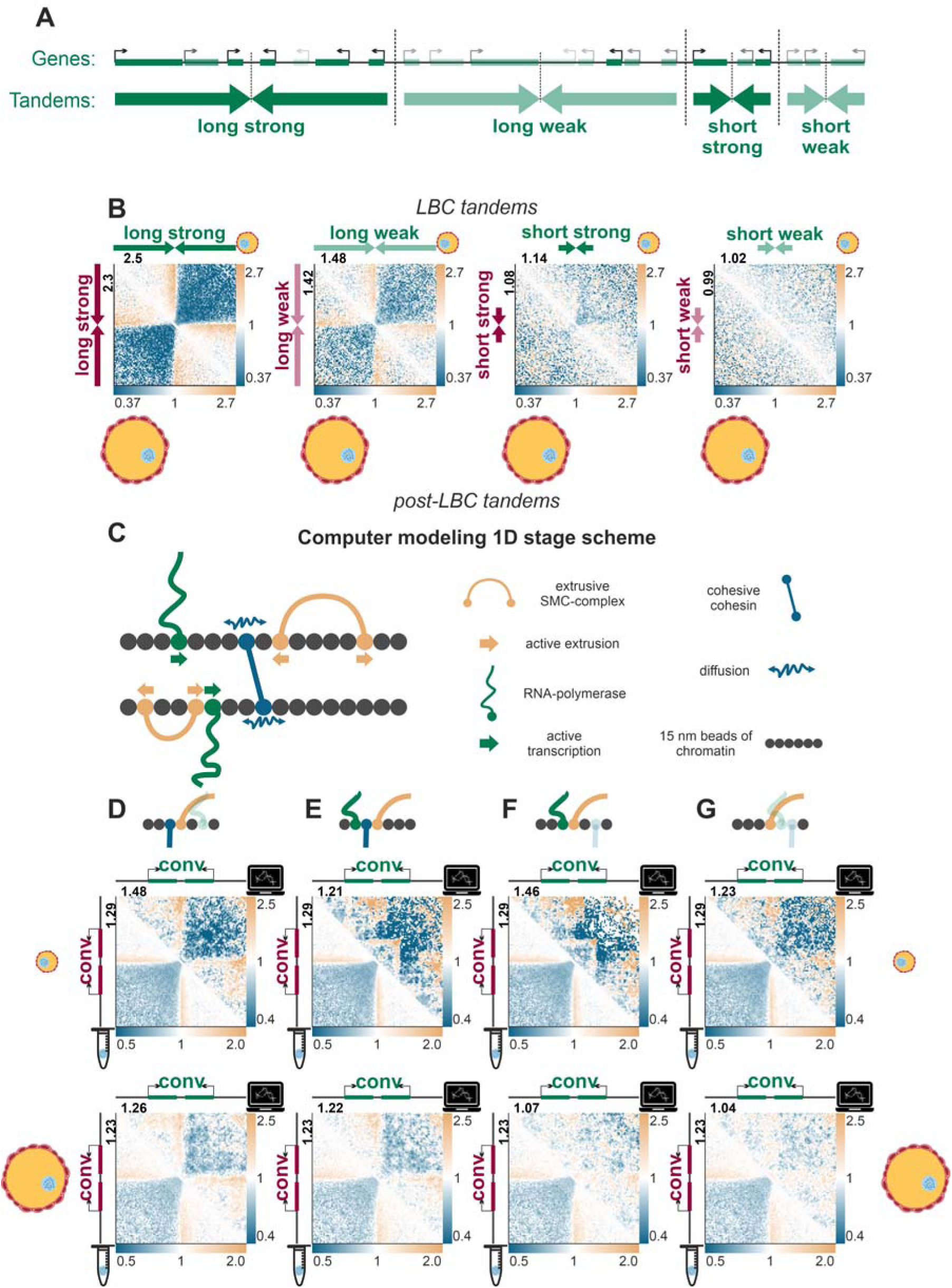
Comparison of the averaged contact maps for convergent gene tandem pairs and computer models at lampbrush chromosome and post-lampbrush chromosome stages. A) Schematic of tandem construction: adjacent co-directional genes are merged into a single tandem unit. Tandem length and “strength” are defined in Methods; transparency encodes transcriptional activity (lower opacity = lower activity). Analyses in this study focus on pairs of neighboring tandems in a convergent orientation. B) Aggregate observed-over-expected (O/E) Hi-C maps for four classes of convergent gene tandem pairs (“long strong”, “long weak”, “short strong”, “short weak”; see Methods) for lampbrush chromosome (LBC, upper triangle) and post-lampbrush chromosome (post-LBC, lower triangle) stages. Insulation scores are shown at the upper left of each panel. All panels show aggregated maps for convergently oriented tandem pairs within a ±500□kb window around their midpoint. Resolution 10 kb. C) Schematic of the one-dimensional simulation framework. D–G) Aggregate O/E maps comparing simulation outputs (upper triangle) versus Hi-C (lower triangle) for LBC (first row) and post-LBC (second row) chromatin; insulation scores as in A; all panels show aggregated maps for convergently oriented gene pairs within a ±500□kb window around their midpoint; 10-kb resolution. Parameter regimes (schematics above each panel set): (D) low loop-extruding SMC bypass probability for cohesive cohesin and near-certain bypass for RNA polymerase; (E) low bypass probability for both; (F) near-certain bypass for cohesive cohesin and low for RNA polymerase; (G) nearcertain bypass for both.

Consistent with Model 2, Figure 9B shows robust insulation at long convergent tandem gene pairs with only a slight decrease from LBC to post-LBC, whereas short “strong” convergent tandem pairs exhibit weak insulation and short “weak” convergent tandem pairs show little to none.

To further test Model 2, we implemented a polymer simulation of a lampbrush-like chromosomal segment (Figure 9C; Supplementary Figure S12; see Methods) with four interaction regimes between loop-extruding SMC complexes and moving barriers (RNA polymerase and cohesive cohesin):

i. low loop-extruding SMC bypass probability for cohesive cohesin, but near-certain bypass for RNA polymerase (Figure 9D);
ii. low bypass probability for both (Figure 9E);
iii. near-certain bypass probability for cohesive cohesin, low for RNA polymerase (Figure 9F);
iv. near-certain bypass probability for both (Figure 9G).

In all regimes, transcribing RNA polymerases push ahead both loop-extruding SMC complexes and cohesive cohesin; initial conditions therefore place cohesive cohesin near tandem ends. The simulations support Model 2: only regimes in which cohesive cohesin functions as a barrier reproduce the post-LBC insulation pattern (Figure 9D–E), and accurate recapitulation of the LBC maps further requires that loop-extruding SMC complexes can efficiently bypass RNA polymerase (Figure 9D). When loop-extruding SMC rarely bypasses RNA polymerase, the model produces artifactual contact enrichment between convergent gene ends (Figure 9E–F).

These integrative analyses and simulations demonstrate that cohesive cohesin functions as a transcription-anchored barrier essential for domain insulation in lampbrush chromosomes. The data strongly support Model 2, where active transcription redistributes cohesive cohesin to gene termini (thus accumulating cohesive cohesin at convergent gene pairs), establishing persistent insulation even after transcriptional reduction. Additionally, the consistency of computer Model 2 with lampbrush-chromosome architecture is supported by the close visual agreement between the simulated structures and FISH-mapping data (see Supplementary Note 3 and Supplementary Figure S13).

## Discussion

### A unified model of lampbrush chromosomes formation

In many vertebrates, oogenesis is accompanied by hypertranscription aimed to accumulate maternal RNA and leading to significant chromosome decondensation into a lampbrush conformation. Our study offers the first comprehensive, genome-wide analysis of chromatin architecture in chicken oocytes during the lampbrush chromosome stage. Despite lampbrush chromosomes being discovered over a century ago (106, 107), our findings provide a new understanding of their architecture and function at unprecedented resolution and across the entire genome. By integrating these results with established knowledge and applying polymer physics, we propose a unified model elucidating the mechanisms underlying lampbrush chromatin organization (Figure 10).

**Figure 10.**
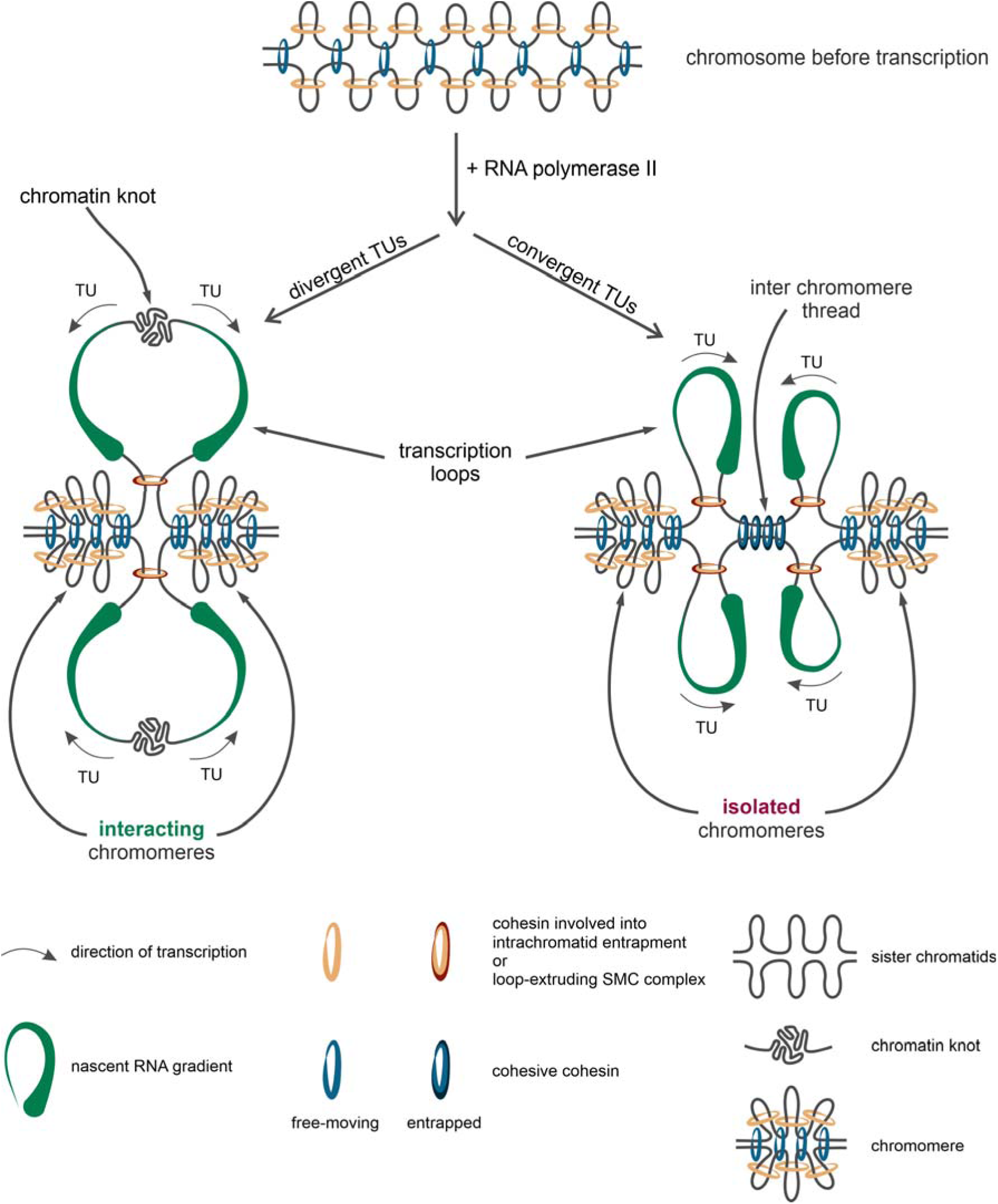
Mechanisms underlying the formation of chromomere-loop complexes in lampbrush chromosomes. Active transcription increases chromatin stiffness and generates pressure that pushes transcription units (TUs) outward from the chromatin axis. Elongating RNA-polymerases remove histones and translocate cohesive cohesin towards the ends of transcription units. In the case of convergent transcribed gene pairs, this results in accumulation of cohesive cohesin complexes between transcription units, anchoring two transcription loops to the chromosomal axes. In the case of divergent transcribed gene pairs, the depletion of cohesive cohesin rings between transcription units results in the detachment of the condensed chromatin knot from the chromatin axes. Loop-extruding SMC complexes active within transcriptionally-silent regions form compact chromatin domains located on chromosomal axis. Transcriptionally-silent regions separated by divergently oriented transcription units can be bridged by extruders moving co-directionally with transcription and/or by intrachromatid entrapment by non-extruding cohesin. However, separation by convergently oriented transcription units forms impenetrable barrier for extrusion due to cohesive cohesin accumulation, precluding interactions between flanking regions. These mechanisms underlie the segmentation of lampbrush chromosome into distinct chromomere-loop complexes.

Hypertranscription is the primary driver of lampbrush chromosome formation, which determines their chromomere-loop organisation (57). Here we highlight several mechanisms linking transcriptional activity to their distinctive organization. First, the accumulation of RNP complexes along the transcription loop axis increases chromatin stiffness (51, 92, 96) and generates pressure that pushes transcription units outward from the chromosome axis. Second, RNA-polymerases travel along a DNA template as a tightly spaced convoy, displacing DNA-bound proteins from transcription units (108, 109). The removal of histones, combined with increased chromatin stiffness, extends the length of transcription loops.

Not all proteins, however, can be displaced by RNA polymerase. Cohesive cohesin, which topologically entraps DNA to hold sister chromatids together, most probably resists disassembly (110). Instead, RNA polymerase translocates SMC complexes along the DNA (110–112), functioning as a motor that slides cohesive cohesin toward the ends of transcription units (110). Similarly, cohesin with extrusion activity is translocated or disassembled during transcription (36–38). This creates directional barriers for the extrusion machinery at transcription unit ends, either through direct interactions with the transcription complex or indirectly via accumulated cohesive cohesin, which blocks reloading of dynamic cohesin extruders (113).

In lampbrush chromosomes, active transcription units remain free of cohesive cohesins, leading to local decoupling of sister chromatids. In contrast, downstream regions of transcription units enriched in cohesive cohesin remain tethered to the chromosomal axis (Figure 10).

A defining feature of LBCs is the presence of transcription loops, both monogenic and multigenic, varying in length (57). Integration of Hi-C maps, oocyte nuclear RNA-seq data, and FISH mapping reveals that transcription loops often appear as Hi-C "cross" patterns, characterized by reduced contacts with adjacent regions. This pattern is consistent with loops being histone-free transcription units extruded from the chromatin axis due to ribonucleoprotein pressure, increased stiffness, and cohesive cohesin depletion (Figure 10).

Transcriptionally inactive regions are enriched with extrusion complexes and cohesive cohesins, leading to their compaction and appearance as contact domains with long-range interactions (Figure 10). FISH analysis shows that transcriptionally inactive regions correspond to intrachromomeric loci (this study and (57, 61)). Our model of inactive chromatin organisation within chromomeres is based on previous models of lampbrush chromomere structure that involve cohesive cohesin and condensin complexes in maintaining lampbrush chromosome axes (50, 114–116). In particular, cohesin complex subunits have been revealed within chromomere cores and in interchromomeric regions (88), while condensin subunit has been found to be enriched within chromomeres (103). Moreover, we have taken into account microscopic observations of the rosette structure of discrete lampbrush chromomeres separated by interchromomeric fibre (48).

Our data reveal how transcription-dependent repositioning of SMC complexes establishes boundaries between chromomere-loop complexes (Figure 10). A single transcribed gene creates a unidirectional barrier at its termination site but does not fully insulate flanking regions, as extrusion complexes can move co-directionally with transcription. Similarly, clusters of closely spaced co-directional genes result in SMC complex accumulation at the end of the last transcription unit, allowing residual interactions between adjacent regions.

In contrast, convergent gene pairs concentrate cohesive cohesin complexes between transcription units, forming an impenetrable barrier and anchoring two transcription loops to the chromosomal axis (Figure 10). Consistent with our Hi-C and cytological data, convergent gene pairs define boundaries between chromomere-loop complexes. Longer genes in a convergent pair increase the likelihood of at least one cohesive cohesin molecule being translocated to the intergenic region, reinforcing the boundary. Thus, the probability of boundary formation scales with gene length.

Unlike intergenic regions between convergent gene ends, those separating divergent gene starts are depleted of cohesive cohesin (Figure 10). Extrusion complexes and histones loaded within these regions mediate their compaction. These regions are tethered to the chromosomal axis if residual cohesive cohesin is present or remain untethered and manifest as a part of lateral loop if it is absent (Figure 10). Consistent with this, we observed small contact domains corresponding to cytologically visible chromatin knots between adjacent transcription units. These knots frequently occur when divergent genes are transcribed in opposite directions from a compact chromatin domain (57), aligning with the described mechanisms.

Extrusion complexes loaded between divergent gene pairs can traverse the genes co-directionally with transcription, bridging flanking regions. Thus, a divergent gene pair can form a loop without strong insulation of neighboring loci. In contrast, as discussed above, convergent gene pairs block extrusion complexes due to cohesive cohesin accumulation between them (Figure 10). This mechanism explains the observed enrichment of convergent gene pairs at contact domain boundaries.

Simulations based on these mechanisms using a physical chromatin model accurately reproduce both the genome-wide statistical properties of chromatin contact distribution and the specific packaging observed at individual loci.

### Opened questions

Although the obtained data unequivocally shows that interaction between RNA polymerases and SMC complexes is the major mechanism explaining LBC formation, nuances of this mechanism remain to be discovered.

It is not clear how many extrusion complexes are involved in LBC organization and what is their subunit composition. It is established that the XCAP-D2 (condensin I subunit) and SMC1A, SMC3, Rad21, STAG1, and STAG2 (cohesin subunits) are present on the lampbrush chromosomes (88, 103). However, it remains unclear whether RAD21 functions solely within cohesive cohesin or also as a part of extrusive cohesin. This leaves the possibility of one to four types of extruders contributing, including cohesin-STAG1, cohesin-STAG2, condensin I, and condensin II, each potentially varying in lifetime, density, and velocity.

From the observed peak positions of the P(s) slope (Supplementary Figure S7A), we estimate an extruder density of approximately 1 per 100–200 Kb in our models. This is comparable to the density of condensin II in interphase or condensin I in prometaphase (117).

Another open question concerns the probability of an extruder traversing chromatin in the direction opposite to RNA polymerase. Banigan et al. (37) assumed the permeability of RNA polymerase to SMC with probability ∼0.01 s^-1, but our analysis suggests that this value is too low to reproduce LBC Hi-C data. In LBCs, RNA polymerase complexes are tightly spaced (109). Given a polymerase velocity of ∼100 bp/sec (93) and the extruder velocity of 675 bp/sec (94) results in the overstep probability of ∼0.15. This oversteps probability indicates that the extruder would frequently become blocked within the gene body. Therefore, the permeability must exceed 0.15 for extruders moving against the direction of transcription to pass through the gene body effectively. This increased permeability would allow extruders to traverse tightly packed transcription units, aligning with the observed chromatin organization in LBCs.

Our model highlights the critical role of cohesive cohesin relocation in response to transcriptional activity, alongside interactions between RNA polymerase and extrusion complexes, in shaping LBC architecture. Interestingly, in yeast, transcription has been shown to promote chromatin loop formation even in the absence of loop extrusion ability in the cohesin variants (112). The authors propose that cohesin can facilitate intrachromatid loop formation by topologically entrapping chromatin near transcription sites. Similarly, the hypertranscription characteristic of the lampbrush chromosome stage could contribute to chromatin loop formation through a transcription-dependent mechanism involving DNA loop capture by cohesin, independent of extrusion activity. This suggests that cohesive cohesins may partially supplement the role of extrusion complexes in establishing LBC architecture, emphasizing a combined mechanism of loop extrusion and transcription-dependent loop capture in shaping these remarkable chromatin structures (Figure 10).

Another unresolved question is why CTCF-mediated insulation, a hallmark of chromatin organization in typical somatic cells, does not shape LBC contact domains in oocytes of birds in contrast to mouse diplotene oocytes at the NSN (non-surrounded nucleolus) stage (42). One hypothesis is that alterations in cohesin subunits during LBC formation, such as unknown post-translational modifications, could result in complexes that lose their CTCF-interacting ability. An alternative explanation is that CTCF is unable to bind active chromatin in LBCs due to the high density of RNA polymerase complexes occupying these regions. This transcription-driven exclusion could prevent CTCF from establishing its characteristic insulatory boundaries. Further investigation is required to determine the precise mechanisms underlying this phenomenon.

### Principles underlying lampbrush chromosome organization are universal across cell types

Although lampbrush chromosomes (LBCs) exhibit a unique type of chromatin organization, the mechanisms underlying their structure are universal and can be observed across various cell types. For instance, transcription loops formed by long, highly expressed genes in somatic interphase nuclei show extensive externalization and reduced cis-contacts, resembling LBC transcription loops (92). Similarly, loss of contacts or chromatin ‘melting’ was observed in the case of highly expressed long neuronal genes (118). LBC transcription loops also resemble actively transcribed genes in polytene chromosomes, which appear as puffs (53). Hi-C maps of *Drosophila* polytene chromosomes revealed absence of TADs or chromatin contact domains in the puffed regions (119). In addition, interbands in polytene chromosomes correspond to inter-TAD regions, which contain clusters of actively transcribed genes with a broad expression pattern (120) or the promoters and 5’ ends of such genes (121, 122). Thus, loss of chromatin contacts within LBC loops is a common feature of highly transcribed genes in different model species and cell types.

Looping of actively-transcribed genes, which do not interact with neighboring transcriptionally inactive chromatin but also do not form interactions with active loops from other domains, resembles the local compartments reported in oocytes (41) and meiotic sperm cells (44). Similar attenuation of long-range interactions within compartments has also been observed in Anopheles mosquitoes (14). Lack of compartments was also detected in the Drosophila polytene chromosomes (119). In all four cases, the chromosomes adopt a rod-like structure, which likely explains the absence of long-range homotopic interactions in these models.

Previous studies have demonstrated that RNA polymerase can act as a "moving barrier" to extruding complexes in bacteria (35), yeast (36), slime mold *Dictyostelium discoideum* (38) and even for vertebrate cohesin (37). However, in vertebrate cells, interactions between cohesin and RNA polymerase generally have a minor impact on chromatin architecture, becoming noticeable only in conditions that extend cohesin’s extrusion activity, such as Wapl+CTCF double knockout, and through extensive data aggregation across the genome (37). In contrast, our findings suggest that extruder-RNA polymerase interactions as well as cohesive cohesin barrier function play a significantly greater role in the organization of lampbrush chromosomes, explaining the majority of short-range interactions in diplotene oocytes. These results suggest that transcription-mediated repositioning of SMC complexes represents an ancient, conserved mechanism shaping chromatin architecture across the tree of life.

## Acknowledgments

This manuscript is dedicated to the memory of Joseph G. Gall. This study was initiated with support of RSF Project №20-64-46021 in 2020-2022. We acknowledge the collective centers of Ministry of Science and Higher Education of the Russian Federation (state project FWNR-2022-0019) for providing facilities for oocyte nuclei isolation and genomic libraries preparation and sequencing. Access to computational resources was provided by Novosibirsk State University, supported by the Ministry of Education and Science of the Russian Federation (FSUS-2024-0018). This research was supported in part through computational resources of HPC facilities at collaborative center «Bioinformatics» ICG SB RAS. Microscopy analysis was performed using the equipment of the Resource Center ‘Molecular and Cell Technologies’ (Saint-Petersburg State University). Part of the bioinformatic analysis and manuscript preparation was supported by grant of the state program of the “Sirius” Federal Territory “Scientific and technological development of the “Sirius” Federal Territory” (agreement №26-03, 27/09/2024). The original manuscript text was composed by authors; proofreading was conducted with the assistance of ChatGPT version 4, after that the text was corrected and edited by the authors.

## Author contributions

V.F. and A.K. conceptualized the study. M.G. conducted the genomics experiments with assistance from A.K., T.K., and A.M., who contributed to oocyte isolation, and A.N., who provided support for staining and NOMe-seq experiments. Cytological analyses, including FISH-mapping experiments, were performed by T.K. and A.K. with contribution from A.M., who performed probe labeling. Bioinformatic analyses were carried out by T.L., with contributions from V.F., A.P., A.N., and V.K. T.L. developed the methodology for polymer simulations and conducted the modeling experiments with input from V.K. T.L. and V.F. drafted the manuscript with contributions by A.K.; authors jointly developed mechanism explaining lampbrush chromosome organization, and A.K. suggested the final scheme of chromomere-loop complex formation. All authors reviewed, revised, and approved the final manuscript.

## Data access

All raw sequencing data and proceed data are available under GEO accession <u>GSE288415</u>.

The program code used for lampbrush chromosome model simulation is freely and publicly available at https://github.com/genomech/LBC_model.

Any additional data in support of the findings of this study can be obtained from the corresponding authors upon reasonable request.

