## Supplementary for "The 3D genomics of lampbrush chromosomes highlights the role of active transcription in chromatin organization"

**Supplementary Note 1. Two types of oocyte Hi-C libraries** **2**
**Supplementary Note 2. The concordance of Hi-C patterns with structural units of lampbrush chromosomes identified by microscopy analysis** **2**
**Supplementary Note 3. Validation of lampbrush chromosome chromatin simulations using FISH-mapping microscopy data in individual regions** **3**
**Supplementary Methods 1. Modeling** **4**
*1D* *4*
*3D* *5*
*Simulated Hi-C contacts* *7*
*Simulated microscopic images* *8*
**Supplementary Figures** **9**
*Figure S1. Schematic of the "Cross Pattern Score" (CPS) computation (see Methods).* *9*
*Figure S2. Single cell Hi-C quality controls.* *11*
*Figure S3. Chromatin organization of lampbrush chromosomes at different genomic distances. Extended version.* *12*
*Figure S4. Comparative analysis of A/B compartmentalization strength across chicken embryonic fibroblasts and oocyte stages.* *13*
*Figure S5. Chromatin domains in chicken lampbrush chromosomes.* *15*
*Figure S6. Comparison between DNA methylation/chromatin accessibility and transcriptional activity.* *16*
*Figure S7. Chromatin domain properties.* *17*
*Figure S8. FISH-mapping of the additional combinations of BAC clone-based probes on lampbrush chromosome 1.* *18*
*Figure S9. The comparison of the Hi-C map patterns with the lampbrush chromosome chromatin domains identified by microscopy visualization for the LBC4 and LBC13.* *20*
*Figure S10. Insulation at various genome features.* *21*
*Figure S11. Examples of de novo annotated genes which are transcribed at the lampbrush chromosome stage and absent in chicken v23 annotation of the ASM2420605v1 genome assembly.* *22*
*Figure S12. Schematic of the physical polymer model used for the 3D stage of lampbrush-chromosome simulations.* *23*
*Figure S13. Comparison of lampbrush chromosome chromatin simulation with FISH-mapping microscopy images.* *25*
*Figure S14. Scaling curves (P(s)) of different stages and types of oocytes.* *26*
*Figure S15. The Hi-C maps averaged near different genes or CTCF sites orientations for chicken oocytes and somatic cells.* *27*
*Figure S16. The Hi-C maps averaged near different genes or CTCF sites orientations for the lampbrush chromosome (LBC) stage oocytes of type 1 and 2, chicken embryonic fibroblasts (CEF) and the in silico mix of the LBC stage oocytes of type 1 with CEF.* *28*
*Figure S17. Examples of the gene pair orientation in borders mistyping.* *29*
**List of Supplementary Tables** **39**
**List of Online Supplementary Materials** **39**
**Bibliography** **40**

### Supplementary Note 1. Two types of oocyte Hi-C libraries

Based on sequencing-derived statistics of the prepared Hi-C libraries, two distinct types of oocytes can be identified. Type 1 oocytes exhibit less than 0.7% interchromosomal contacts, while type 2 oocytes display more than 3.5% interchromosomal contacts. In addition, type 2 oocytes show an enrichment of long-range contacts (see P(s) in Supplementary Figure S14). The origin of this dichotomy remains unclear. Notably, type 2 oocytes at the lampbrush chromosome (LBC) stage demonstrate clustering of domain boundaries highly similar to those observed in chicken fibroblasts, despite the visual differences in their Hi-C contact maps that do not suggest fibroblast contamination. Moreover, microscopic inspection of the samples supports the absence of somatic cell contamination.

However, contact maps averaged around CTCF binding sites in type 2 oocytes resemble a mixture of signals from type 1 oocytes and fibroblasts (1) (Supplementary Figures S15-16), raising the possibility of biological mixing. In contrast, the saddle strength profiles of type 2 oocytes deviate from those of computational mixtures between fibroblast and type 1 oocyte nuclei (Supplementary Figure S4), arguing against simple contamination.

Two alternative explanations can be considered: 1) minor contamination of a single oocyte by a single somatic cell, which appears unlikely given experimental precautions, or 2) type 2 oocytes represent an earlier, pre-lampbrush developmental stage, where residual CTCF-associated contact domains remain detectable—a more plausible scenario supported by the data. For post-LBC oocytes, type 2 likely corresponds to a later developmental stage than post-LBC type 1, as suggested from aggregated observed-over-expected maps (Supplementary Figures S15-16). Only type 1 oocytes were included in the main analyses of contact domains.

### Supplementary Note 2. The concordance of Hi-C patterns with structural units of lampbrush chromosomes identified by microscopy analysis

Four regions of interest covered by BAC clone-based probes were analysed by FISH mapping on isolated lampbrush chromosome preparations (Supplementary Table S3).

Region of interest B (70-73 Mb) on lampbrush chromosome 4 comprises 3 apparent contact domains (Supplementary Figure S9A). A clear “cross” pattern on chromatin contact map can be seen for region 7. In accordance, the actively transcribed gene within this region forms a medium sized transcription loop with a prominent RNP matrix (Supplementary Figure S9B,C).

Regions 1 and 2 also label transcription loops, but with a different density of RNP fibrils, corresponding to the nuclear RNA-seq profile (Supplementary Figure S9A). The RNP matrix for the transcription loop formed by region 1 is very thin, whereas the RNP matrix for the transcription loop formed by region 2 is more prominent.

The intra-domain regions 3, 4, 5 and 6 comprise untranscribed genomic segments together with short transcribed genes (Supplementary Figure S9A). According to FISH mapping on isolated lampbrush chromosomes, the BAC clone-based probe 4 labels a dense DAPI-positive chromomere and is in close contact with region 5 (Supplementary Figure S9C). On the obtained Hi-C maps, regions 3 and 6 correspond to the contact domain boundaries. In our FISH mapping, probe 3 labels two chromomere parts connected by an inter-chromomere thread (Supplementary Figure S9C), while probe 6 labels chromatin knots close to the chromomere and region 5 (Supplementary Figure S9C).

On lampbrush chromosome 13, six BAC clone-based probes were localized by FISH mapping (Supplementary Figure S9F). The regions covered by probes 2 and 5 correspond to the “cross” pattern on the Hi-C maps obtained (Supplementary Figure S9F). According to nuclear RNA-seq data there are two divergent transcription units in each of these genomic regions. FISH mapping revealed that genomic regions 2 and 5 correspond to relatively long transcription loops with tiny chromatin knots in the middle (Supplementary Figure S9D,E). The intra-domain regions 1 and 4 were mainly mapped within small chromomeres but with additional short transcription loops (Supplementary Figure S9D,E). Regions 3 and 6 were selected as regions at the boundaries of chromatin contact domains on the Hi-C maps (Supplementary Figure S9F). In the nuclear RNA-seq profile, there were two convergent transcription units in each of these genomic regions. On isolated lampbrush chromosome preparations, genomic regions 3 and 6 were mapped within parts of the transcription loops; additionally, region 6 was localized within a chromomere (Supplementary Figure S9D,E).

**Supplementary Note 3. Validation of lampbrush chromosome chromatin simulations using FISH-mapping microscopy data in individual regions**

To confirm that our simulations reproduce not only genome-averaged Hi-C patterns but also the 3D architecture of individual loci, we simulated chromatin structures for genomic region on chromosome 1 previously analyzed by FISH (Supplementary Figure S13A). To compare the simulation results with FISH data, simulated chromatin structures were flattened and chromatin segments were color-coded in blue scale based on AT-fraction (to mimic DAPI staining) or assigned pseudocolors matching BAC clone hybridization images (Supplementary Figure S13B-D). To visualize the nascent RNA on a chromatin, we plotted each bead using bead size proportional to the model-predicted ones.

The simulated structures recapitulate hallmark features of lampbrush chromosomes observed by microscopy: chromatin knots (Supplementary Figure S13B), transcription loops (Supplementary Figure S13C) and chromomeres (Supplementary Figure S13D). After planar flattening, the apparent radial extent of loops increases. Consistent with experimental observations (2), cohesin co-localizes with chromomere cores in the simulations (Supplementary Figure S13B).

In agreement with BAC 8 FISH-mapping data, a chromatin knot can appear either on the axis or radially displaced (Supplementary Figure S13B). Notably, in the native 3D (non-flattened) configuration both nodules reside near the axis within a typical chromomere radius. This supports a model in which the positions of chromatin knots on transcription loops (and the loop contour itself) are regulated by SMC-mediated tethering of the loop segments to the chromomeres. The same mechanism is consistent with RNA-FISH mapping for BAC 9 (Supplementary Figure S13C), which reveals an asymmetric loop relative to the axis: on one sister chromatid gene end is tethered to a chromomere, whereas on the other one gene end is not tethered.

Supplementary Figure S13D further illustrates the relative arrangement of multiple BAC clones on LBCs. The simulated geometry matches the overall ordering seen by FISH and additionally indicates the extent to which individual BACs may hybridyze into adjacent chromomeres, occasionally causing two nearby foci to merge or, conversely, producing the visual impression of extra minor foci.

Finally, we demonstrate that polymer models can successfully simulate the 3D structures of complete chicken chromosomes. We found that the 9 Mb chromosome segment models remarkably resemble to the cytologically observed structures of lampbrush chromosome segments (see online supplementary data at https://genedev.bionet.nsc.ru/ftp/by_User/tlagunov/LBCs/htmls/). These models capture the key architectural features of LBCs, providing a robust framework for understanding their unique organization and facilitating direct comparisons with experimental data.

**Supplementary Methods 1. Modeling**

*1D*

The first stage of modeling consists of creating one-dimensional arrays that carry information about the beads of the polymer chain and the objects placed on it: cohesive cohesins, active extruders and RNA polymerases.

The RNA polymerase approximate size is 15 nm (3), thus it covers the ~45 bp of DNA in the transcription process. So, the 45 bp bead of 15 nm size was chosen for actively transcribed regions. We will assume that a chromosome region without transcriptional activity is densely packed with nucleosomes. We used the 5 times condensation for the chromatin that is packed by nucleosomes (estimation based on (4)). Thus, the inter-gene chromosome regions were represented by beads containing 225 bp of 15 nm size.

LBC genes differ from each other by their length and transcription activity (i.e. total RNA-seq signal, see Methods). We assume the constant RNA polymerase velocity (~100 bp/sec (5)), thus the simulated genes should differ from each other by RNA polymerase density. Using the available marking of transcription units with the high activity in combination with their *rnap_sep* it will be possible to immediately select chromatin regions where there will be almost no possibility of nucleosome landing due to the high density of RNA polymerases (*rnap_sep* < 300 bp). This estimation contributes with the maximum reported distance between RNA polymerases of newt lampbrush chromosomes (~80 nm equal to ~245 bp for naked DNA, (6)). Dividing the length of such a region by the length occupied by a single RNA polymerase (45 bp) we obtain an integer number of beads on it, where each one accounts for approximately 45 bp. For less actively transcribed genes (*rnap_sep* >= 300 bp) the linear condensation approximation was used (*1*). The direction of movement and the probability of landing for future RNA polymerases are also indicated here. An analogical operation is carried out with regions outside of highly active transcription units where each bead accounts for approximately 225 bp. All the resulting beads are collected into a single array – one sister chromatid.

1) $$BS = 45\min\left(\frac{\lambda_t}{300}, 5\right)$$

Landing probability for RNA polymerases calculates as $\frac{\tau}{V_p\lambda_t}$, where $\tau$ - time step duration in seconds, $V_p$- polymerase velocity, $\lambda_t$- estimated distance (separator) between polymerases for gene *t*.

At each time step (equal to 0.2 seconds), RNA polymerases land on the bead at the beginning of the transcription unit with a given probability if it is free. Previously landed RNA polymerases move to the next bead after a sufficient number of time steps, depending on the size of the current bead. The total accumulated length of the RNA tail for each RNA polymerase is recorded. Intron sequences are excised at the designated locations. If a bead in the direction of RNA polymerase movement is occupied by cohesin or loop-extruding factors (LEF), then the RNA polymerase begins to push this subunit and all subsequent subunits in the direction of movement until they run into another RNA polymerase. After the collision of SMC subunits, the RNA polymerase does not stop moving, removes LEFs located on its path and ignores cohesive cohesins.

At the initial moment of time, cohesins are loaded onto the chromatids. One part of the complex is loaded on the first chromatid, and the second one is symmetrically loaded on the other. The cohesin packing density is equal to 100 Kb (estimation of 8-10 cohesins per Mb in late prometaphase (7)).

All active extruders of the model are also loaded onto chromatin at the initial time with a packing density of 90 Kb. As mentioned earlier in the Methods, condensins are loaded only on a pair of beads free of other objects, placing one of their subunits on one bead (thereby occupying it). In accordance with their speed, each time step attempts to move one of the subunits to the third bead (achieving the step length of ~40nm (8)) from the current position in the direction of movement, while the other does not move. But if the third bead is occupied by another LEF or RNA polymerase then the LEF tries to step on the second bead or on the first bead (if the second one is occupied too). If all three beads in the direction of moving are occupied, then the LEF subunit is stuck. Regardless of the movement success, the active subunit is fixed in place and the cycle is repeated for the other, thus effectively reproducing bidirectional extrusion. If there is an RNA polymerase in the path of movement, the active extruder can step over it with probability of 0.01 (based on (9)). Passing through another LEF or cohesive cohesin is not performed.

*3D*

Simulation of objects with a large number of beads requires large computing power (for example, a 20 Mb region corresponds to ~500,000 beads), so the optimal approach is to cluster them into new bigger beads that reproduce properties of a group of 10 beads. Physical parameters of new beads are calculated based on fractal dimensions.

The bond distance between neighboring beads is calculated as follows. First, the average fractal dimension ($D_f$) for the cluster is calculated ($D_f = 1$ corresponds to a straightened DNA strand, such places are usually encountered during active RNA polymerase binding, $D_f = 3$ corresponds to the densest chromatin packing, places with a high concentration of nucleosomes) (*2*). In case of different bead types in one cluster (active genes, less active genes, non-transcribed chromatin), the average value of fractal dimension in a cluster was used. Then, the estimated fractal dimension value was used to calculate the linear size of the cluster $L$, simulated as bond length (*3*).

2) $$D_f = \frac{BS_c}{90} + 0.5$$

3) $$L = r_b \cdot cs^{\frac{1}{D_f}}$$

Here $D_f$ is the fractal dimension, $BS_c$ is the size of the bead cluster *c* in bp, $r_b$ is the size of the base bead, *cs* is the number of beads in the cluster.

Another parameter of the bead cluster is the radius. It is determined from the following considerations: if this is a cluster of only nucleosomes, then the fractal dimension is maximal ($D_f$ = 3) and the formula (*3*) can be used, otherwise it becomes necessary to take into account the nascent RNA-protein complexes, simulated as chains hanging from the polymerases. The effective size of beads formed from such chains is calculated using the following formulas (*4,5*) (based on (6)).

4) $$R_t = \frac{a^{\frac{5}{4}} N^{\frac{3}{4}}}{\pi^{\frac{1}{4}} \lambda_t^{\frac{1}{4}}}$$

5) $$N = \frac{l_b \cdot 0.33}{2a}$$

Here $N$ is the total number of chain links (monomers) emanating from the RNA polymerases of one cluster; $a$ is the size of one monomer equal to 15 nm (estimated from SEM microphotographs in (10)), $l_b$ is a total length of RNA on the RNA polymerase in base pairs (1 bp length is 0.33 nm).

The above $\lambda_t$ was calculated based on the marked transcription units as $k_{stage}/FPKM_t$ normalized to the minimum possible distance between RNA polymerases - 45 bp (the distance between neighbor RNA polymerases that touch each other). $k_{stage} = 65$ for the lampbrush chromosome stage and $k_{stage} = 1200$ for the post-lampbrush chromosome stage (the values were estimated based on the linear compaction of transcription loops). It is worth noting that in the case of overlapping sections of transcription units, the section with the smaller FPKM was excluded from the calculations.

The number and lifetime of LEFs were determined in two steps: a baseline estimate of the number of extruders based on the peak in the derivative P(s), which corresponds approximately to the average loop size (11), on the other hand, an abrupt drop in P(s) after 1 Mb probably indicates the removal of condensins (the end of their lifetime). After that, the manual fitting of the lifetime and number of condensins was applied to the simulated region with BAC clones (achieving the correspondence of Hi-C contact maps and FISH microscopic images).

At the stage of spatial modeling, the bead-clusters of the chromatids are arranged in the form of a flat fractal (the sister chromatid is arranged parallel at a distance equal to the length of the cohesin), which ensures optimal initial conditions (compact size, sister chromatid spatial separation and absence of entangles).

LEFs and cohesins create harmonic bonds between given beads, at each time step all bonds are reassigned in accordance with 1D arrays. After each such reassignment of bonds, 1500 iterations of the molecular simulation are performed, which is sufficient for the system to reach local equilibrium (energy minimization at the initial conformation is performed in 15000 iterations to avoid residual stresses in newly formed bonds).

The potentials effects on the beads movement are (Supplementary Figure S12):

- Linear connectivity. Represents an existence of a polymer chain (DNA) or SMC complex (~40 nm) connecting bead clusters (*6-8*).
- Volumetric attraction and repulsion of bead clusters containing only nucleosomes (*9*). Here, the adhesion of nucleosomes caused by liquid-liquid phase separation, the presence of short nucleosome tails (weak effect, its influence on the observed structures is assumed to be insignificant), and the effects of volume repulsion were taken into account.
- Repulsive volumetric interaction caused by bead clusters with RNA polymerases. The presence of the RNA-protein complex hanging as a chain from the RNA polymerases must be taken into account when calculating interactions between beads, and included in the interaction potential (*10*).

6) $$U_{harm} = \frac{b(l-L)^2}{2}$$

7) $$D = wiggle \cdot L$$

8) $$b = \frac{2U_{harm}}{D^2}$$

9) $$U_{nucl}(r) = H(\sigma + R_1 + R_2 - r)E_{nucl}\left(\frac{r}{R_1+R_2} - 1\right)^2$$

10) $$U_{poly}(r) = H(R_1 + R_2 - r)E_{poly}\left(\frac{r}{R_1+R_2} - 1\right)^2$$

Here *b* is an elasticity coefficient, *l* is a distance between adjacent bead clusters, $L$ is a link length between beads, *wiggle* is an wiggle percentage of length (0.02 for LEFs and cohesins), $D$ is a distance at which bond energy equals kT (1 nm for all bonds between bead clusters), $H$ is a Heaviside step function, $\sigma$ is an adhesion term representing nucleosome tails, *r* is a distance between bead clusters.

The energies $E_{nucl}$, $E_{poly}$ and $\sigma$ were selected for the best fit to the Hi-C data and microscopic images. Interestingly, the adhesion distance of nucleosomes given as $\sigma$ converges with the length of short nucleosome tails (12).

The influence of the persistent length is directly determined by an extremely small parameter (1e-3); it is believed that the repulsion between beads fully realizes persistence length.

The masses of the beads were taken to be equal to 0.1 Da, which allowed the influence of Brownian motion to be increased many times, thereby reducing the time required for calculation.

The chromosome 1 part from 55 Mb to 64 Mb was simulated. The 3D simulation stage was initiated 5 maximal LEF lifetimes after LEF loading and then run for an additional 2 maximal LEF lifetimes—sufficient for the polymer to reach a stable 3D configuration. For each parameter setting, we performed 20 independent replicas, yielding 20 single-oocyte fragments per setting. The parameter space followed a 4×2 design: four encounter regimes governing how loop-extruding SMC complexes bypass (i) RNA polymerase and (ii) cohesin in its cohesive state, crossed with two RNA polymerase densities corresponding to the LBC and post-LBC stages. The four regimes were:

(i) low loop-extruding SMC bypass probability for cohesive cohesin (p=0.05 when the next bead was available) and near-certain bypass of RNA polymerase (p=1.0 when the next bead was available);
(ii) low bypass probability for both barriers (cohesive cohesin p=0.05 when the next bead was available; RNA polymerase p=0.2 when the next bead was available);
(iii) near-certain bypass probability of cohesive cohesin (p=1.0 when the next bead was available) and low bypass of RNA polymerase (p=0.2 when the next bead was available);
(iv) near-certain bypass probability of both barriers (p=1.0 when the next bead was available).

<u>*Simulated Hi-C contacts*</u>

When talking about the captured conformations of "oocytes", the question arises of collecting contact maps under conditions of different bond lengths between beads and the number of contacts taken into account. The contacts themselves are collected as beads located at a maximum distance of 10 minimum cluster radius equal ~160 nm (minimum cluster radius is ~16 nm). Thus, we obtain a matrix of the number of pairwise contacts of beads. To reduce the map to a more realistic form, the number of contacts must be aligned with the number of *Dpn*II restriction sites (normally no more than 4 counts for each *Dpn*II site per single oocyte). Each value of contacts $C_{ij}$ of bead pair *i* and *j* was divided by square root of their coverage multiplication to force the sum of contacts of each bin close to 1. Next, value $C_{ij}$ was multiplied by square root of the desired bin coverage multiplication which is equal to the bead size $B_i$ in base pairs divided by 256 (average *Dpn*II site presence in genome) and multiplied by 4 (*11*). This procedure was applied twice to reduce standard deviation of bins coverage. Knowing the sizes of the beads in base pairs, the resulting matrix is converted into a contact map in bp units and saved as ".mcool" maps with an experiment-like resolution list.

11) $$C_{ij} = \frac{C_{ij}\sqrt{\frac{4B_i}{256} \cdot \frac{4B_j}{256}}}{\sqrt{cov_i \cdot cov_j}}$$

*Simulated microscopic images*

Chromosome preparations examined in light microscopy are essentially flattened for the purpose of fixation, labelling, presentation in a more convenient form for visual analysis and due to the limitations of microscopic equipment. During this process, some three-dimensional patterns are distorted and in the case of lampbrush chromosomes, they can become more pronounced. For example, loops with high transcriptional activity located close to each other could be more relaxed in 3D, but in the process of flattening, the displaced volume of RNA is forced to redistribute in the 2D dimension, creating additional pressure both inside and outside the loops. Such loops can become more elongated and at the same time be observed as lateral, rather than superimposed on the core. The model objects we created are intact three-dimensional conformations of chromatin (it is possible to collect artificial contact maps from them). To reproduce microscopic images, we acted on this conformation with an external force along the Z axis (since the simulated chromosome is usually elongated along the X axis):

12) $$U_{flat}(z) = H(|z|) \cdot z^2 \cdot E_{flat}$$

Here $U_{flat}$ is flattening potential along the z axis, $E_{flat} = 10^{-6}$is a force multiplier chosen to be soft enough to smoothly compress the chromatin over $10^7$ molecular dynamic steps(compression time) without introducing significant distortions into it. *H* - Heaviside function.

**Supplementary Figures**

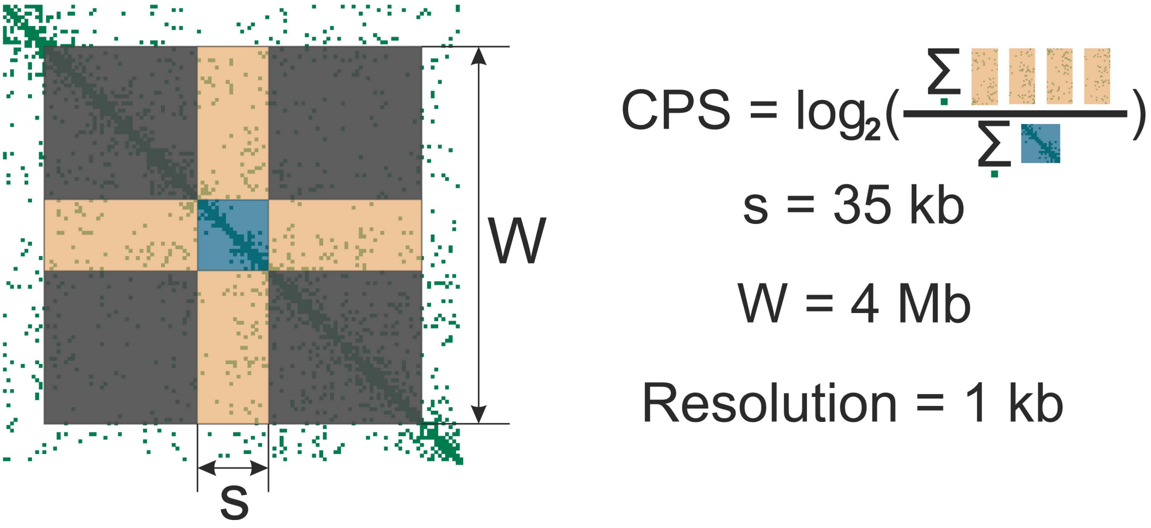


**Figure S1. Schematic of the “Cross Pattern Score” (CPS) computation (see Methods).**

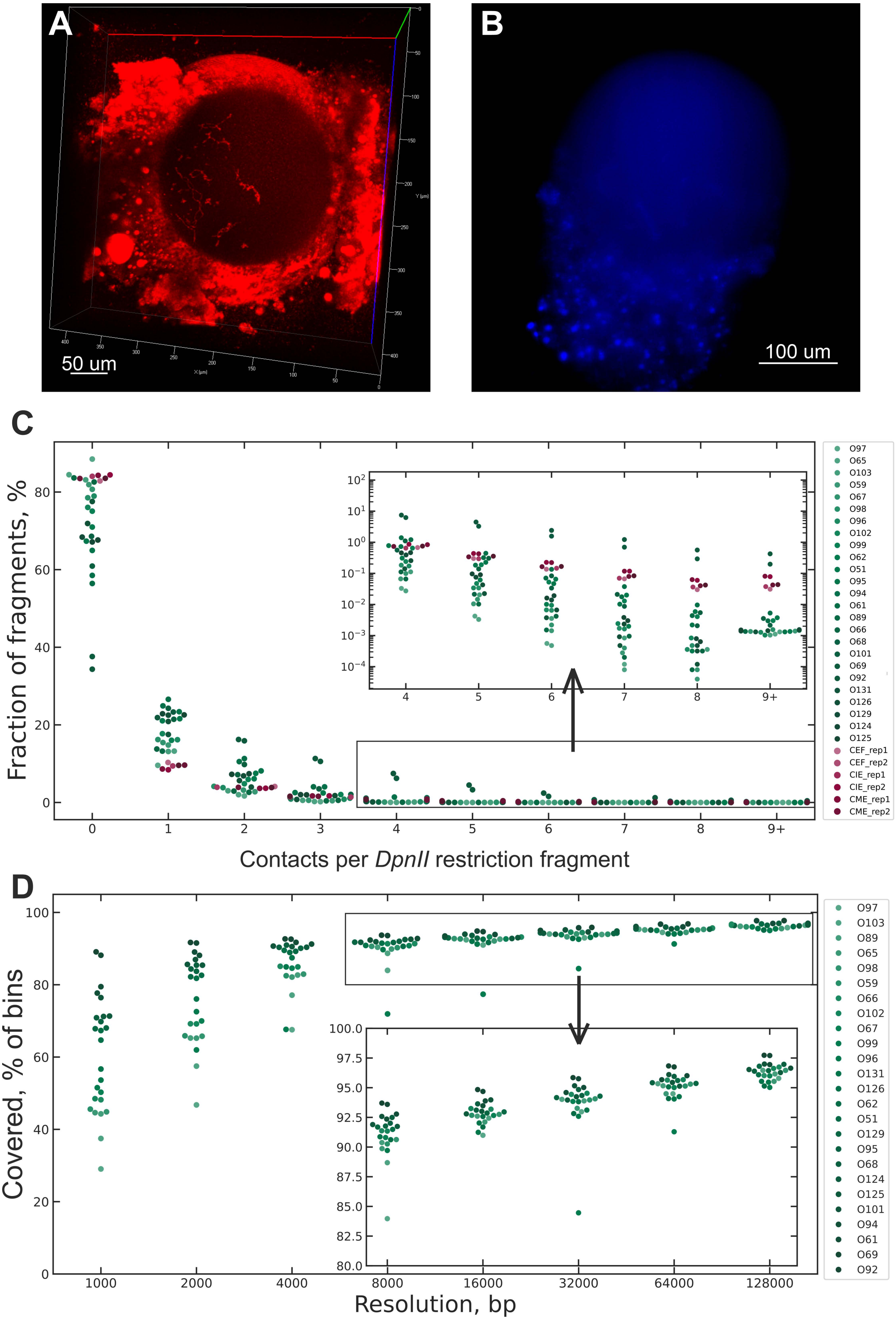
A
50 um
B
100 um
C
Fraction of fragments, %
Contacts per DpnII restriction fragment
0
1
2
3
4
5
6
7
8
9+
O97
O65
O103
O59
O67
O98
O96
O102
O99
O62
O51
O95
O94
O61
O89
O66
O68
O101
O69
O92
O131
O126
O129
O124
O125
CEF_rep1
CEF_rep2
CIE_rep1
CIE_rep2
CME_rep1
CME_rep2
D
Covered, % of bins
Resolution, bp
1000
2000
4000
8000
16000
32000
64000
128000
O97
O103
O89
O65
O98
O59
O66
O102
O67
O99
O96
O131
O126
O62
O51
O129
O95
O68
O124
O125
O101
O94
O61
O69
O92

**Figure S2. Single cell Hi-C quality controls.**

A) Lampbrush chromosomes stained with SYTO 61 (red) after formaldehyde fixation preserve their morphology.

B) DAPI-stained sample containing chicken oocyte nucleus with manually added erythrocytes.

C) Fraction of *Dpn*II fragments with number of Hi-C contacts ranging from 1 to 8 (or more) after duplicate filtering of single oocytes, chicken embryonic fibroblasts (CEF, see Methods), chicken immature and mature erythrocytes (CIE and CME, see Methods).

D) Fraction of bins with at least one measured interaction shown for each single nucleus Hi-C sample at different resolutions.

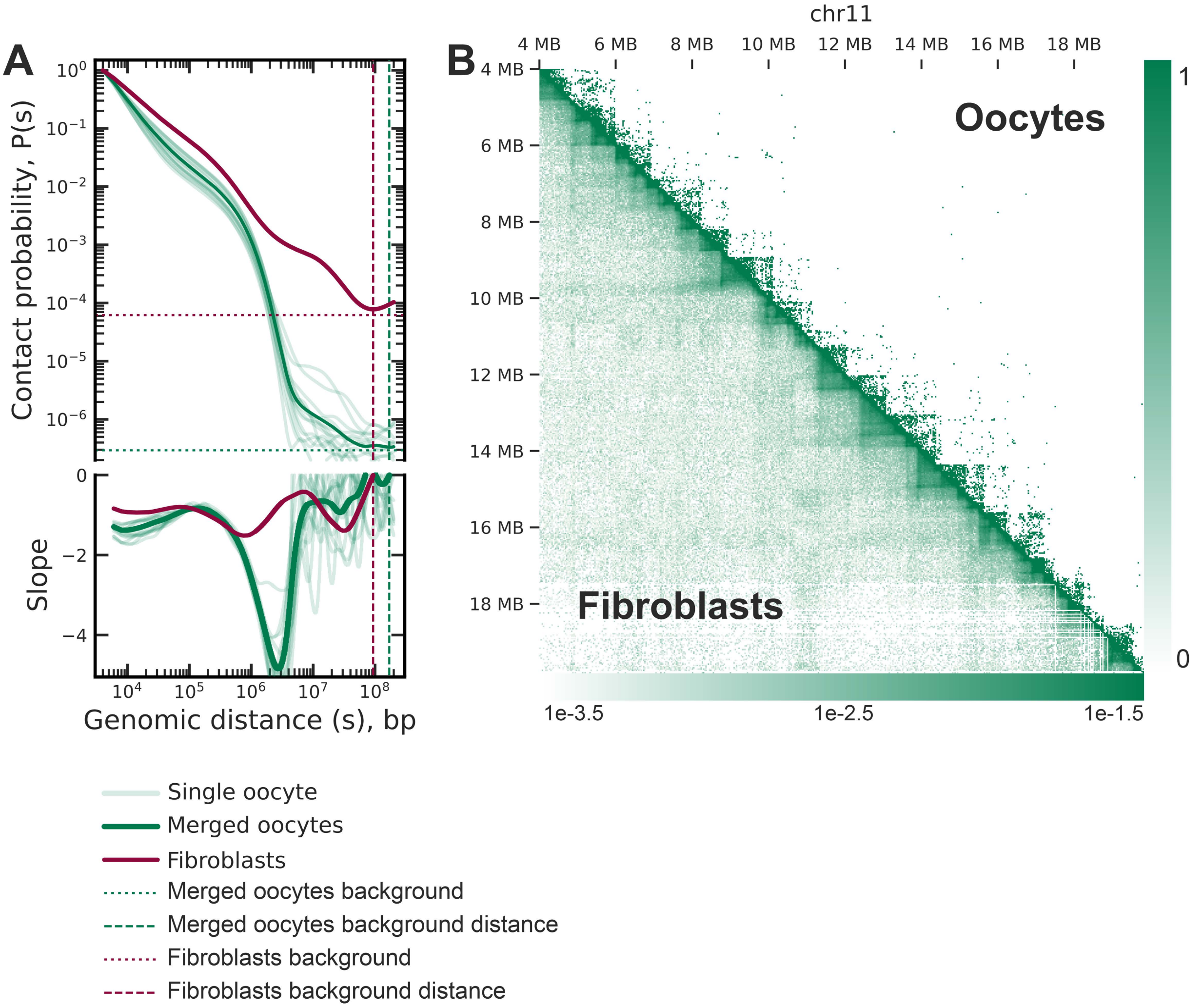


**Figure S3. Chromatin organization of lampbrush chromosomes at different genomic distances. Extended version.**

A) Contact probability versus the genomic distance for single and merged oocyte data (lampbrush chromosome stage, LBC) and chicken embryonic fibroblasts (see Methods). Horizontal dashed lines indicate background inter-chromosomal contact probabilities, while vertical dashed lines denote genomic distances at which these background levels are reached.

B) Hi-C contact maps comparing pseudo-bulk chicken oocytes and fibroblast cells for chromosome 11. The fibroblast map was balanced using *cooler balancing* tool (see Methods). Each bin represents 32 Kb.

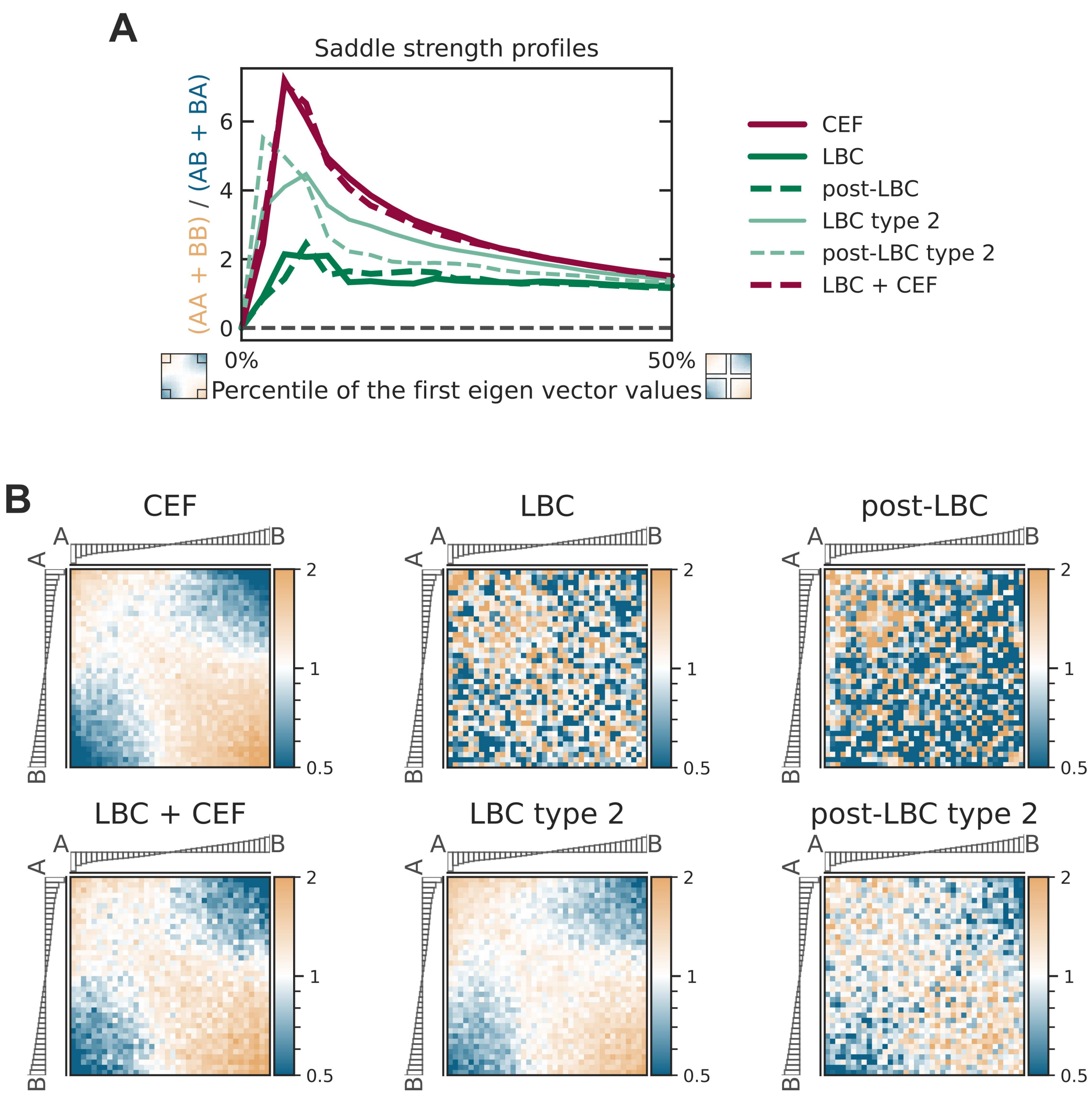


**Figure S4. Comparative analysis of A/B compartmentalization strength across chicken embryonic fibroblasts and oocyte stages.**

A) Saddle strength profiles derived from Hi-C data for the following conditions: downsampled bulk chicken embryonic fibroblasts (CEF, see Methods), oocytes at the lampbrush chromosome (LBC) stage, post-LBC oocytes, LBC and post-LBC oocytes of type 2 (see Supplementary Note 1), and a merged dataset combining LBC and CEF (representing a putative LBC type 2 chromatin state). The X-axis indicates the percentile of the first principal eigenvector used for binning, with increasing percentiles corresponding to broader averaging windows. The Y-axis reflects the compartmentalization strength, calculated as the ratio of intra-compartment (AA + BB) to inter-compartment (AB + BA) contact frequencies. Eigenvector decomposition was performed on CEF Hi-C data at 100 Kb resolution.

B) Saddle plots representing log-transformed observed-over-expected contact enrichments across percentile bins of the first eigenvector values. Each heatmap shows average compartmental interaction strength across all autosomes for each pair of percentile groups. Marginal barplots display the mean eigenvector values for each percentile group. Color intensity reflects the normalized interaction enrichment, with higher values corresponding to stronger compartmental segregation.

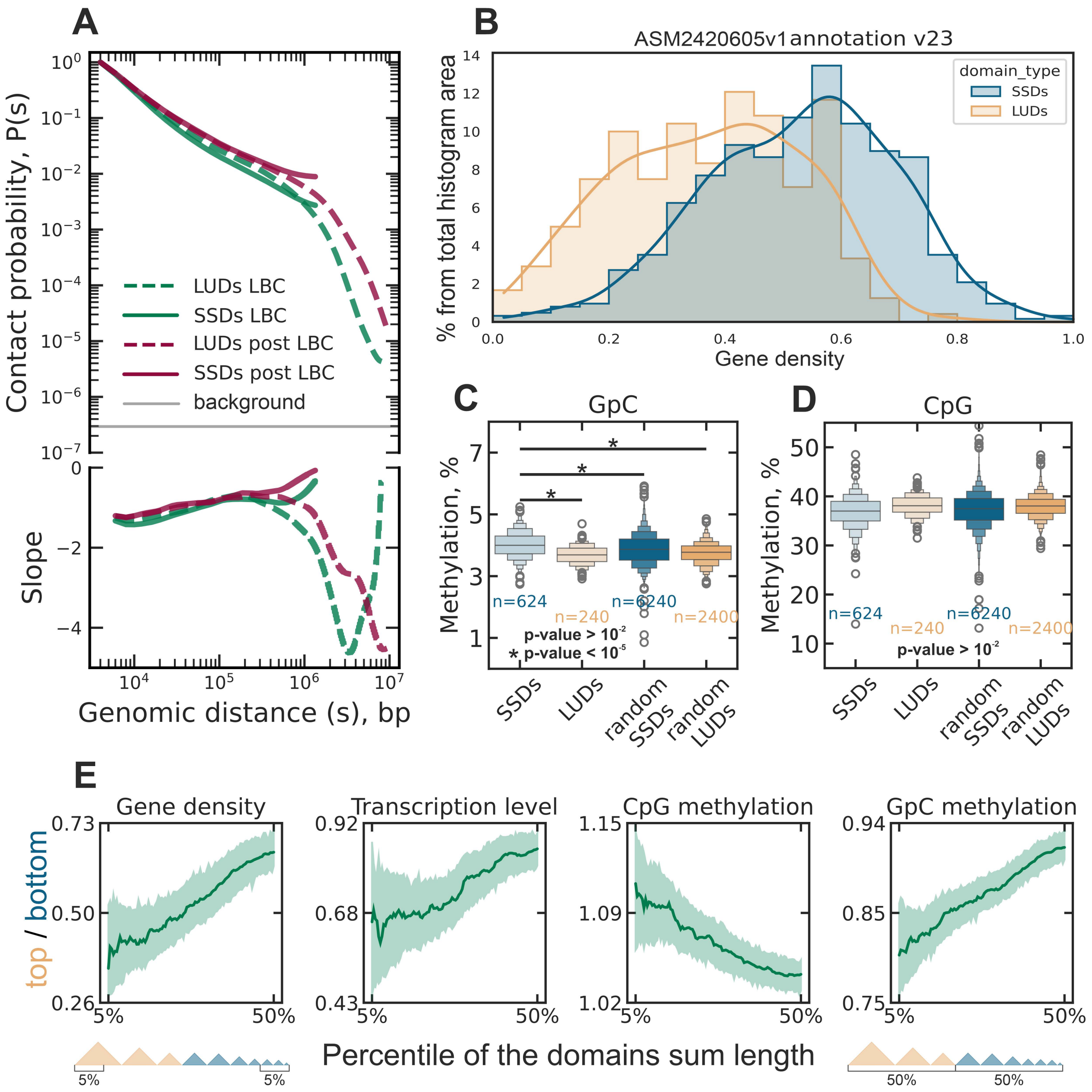
A
Contact probability, P(s)
Slope
Genomic distance (s), bp
LUDs LBC
SSDs LBC
LUDs post LBC
SSDs post LBC
background
B
ASM2420605v1annotation v23
% from total histogram area
Gene density
domain_type
SSDs
LUDs
C
GpC
Methylation, %
n=624
n=240
n=6240
n=2400
p-value > 10^-2
* p-value < 10^-5
SSDs
LUDs
random SSDs
random LUDs
D
CpG
Methylation, %
n=624
n=240
n=6240
n=2400
p-value > 10^-2
E
Gene density
Transcription level
CpG methylation
GpC methylation
top / bottom
Percentile of the domains sum length
5%
50%

**Figure S5. Chromatin domains in chicken lampbrush chromosomes.**

A) Contact scaling curves (P(s)) for LUDs and SSDs in normal and late oocytes.

B) The gene density distribution in LUDs and SSDs (based on ASM2420605v1 assembly, v23 annotation (13)).

C) The GpC methylation distribution in LUDs and SSDs (with randomly shuffled domains as control). The GpC methylation profile was obtained from NOMe-seq experiment (see Methods).

D) The CpG methylation distribution in LUDs and SSDs (with randomly shuffled domains as control). The CpG methylation profile from NOMe-seq experiment was merged with the methylation profile from (14) (see Methods).

E) Percentile–coverage curves for gene density, transcriptional output, CpG methylation, and GpC methylation. Domains were ranked by length, and for each genome coverage fraction (X = 0–50%), the largest and smallest subsets whose cumulative lengths equaled X% of the genome were selected. For each feature, the length-weighted mean values were calculated in both subsets and plotted their ratio (long/short), with 95% confidence intervals estimated by bootstrap resampling.

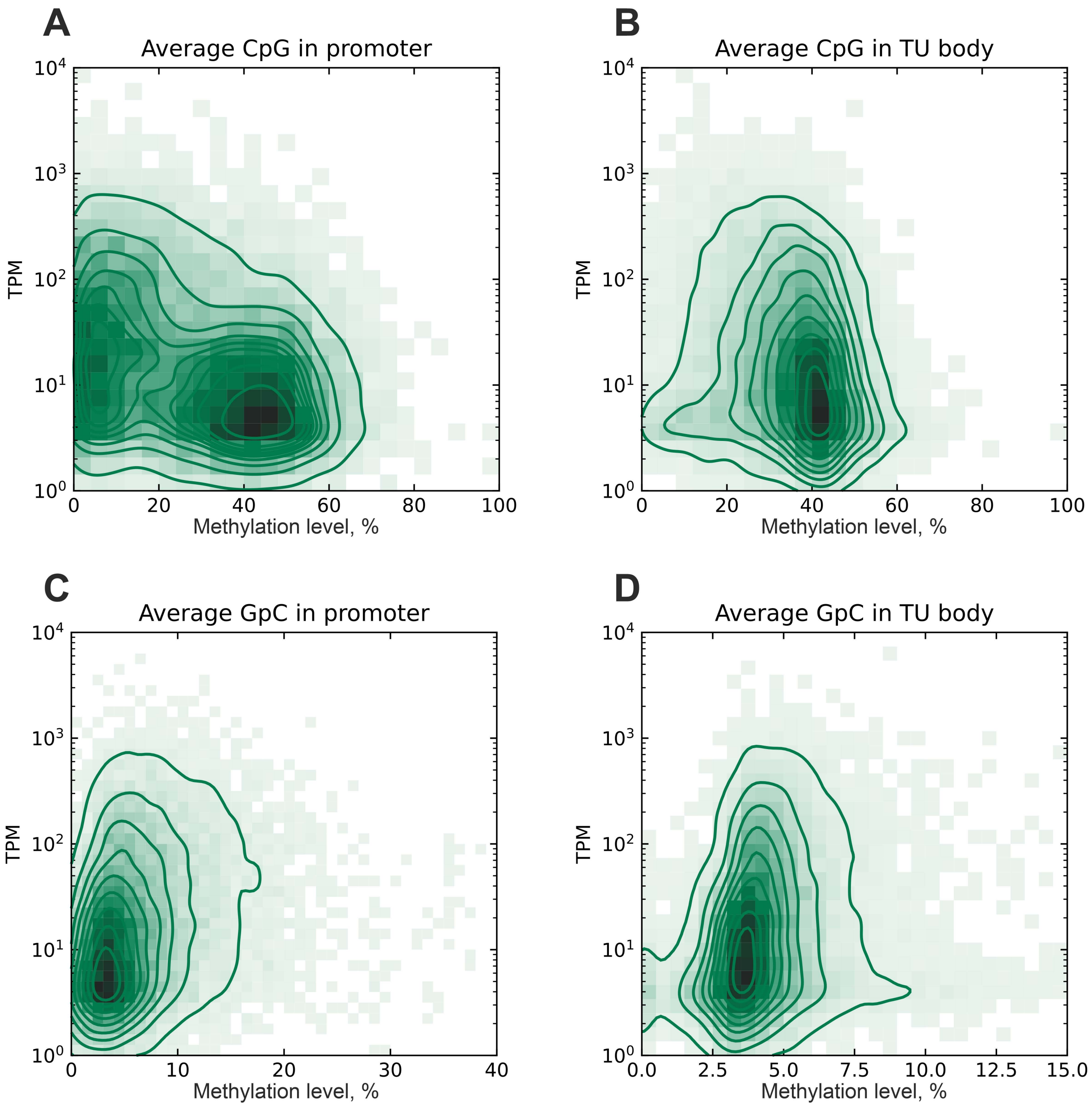


**Figure S6. Comparison between DNA methylation/chromatin accessibility and transcriptional activity.**

CpG methylation level in promoter (A) and gene body (B) vs the transcription activity. GpC methylation (NOME-seq measured proxy of chromatin accessibility) level in promoter (C) and gene body (D) vs the transcription activity.

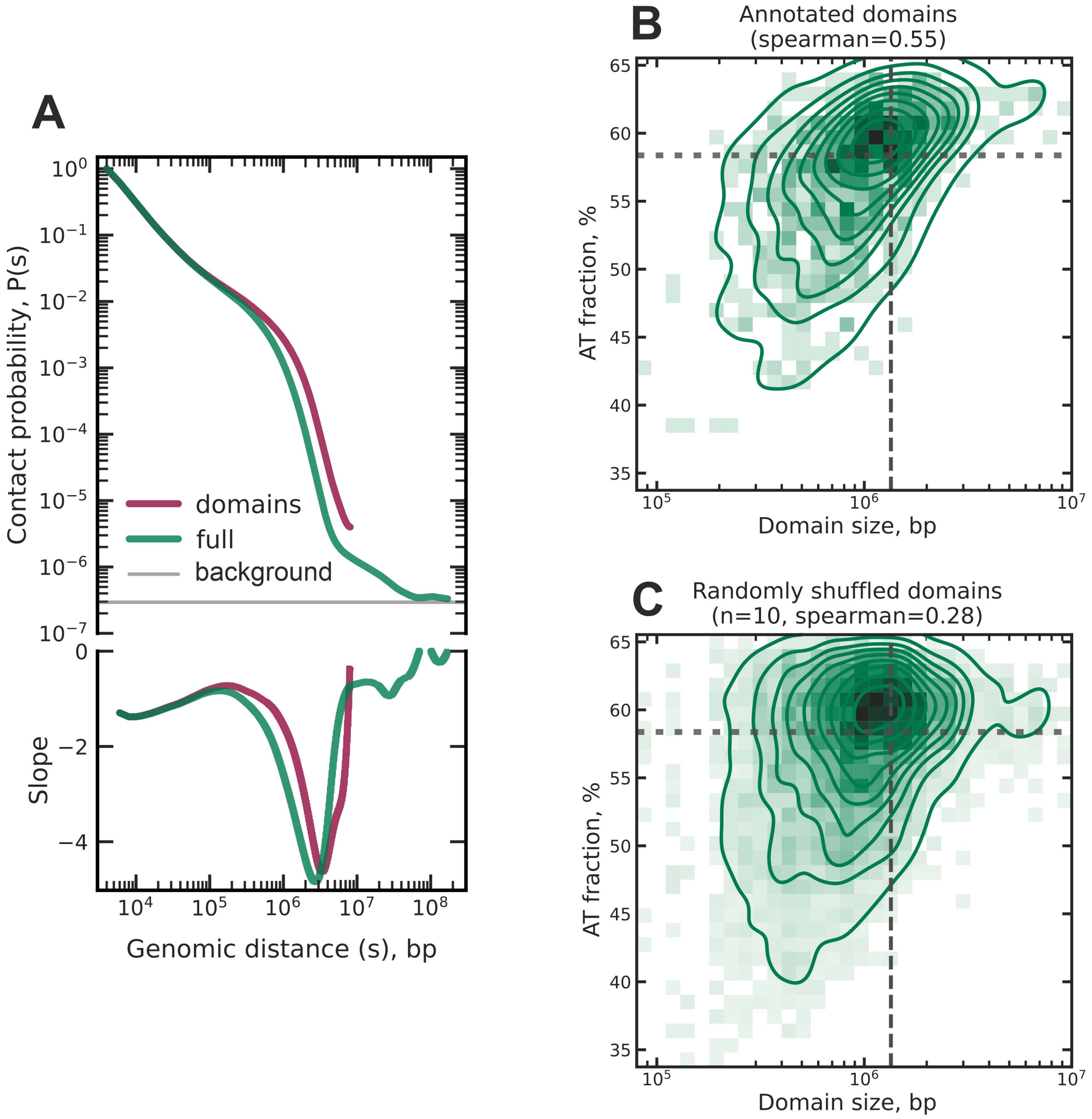


**Figure S7. Chromatin domain properties.**

A) Scaling curves (P(s)) comparing intra-domain contacts and all chromosomal contacts for normal oocytes.

B) Domain AT fraction vs length. Vertical dashed line demonstrates the N50 of domain sizes (threshold between LUDs and SSDs). Horizontal dashed line demonstrates average AT fraction for the chicken genome.

C) Same as B for randomly shuffled domains.

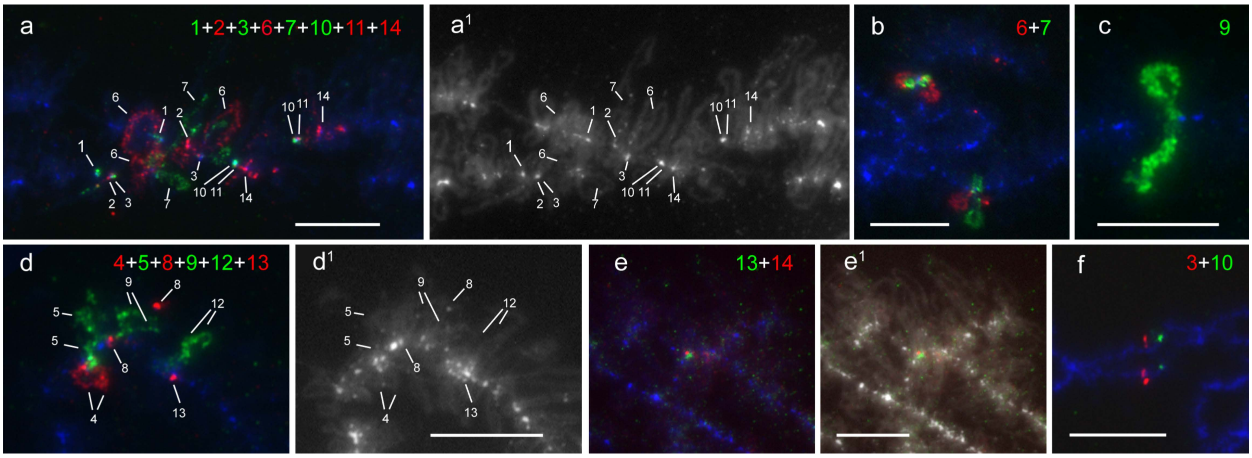


**Figure S8. FISH-mapping of the additional combinations of BAC clone-based probes on lampbrush chromosome 1.**

FISH mapping images for different combinations of BAC clone-based probes on lampbrush chromosome 1 (probe combinations of the corresponding colours are indicated on the panels). Genomic positions of BAC clones and corresponding Hi-C map pattern are shown on the Figure 5A. Chromosome fragments are oriented with their left end toward the left telomere. DNA+RNA FISH is shown on panels a, b, d, e, f, g; RNA FISH is shown on panel c. All panels include DAPI signal (blue or grayscale). On the panels a, a1 and d, d1 fluorescent signals from each of BAC clone-based probes are indicated. Panels a1 and d1 represent grayscale DAPI images with the positions of FISH-signals seen on a and d, correspondingly. On the panels c, d and d1 half-bivalents are shown. Schematic representation of chromomere-loop organization for a fragment of lampbrush chromosome 1 derived from FISH mapping data is shown on the Figure 5C. Scale bars: 10 μm.

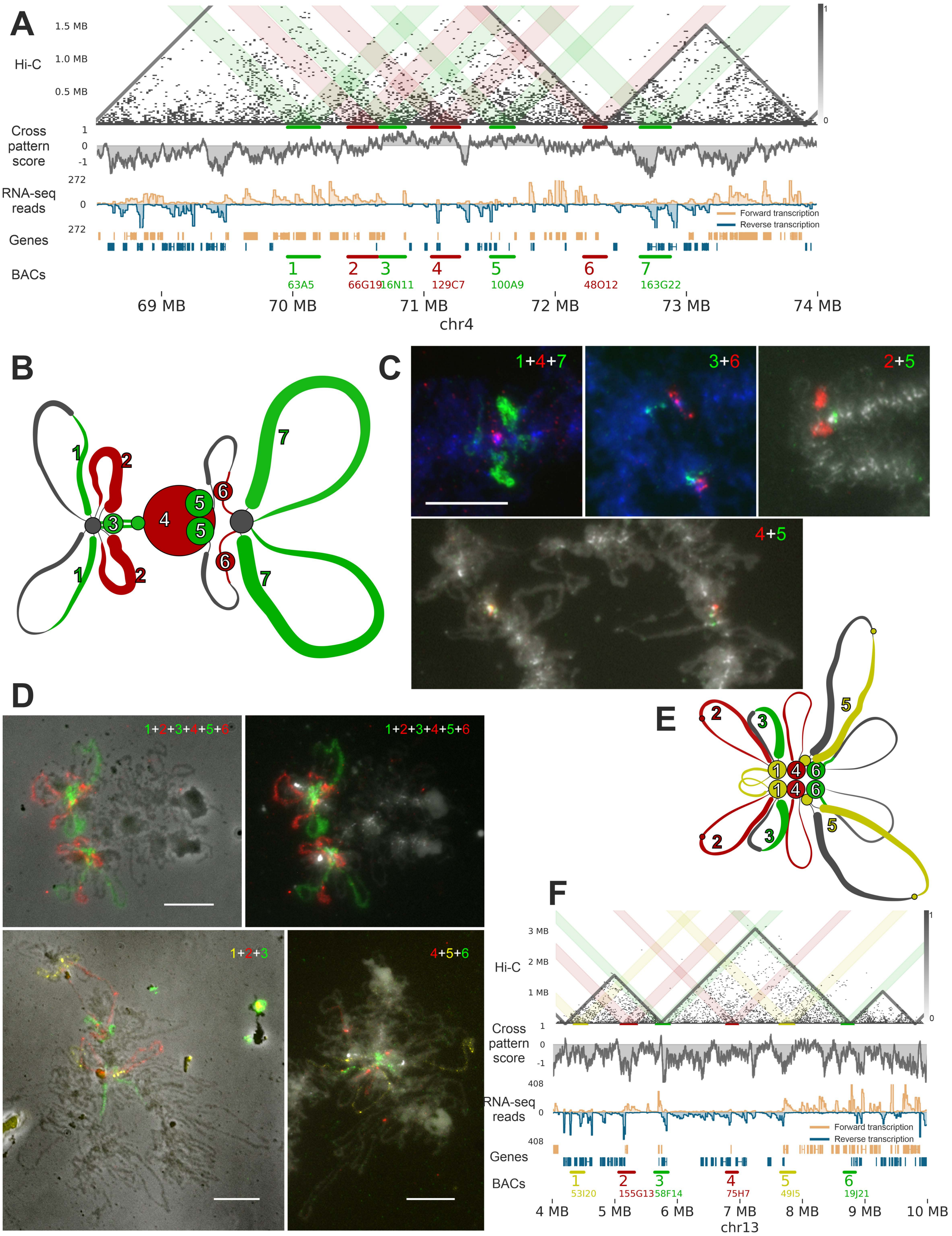
A
Hi-C
1.5 MB
1.0 MB
0.5 MB
Cross pattern score
RNA-seq reads
272
Genes
BACs
Forward transcription
Reverse transcription
1 63A5
2 66G19
3 16N11
4 129C7
5 100A9
6 48O12
7 163G22
69 MB
70 MB
71 MB
72 MB
73 MB
74 MB
chr4
B
C
1+4+7
3+6
2+5
4+5
D
1+2+3+4+5+6
1+2+3
4+5+6
E
F
3 MB
2 MB
1 MB
Hi-C
Cross pattern score
RNA-seq reads
408
Genes
BACs
Forward transcription
Reverse transcription
1 53I20
2 155G13
3 58F14
4 75H7
5 49I5
6 19J21
4 MB
5 MB
6 MB
7 MB
8 MB
9 MB
10 MB
chr13

**Figure S9. The comparison of the Hi-C map patterns with the lampbrush chromosome chromatin domains identified by microscopy visualization for the LBC4 and LBC13.**

A) From top to bottom respectively: the oocyte nuclei Hi-C map of the chromosome 4 ROI B, with the colored highlights depicting interactions of the regions covered by BAC clone based probes; Cross Pattern Score (see Methods); total stranded RNA-seq data for lampbrush stage oocyte nuclei (15); annotation of transcription units (based on *de novo* transcriptome annotation, see Methods); numbers and names of BACs (see Table S3). Resolution 16 kb.

B) The scheme of the chromomere-loop organisation of lampbrush chromosome 4 ROI B, based on the FISH-mapping data (half-bivalent is shown). The numbers indicate the location of BAC clone based FISH-probes aligned on chromosome 4 ROI B.

C) Fragment of chicken lampbrush chromosome 4 showing DNA+RNA FISH mapping images of different BAC clone based probes combinations in the ROI B. Either half-bivalent or two half-bivalents are shown. Fluorescent images are merged with chromatin stained with DAPI (blue or grayscale).

D) Fragments of or whole chicken lampbrush chromosome 13 showing DNA+RNA FISH mapping images of different combinations of BAC clone based probes. Either half-bivalent or two half-bivalents are shown. Fluorescent images are either merged with phase contrast images or with chromatin images stained with DAPI (grayscale).

E) The scheme of the chromomere-loop organisation of lampbrush chromosome 13 fragment, based on the FISH-mapping data (half-bivalent is shown). The numbers indicate the location of BAC clone based FISH-probes aligned on chromosome 13.

F) From top to bottom respectively: the oocyte nuclei Hi-C map of chromosome 13 fragment, with the colored highlights depicting interactions of the regions covered by BAC clone based probes; Cross Pattern Score (see Methods); total stranded RNA-seq data for lampbrush stage oocyte nuclei (15); annotation of transcription units (based on *de novo* transcriptome annotation, see Methods); numbers and names of BACs (see Table S3). Resolution 16 kb.

Scale bars – 10 μm.

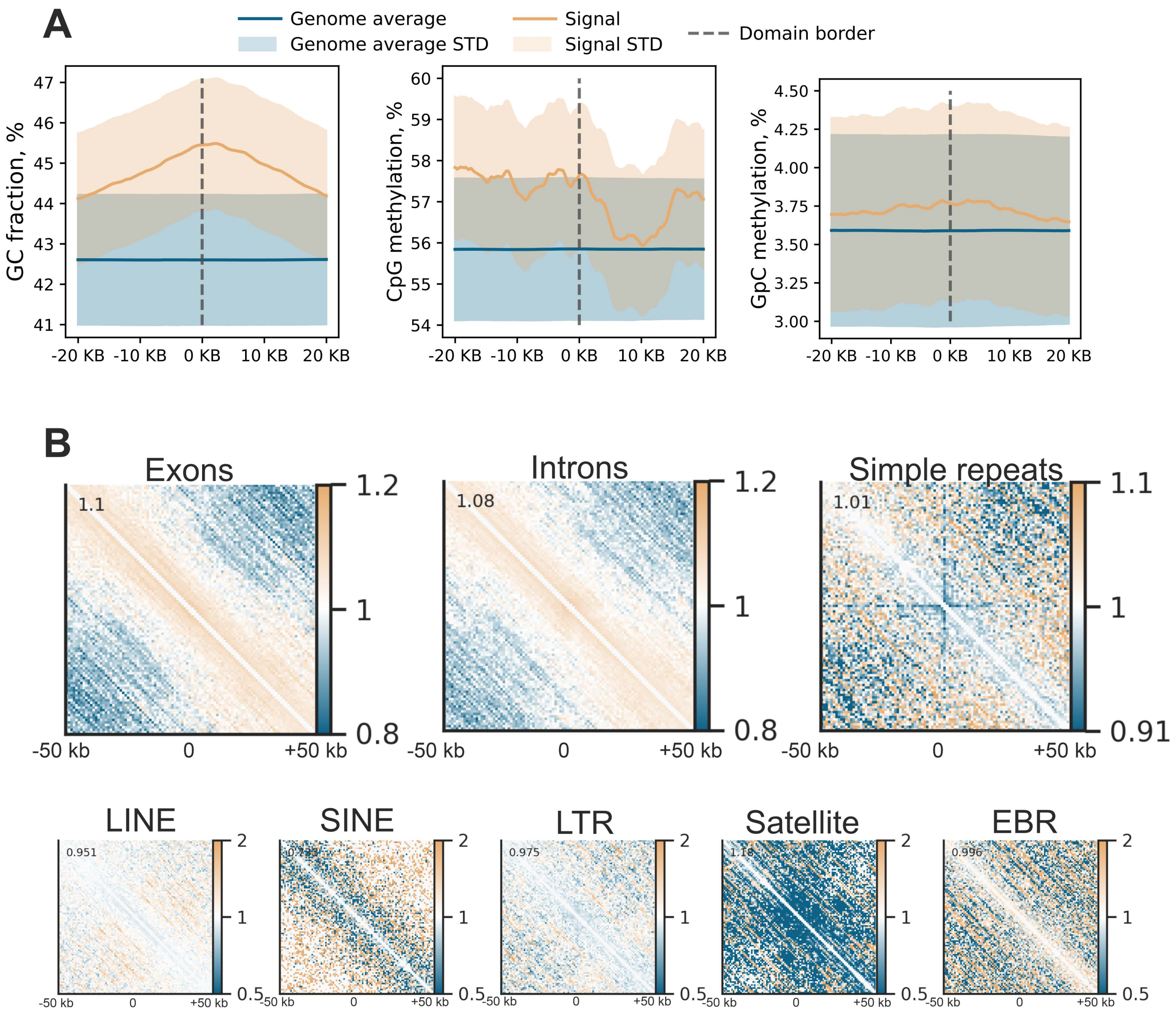


**Figure S10. Insulation at various genome features.**

A) GC fraction, CpG and GpC methylation across the domain boundaries.

B) The Hi-C contact maps for chicken oocyte nuclei averaged across different annotated genome elements

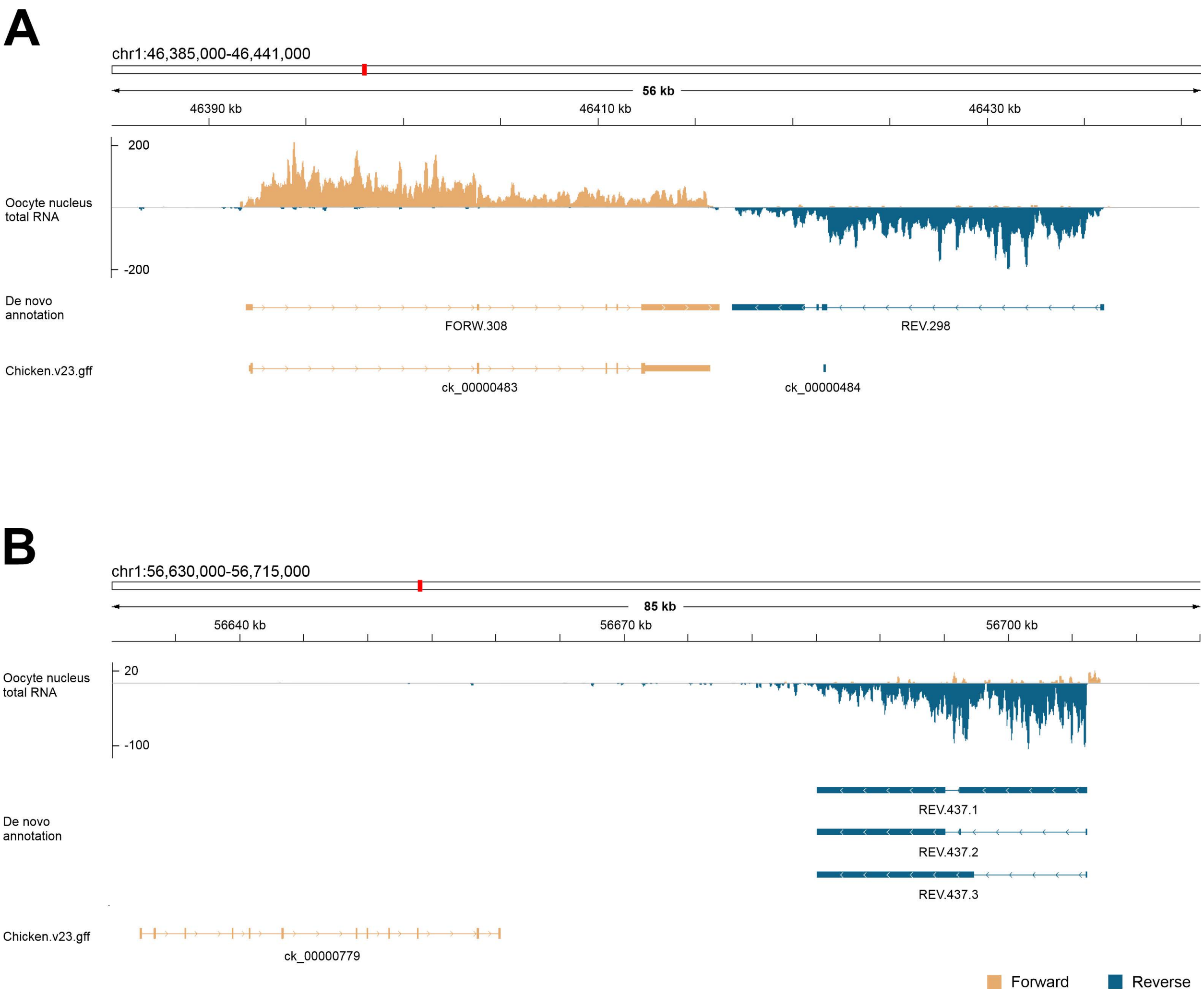


**Figure S11. Examples of de novo annotated genes which are transcribed at the lampbrush chromosome stage and absent in chicken v23 annotation of the ASM2420605v1 genome assembly.**

A) De novo annotated gene FORW.308 corresponds to ck_00000483 in chicken v23; de novo annotated gene REV.298 is absent in chicken v23 annotation.

B) ck_00000779 is not expressed at the LBC; de novo annotated gene REV.437 is absent in chicken v23 annotation. The track "De novo annotation" depicts genes annotated in this work (see Methods). The track "chicken.v23.gff" depicts the chicken v23 annotation of the ASM2420605v1 genome assembly (13). Oocyte nucleus total RNA-seq data was taken from (15).

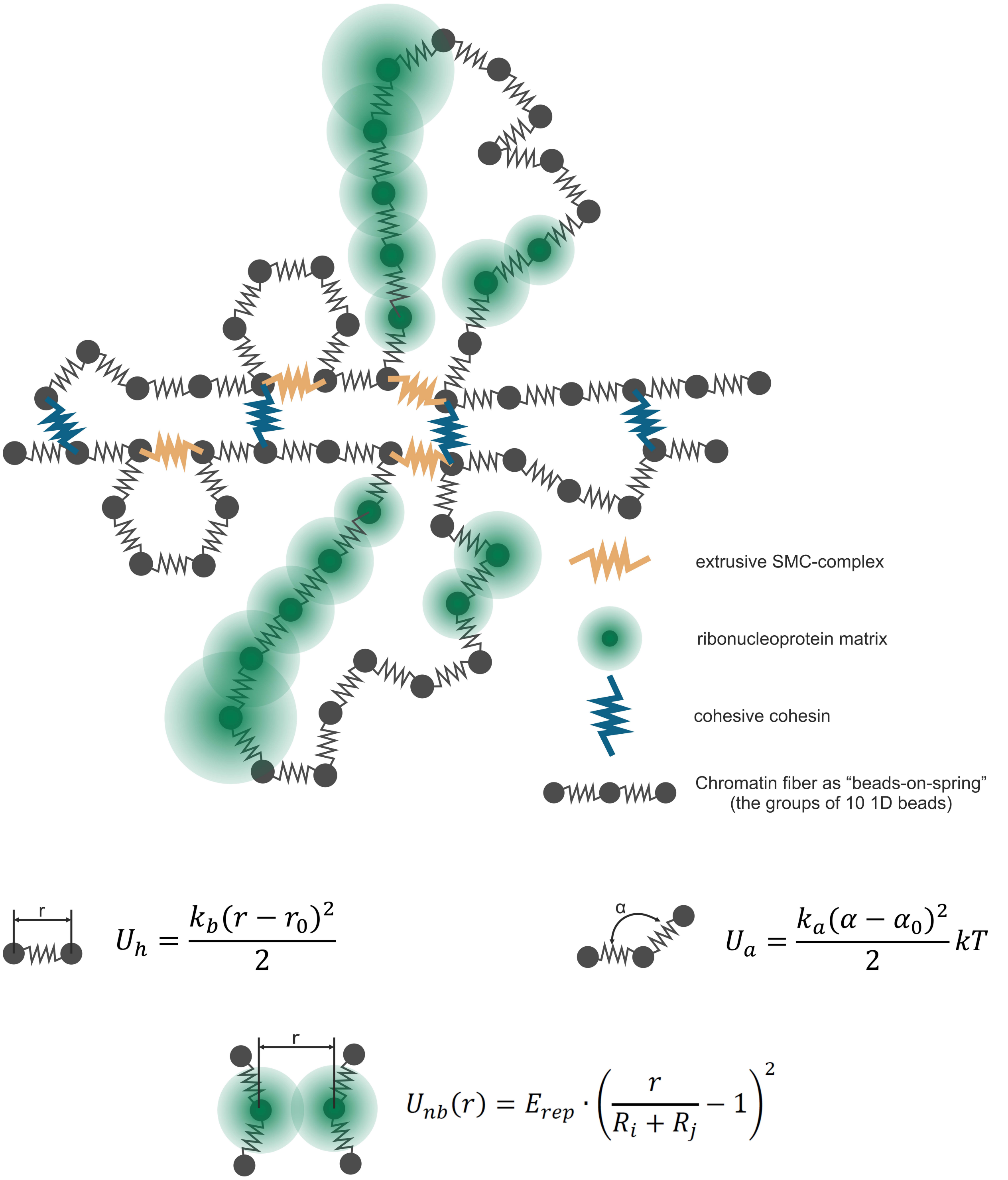


**Figure S12. Schematic of the physical polymer model used for the 3D stage of lampbrush-chromosome simulations.**
Chromatin is represented as a bead-on-spring polymer (groups of ten 1D bins per 3D bead). SMC complexes are modeled as additional harmonic bonds connecting non-adjacent beads. The ribonucleoprotein (RNP) matrix is implemented as soft, mutually repulsive domains whose force decays approximately linearly with separation (i.e., a quadratic repulsive potential). The expressions show the functional form of the bonded (stretch and angle) and non-bonded repulsive interactions; parameter definitions and values are provided in Methods and Supplementary Note 3.

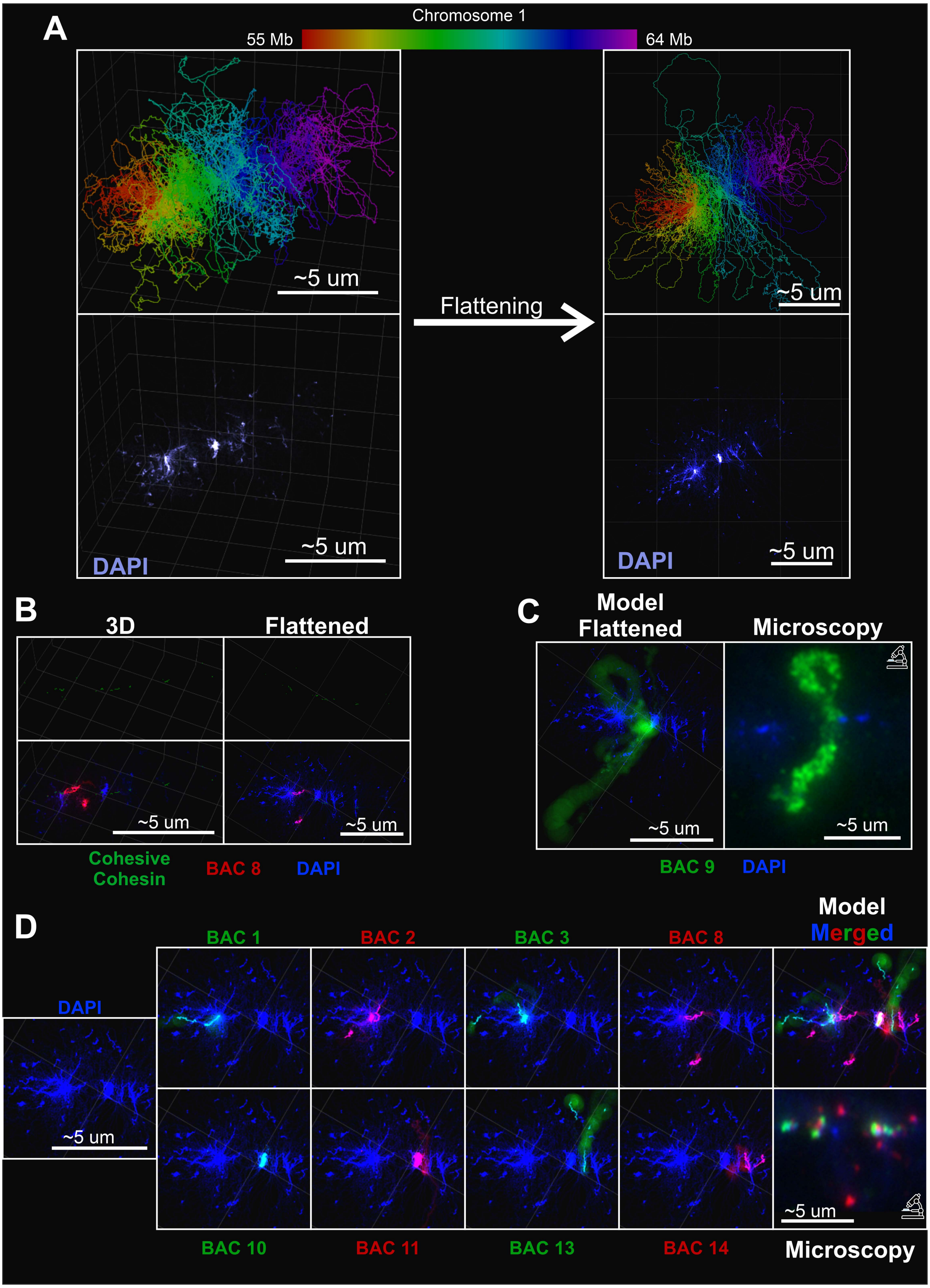
A
Chromosome 1
55 Mb
64 Mb
~5 um
Flattening
DAPI
B
3D
Flattened
Cohesive Cohesin
BAC 8
C
Model Flattened
Microscopy
BAC 9
D
BAC 1
BAC 2
BAC 3
Model Merged
BAC 10
BAC 11
BAC 13
BAC 14
Microscopy

**Figure S13. Comparison of lampbrush chromosome chromatin simulation with FISH-mapping microscopy images.**

A) Visualization of the polymer model for chromosome 1, 55–64 Mb. Left: 3D configuration in free space; right: the same configuration after flattening (see Methods). Top row: segments are colored by genomic coordinate (rainbow colormap); bottom row: segments are shaded by AT-fraction (DAPI-like intensity).

B) Comparison of the 3D configuration and its flattened configuration. Top panels: only cohesive cohesin complexes are shown. Bottom panels: the same views with DAPI-like intensity and an overlaid simulated FISH signal for BAC 8 (see Methods).

C) Comparison of BAC 9 and DAPI signals in the flattened simulation and in the corresponding microscopy image. Simulated panels display DAPI-like intensity with the FISH-like BAC 9 locus overlaid; experimental panels show DAPI staining and BAC 9 RNA FISH.

D) Comparison of the flattened simulation with the corresponding microscopy image (see Methods). The experimental panel shows DAPI staining with DNA+RNA FISH for BAC clones 1, 2, 3, 8, 10, 11, 13, and 14. The modeled panel displays DAPI-like intensity for the same genomic interval with the corresponding BAC loci overlaid. For the modeled dataset, we present DAPI-like alone, DAPI-like with each BAC shown individually, and the merged overlay of all signals.

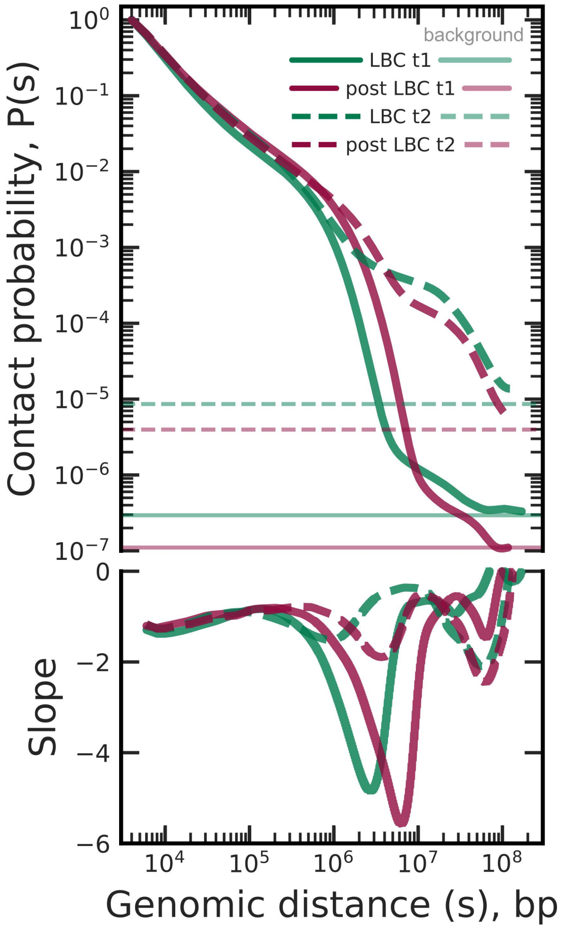


**Figure S14. Scaling curves (P(s)) of different stages and types of oocytes.**
See Supplementary Note 1 for details of oocyte types

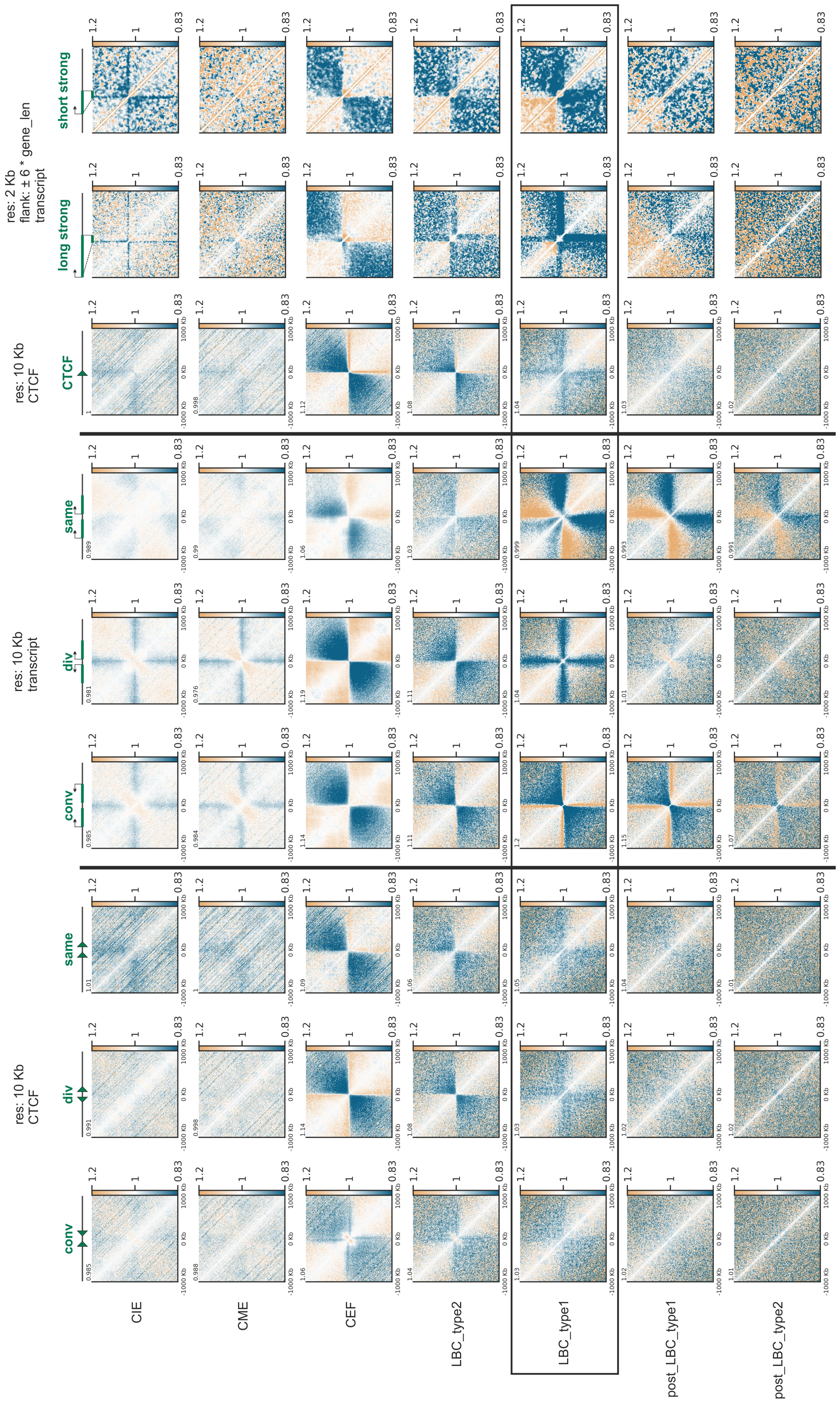


**Figure S15. The Hi-C maps averaged near different genes or CTCF sites orientations for chicken oocytes and somatic cells.**

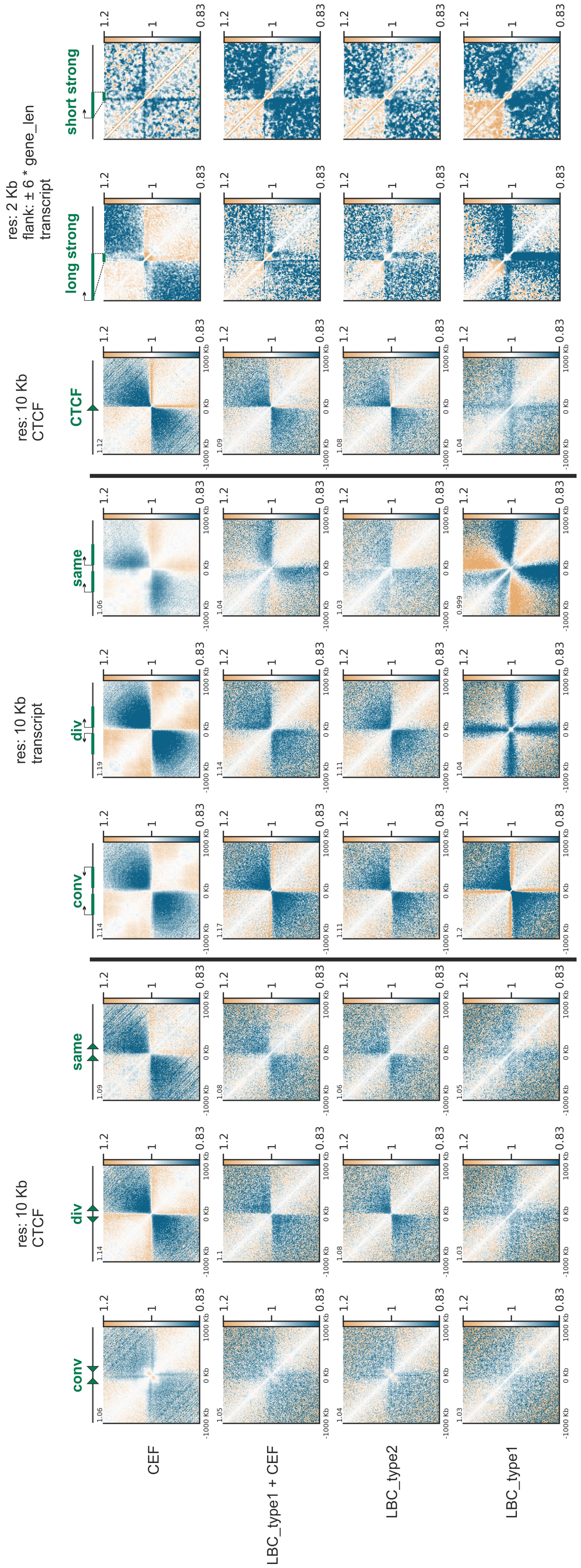


**Figure S16. The Hi-C maps averaged near different genes or CTCF sites orientations for the lampbrush chromosome (LBC) stage oocytes of type 1 and 2, chicken embryonic fibroblasts (CEF) and the *in silico* mix of the LBC stage oocytes of type 1 with CEF.**

**Figure S17. Examples of the gene pair orientation in borders mistyping.**

All borders in Hi-C map were assigned to the closest gene pair orientation type. The closest gene pair was chosen by measuring the distance between the exact border coordinate and the middle point inside the gene pair. Note that, the genes with length less than 5 Kb and FPKM <= 0 were excluded from the couples formation. Additionally, one should note that the error in defining domain boundaries is greater than 16 Kb for some domains. All figures contains the red border tracks (the red line denotes ±16 Kb from the border coordinate) for each of four possible orientations: Forward-Reverse (FR, convergent), Forward-Forward (FF, co-directional), Reverse-Forward (RF, divergent) and Reverse-Reverse (RR, co-directional). The histograms denotes the total stranded RNA-seq data for lampbrush-stage oocyte nuclei (15), the “valid_transripts.gtf” file contains the annotation of transcription units based on de novo transcriptome analysis with one alternative transcript with the highest FPKM chosen for each gene (see Methods).

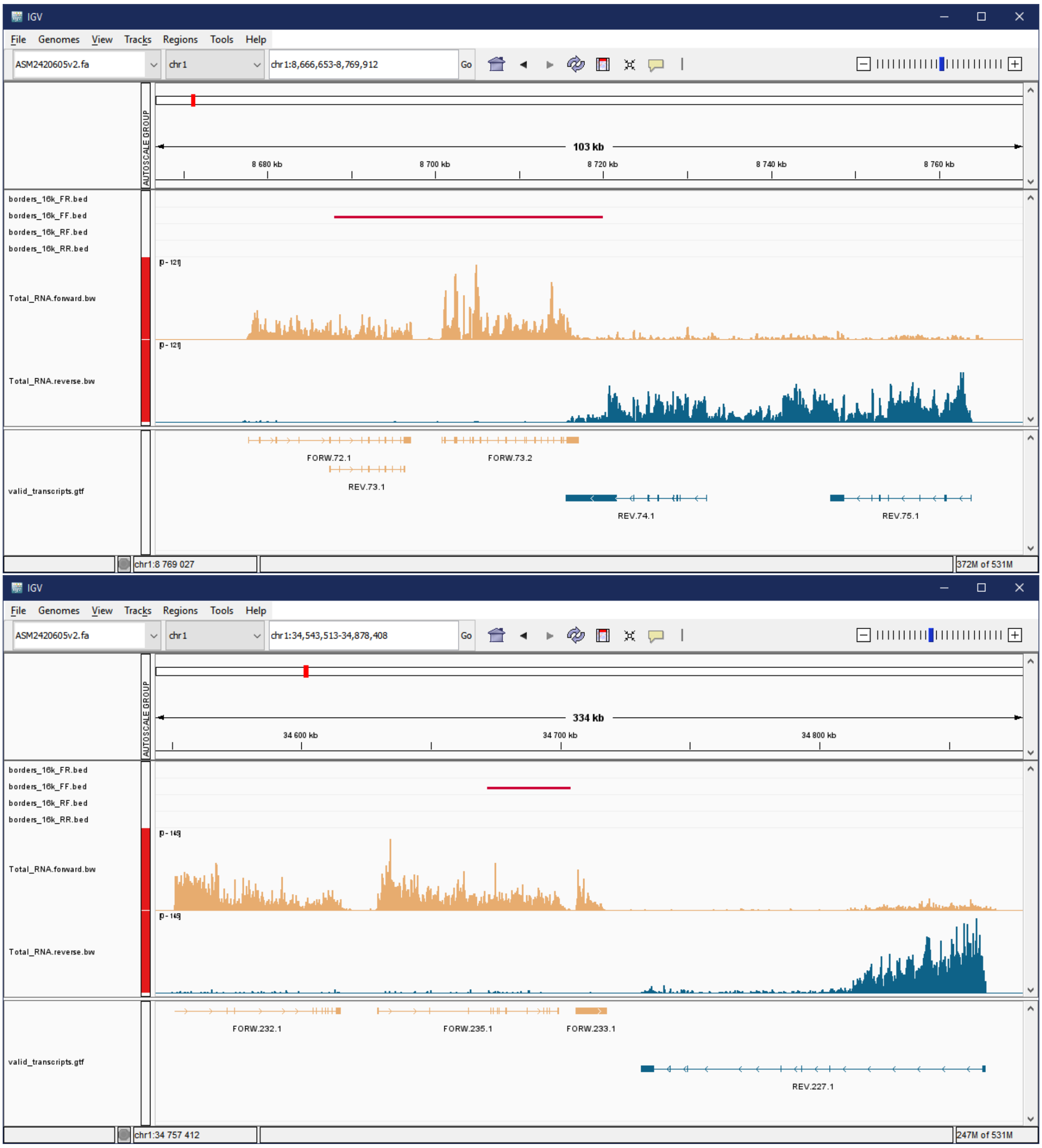

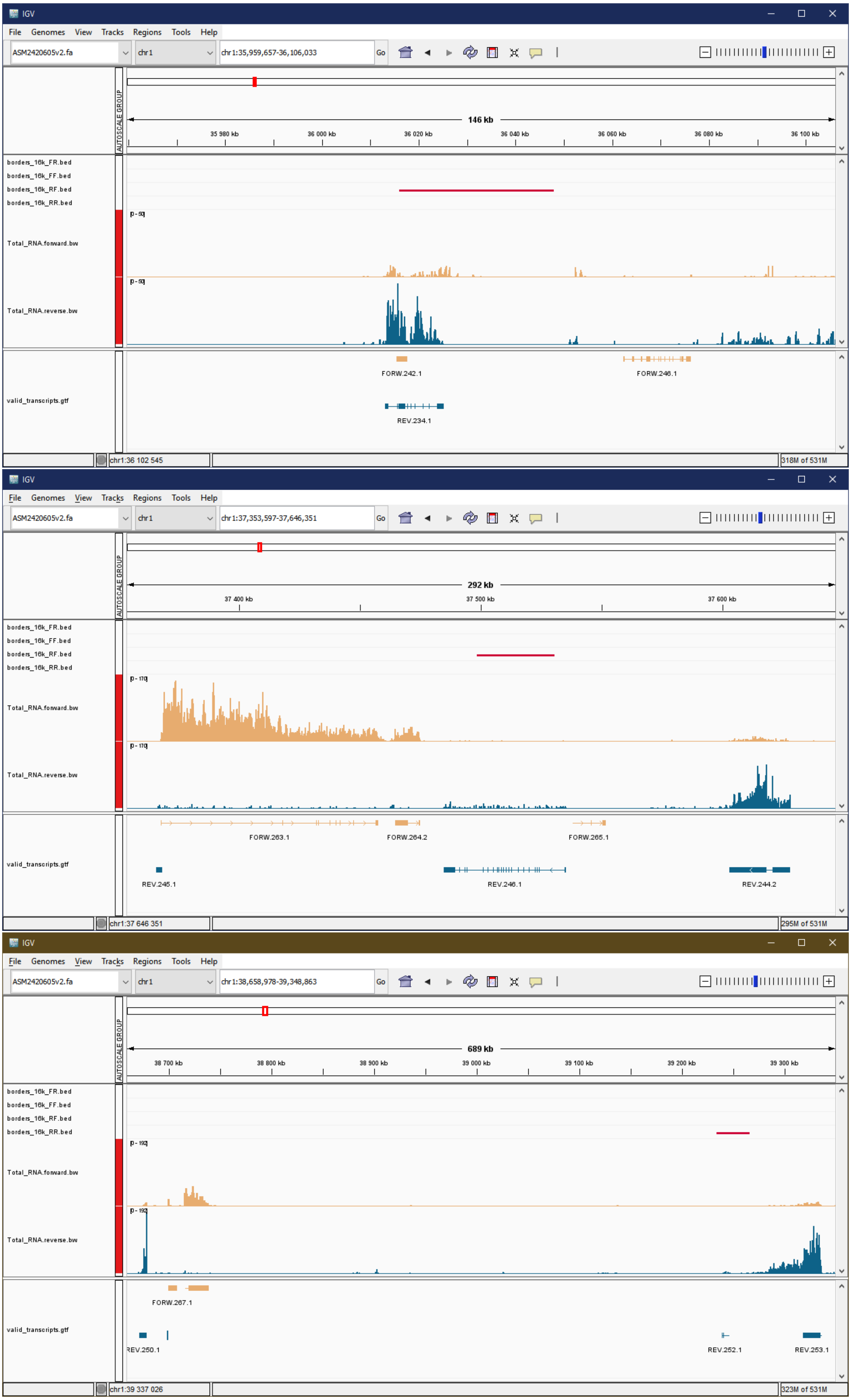
IGV
File Genomes View Tracks Regions Tools Help
ASM2420605v2.fa
chr1
chr1:35,959,657-36,106,033
Go
AUTOSCALE GROUP
146 kb
35 980 kb
36 000 kb
36 020 kb
36 040 kb
36 060 kb
36 080 kb
36 100 kb
borders_16k_FR.bed
borders_16k_FF.bed
borders_16k_RF.bed
borders_16k_RR.bed
Total_RNA.forward.bw
Total_RNA.reverse.bw
valid_transcripts.gtf
FORW.242.1
FORW.246.1
REV.234.1
chr1:36 102 545
318M of 531M
IGV
File Genomes View Tracks Regions Tools Help
ASM2420605v2.fa
chr1
chr1:37,353,597-37,646,351
Go
AUTOSCALE GROUP
292 kb
37 400 kb
37 500 kb
37 600 kb
borders_16k_FR.bed
borders_16k_FF.bed
borders_16k_RF.bed
borders_16k_RR.bed
Total_RNA.forward.bw
Total_RNA.reverse.bw
valid_transcripts.gtf
FORW.263.1
FORW.264.2
FORW.265.1
REV.245.1
REV.246.1
REV.244.2
chr1:37 646 351
295M of 531M
IGV
File Genomes View Tracks Regions Tools Help
ASM2420605v2.fa
chr1
chr1:38,658,978-39,348,863
Go
AUTOSCALE GROUP
689 kb
38 700 kb
38 800 kb
38 900 kb
39 000 kb
39 100 kb
39 200 kb
39 300 kb
borders_16k_FR.bed
borders_16k_FF.bed
borders_16k_RF.bed
borders_16k_RR.bed
Total_RNA.forward.bw
Total_RNA.reverse.bw
valid_transcripts.gtf
FORW.267.1
REV.250.1
REV.252.1
REV.253.1
chr1:39 337 026
323M of 531M

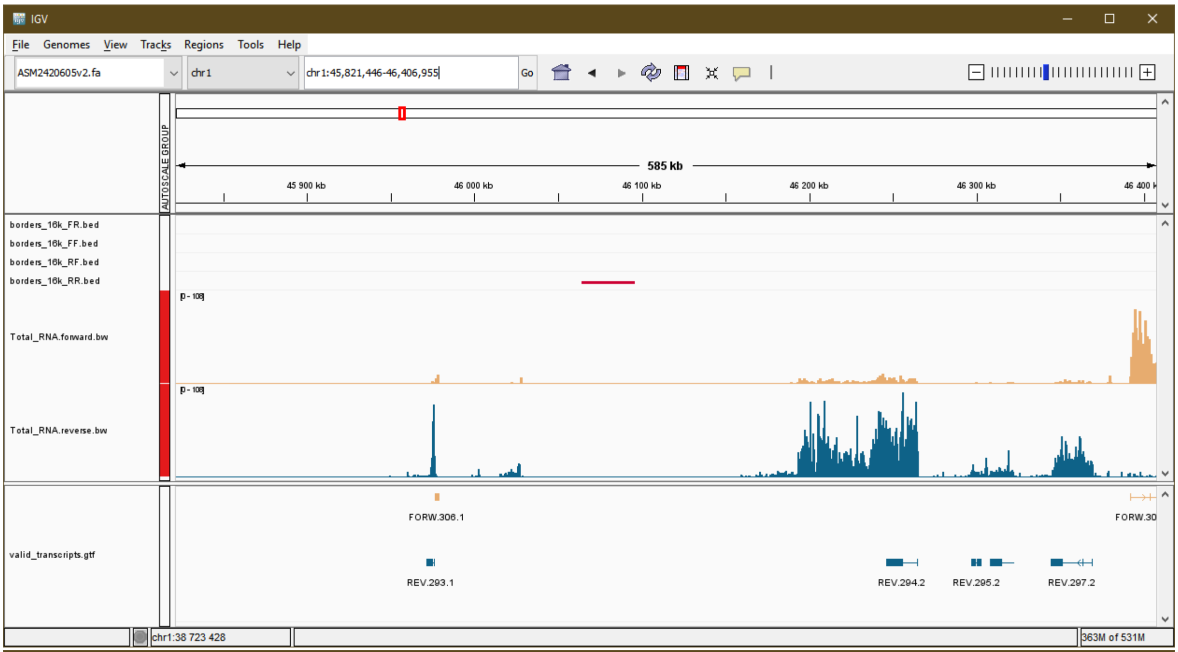
IGV
File Genomes View Tracks Regions Tools Help
ASM2420605v2.fa
chr1
chr1:45,821,446-46,406,955
Go
AUTOSCALE GROUP
585 kb
45 900 kb
46 000 kb
46 100 kb
46 200 kb
46 300 kb
borders_16k_FR.bed
borders_16k_FF.bed
borders_16k_RF.bed
borders_16k_RR.bed
Total_RNA.forward.bw
Total_RNA.reverse.bw
valid_transcripts.gtf
FORW.306.1
REV.293.1
REV.294.2
REV.295.2
REV.297.2
chr1:38 723 428
363M of 531M

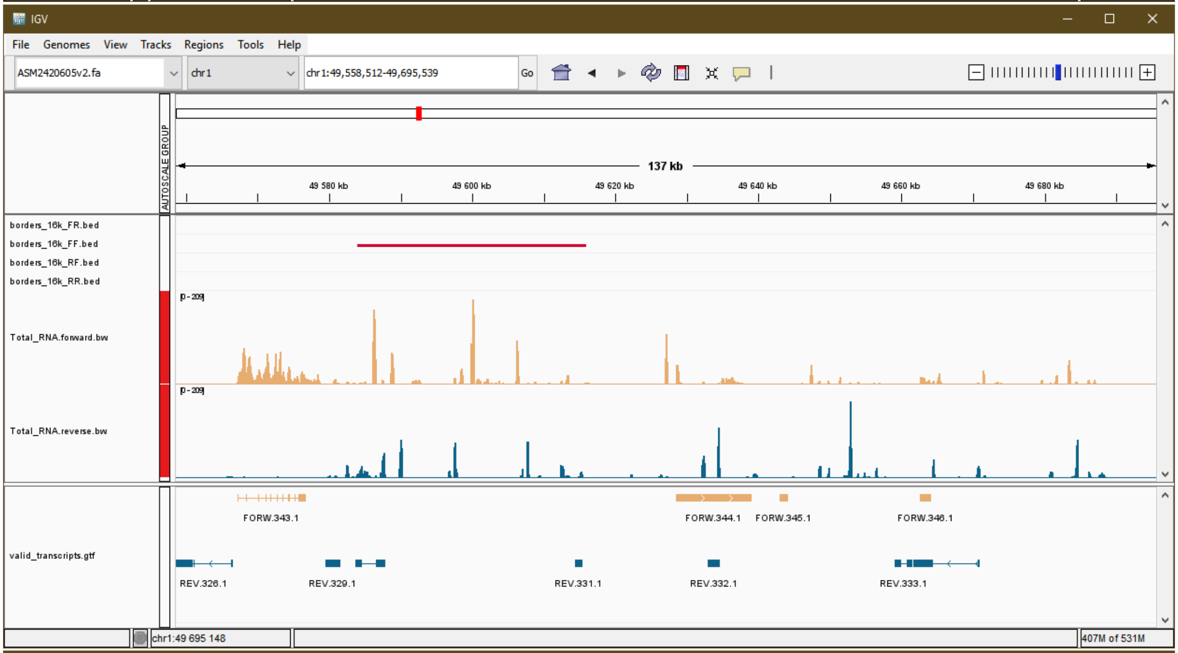
IGV
File Genomes View Tracks Regions Tools Help
ASM2420605v2.fa
chr1
chr1:49,558,512-49,695,539
Go
AUTOSCALE GROUP
137 kb
49 580 kb
49 600 kb
49 620 kb
49 640 kb
49 660 kb
49 680 kb
borders_16k_FR.bed
borders_16k_FF.bed
borders_16k_RF.bed
borders_16k_RR.bed
Total_RNA.forward.bw
Total_RNA.reverse.bw
valid_transcripts.gtf
FORW.343.1
FORW.344.1
FORW.345.1
FORW.346.1
REV.326.1
REV.329.1
REV.331.1
REV.332.1
REV.333.1
chr1:49 695 148
407M of 531M

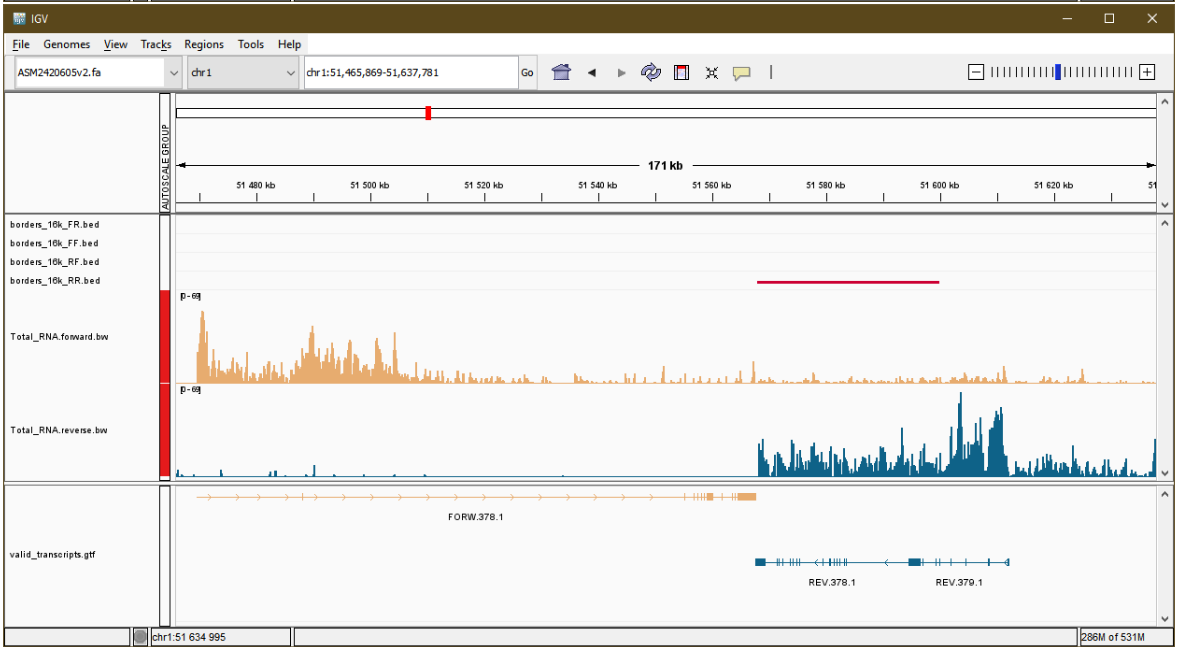
IGV
File Genomes View Tracks Regions Tools Help
ASM2420605v2.fa
chr1
chr1:51,465,869-51,637,781
Go
AUTOSCALE GROUP
171 kb
51 480 kb
51 500 kb
51 520 kb
51 540 kb
51 560 kb
51 580 kb
51 600 kb
51 620 kb
borders_16k_FR.bed
borders_16k_FF.bed
borders_16k_RF.bed
borders_16k_RR.bed
Total_RNA.forward.bw
Total_RNA.reverse.bw
valid_transcripts.gtf
FORW.378.1
REV.378.1
REV.379.1
chr1:51 634 995
286M of 531M

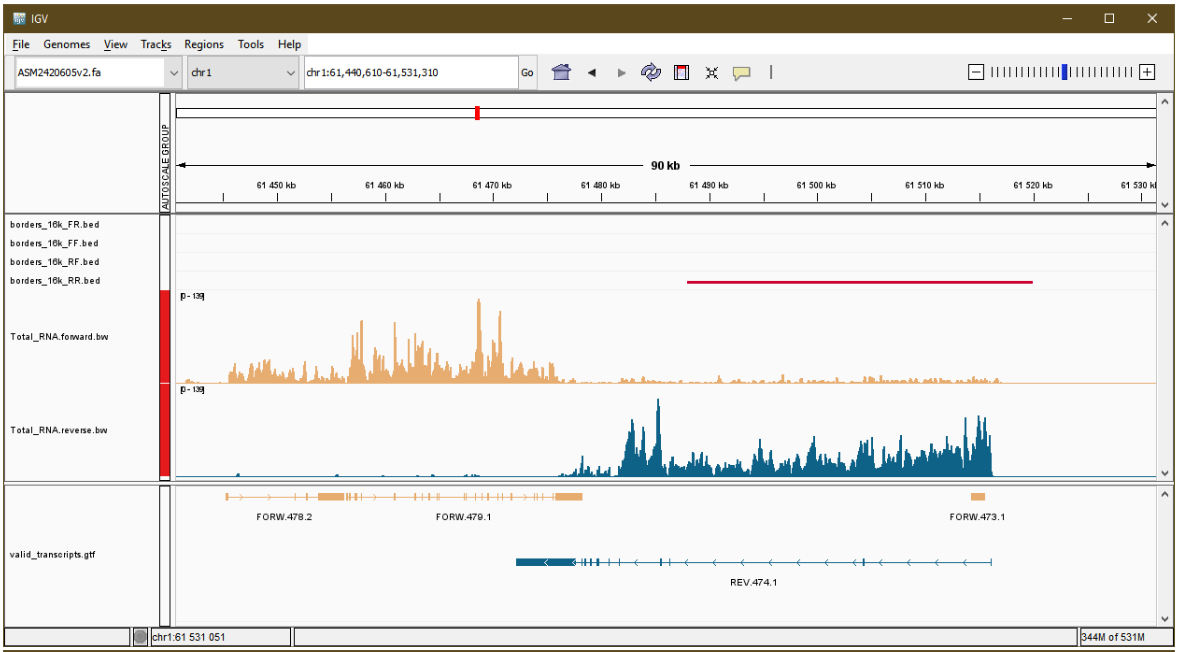
IGV
File Genomes View Tracks Regions Tools Help
ASM2420605v2.fa
chr1
chr1:61,440,610-61,531,310
Go
AUTOSCALE GROUP
90 kb
61 450 kb
61 460 kb
61 470 kb
61 480 kb
61 490 kb
61 500 kb
61 510 kb
61 520 kb
61 530 kb
borders_16k_FR.bed
borders_16k_FF.bed
borders_16k_RF.bed
borders_16k_RR.bed
Total_RNA.forward.bw
Total_RNA.reverse.bw
valid_transcripts.gtf
FORW.478.2
FORW.479.1
FORW.473.1
REV.474.1
chr1:61 531 051
344M of 531M

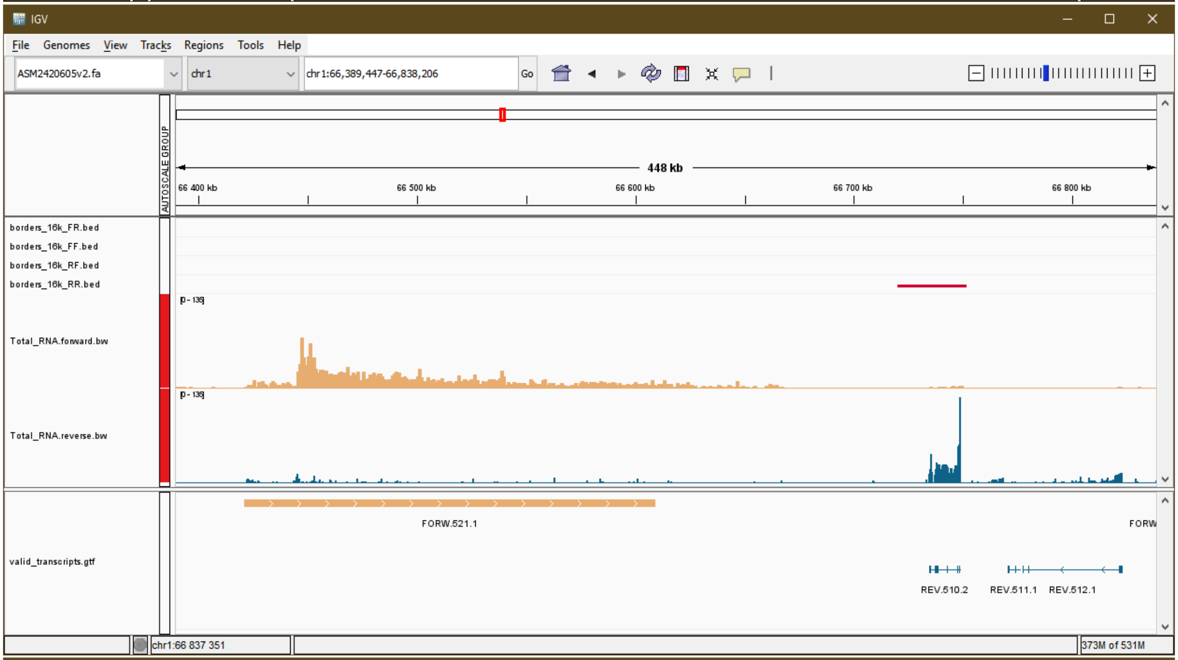
IGV
File Genomes View Tracks Regions Tools Help
ASM2420605v2.fa
chr1
chr1:66,389,447-66,838,206
Go
AUTOSCALE GROUP
448 kb
66 400 kb
66 500 kb
66 600 kb
66 700 kb
66 800 kb
borders_16k_FR.bed
borders_16k_FF.bed
borders_16k_RF.bed
borders_16k_RR.bed
Total_RNA.forward.bw
Total_RNA.reverse.bw
valid_transcripts.gtf
FORW.521.1
FORW
REV.510.2
REV.511.1
REV.512.1
chr1:66 837 351
373M of 531M

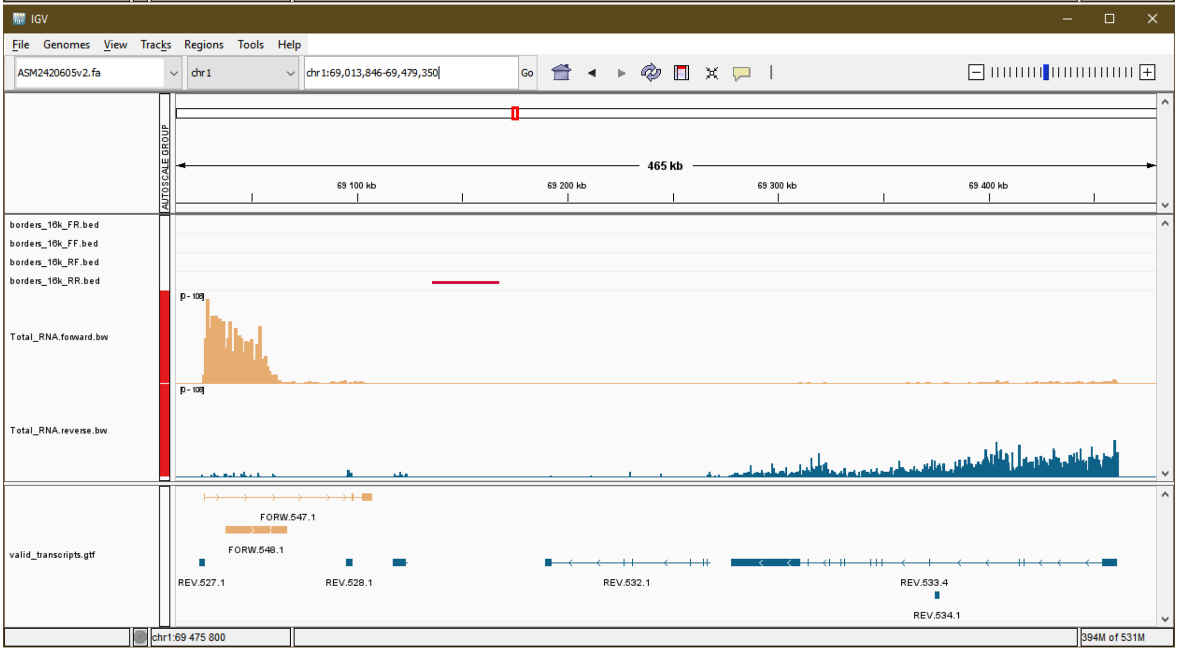
IGV
File Genomes View Tracks Regions Tools Help
ASM2420605v2.fa
chr1
chr1:69,013,846-69,479,350
Go
AUTOSCALE GROUP
465 kb
69 100 kb
69 200 kb
69 300 kb
69 400 kb
borders_16k_FR.bed
borders_16k_FF.bed
borders_16k_RF.bed
borders_16k_RR.bed
Total_RNA.forward.bw
Total_RNA.reverse.bw
valid_transcripts.gtf
FORW.547.1
FORW.548.1
REV.527.1
REV.528.1
REV.532.1
REV.533.4
REV.534.1
chr1:69 475 800
394M of 531M

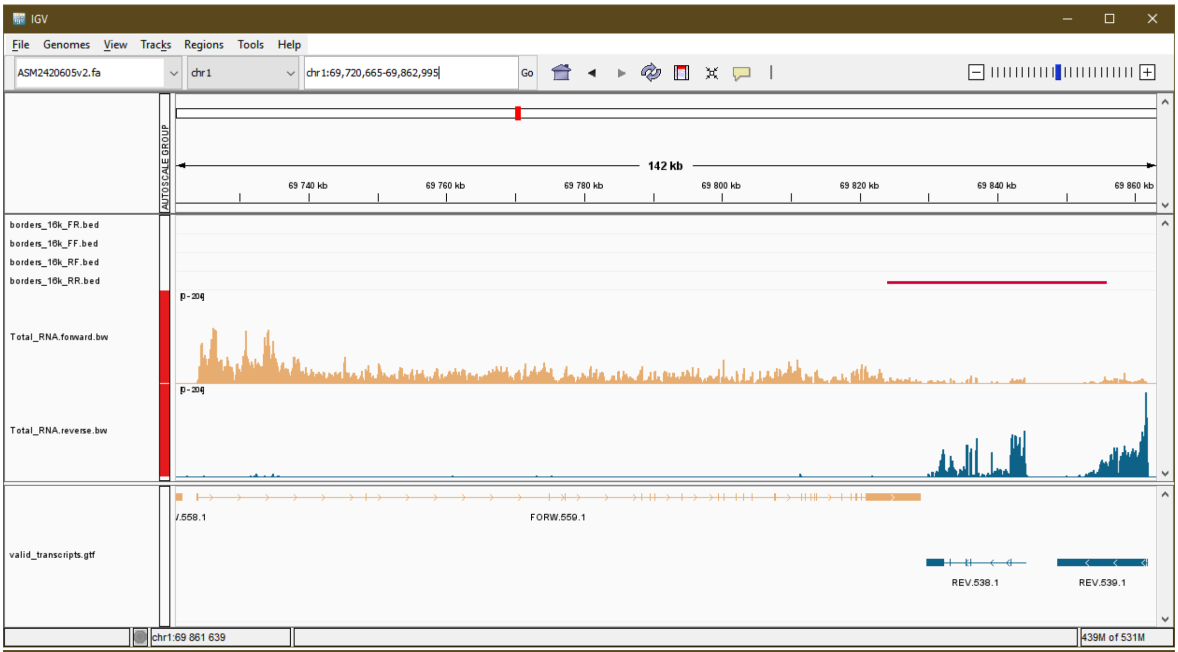
IGV
File Genomes View Tracks Regions Tools Help
ASM2420605v2.fa
chr1
chr1:69,720,665-69,862,995
Go
AUTOSCALE GROUP
142 kb
69 740 kb
69 760 kb
69 780 kb
69 800 kb
69 820 kb
69 840 kb
69 860 kb
borders_16k_FR.bed
borders_16k_FF.bed
borders_16k_RF.bed
borders_16k_RR.bed
Total_RNA.forward.bw
Total_RNA.reverse.bw
valid_transcripts.gtf
FORW.559.1
REV.538.1
REV.539.1
chr1:69 861 639
439M of 531M

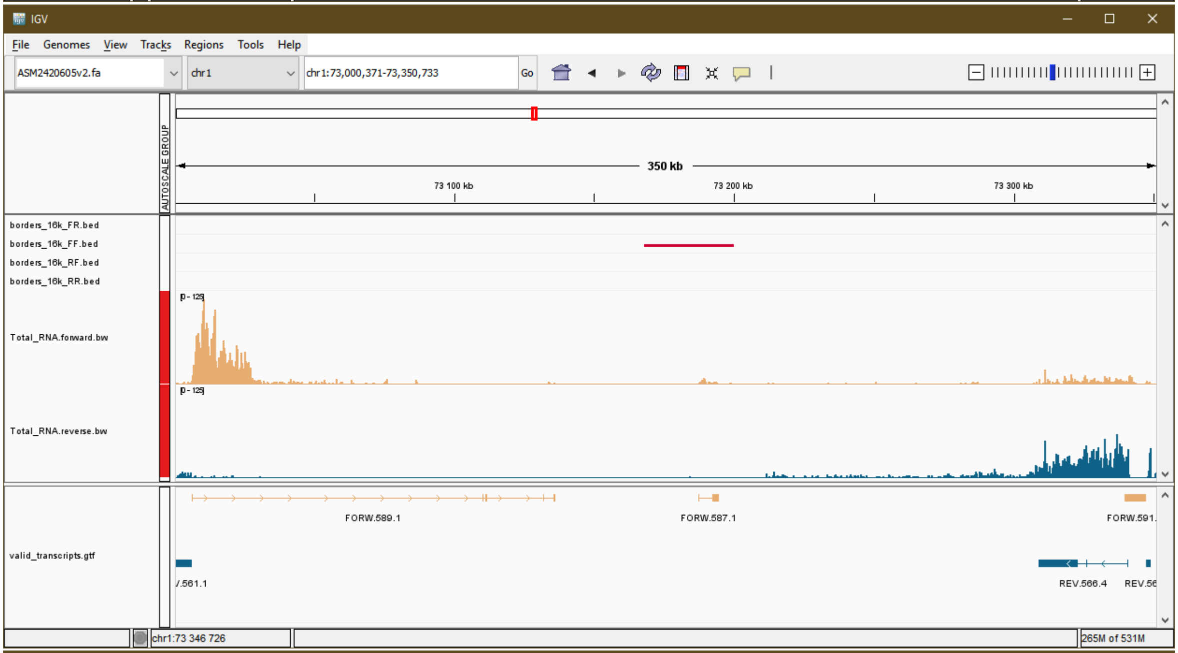
IGV
File Genomes View Tracks Regions Tools Help
ASM2420605v2.fa
chr1
chr1:73,000,371-73,350,733
Go
AUTOSCALE GROUP
350 kb
73 100 kb
73 200 kb
73 300 kb
borders_16k_FR.bed
borders_16k_FF.bed
borders_16k_RF.bed
borders_16k_RR.bed
Total_RNA.forward.bw
Total_RNA.reverse.bw
valid_transcripts.gtf
FORW.589.1
FORW.587.1
REV.566.4
chr1:73 346 726
265M of 531M

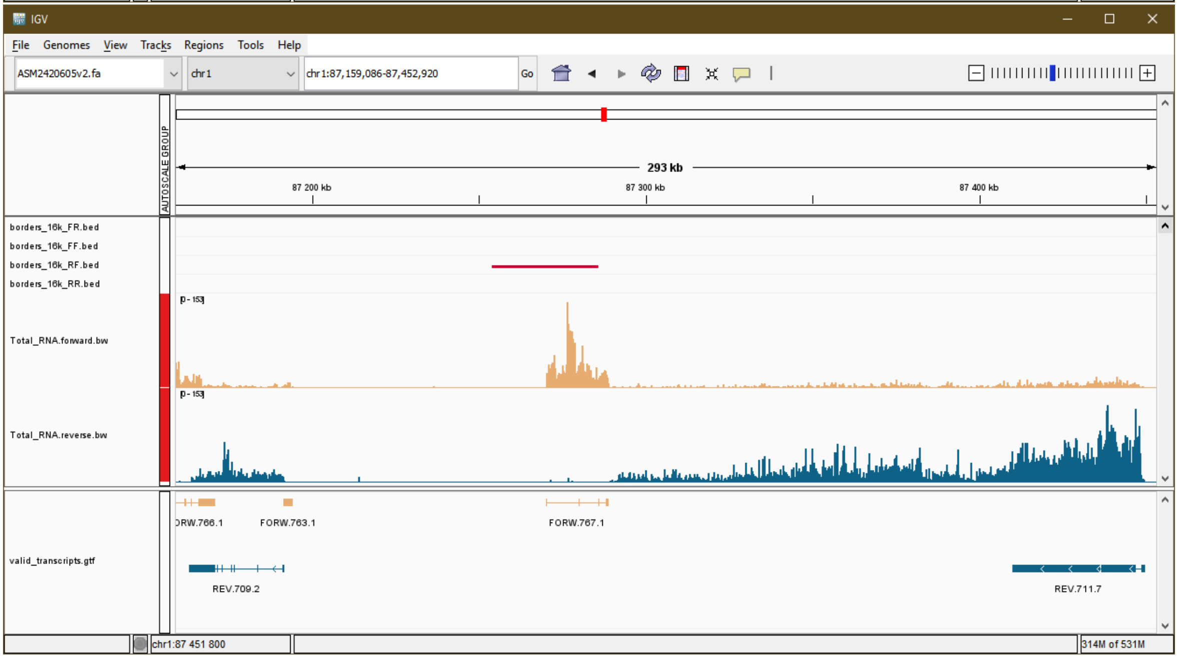
IGV
File Genomes View Tracks Regions Tools Help
ASM2420605v2.fa
chr1
chr1:87,159,086-87,452,920
Go
AUTOSCALE GROUP
293 kb
87 200 kb
87 300 kb
87 400 kb
borders_16k_FR.bed
borders_16k_FF.bed
borders_16k_RF.bed
borders_16k_RR.bed
Total_RNA.forward.bw
Total_RNA.reverse.bw
valid_transcripts.gtf
FORW.763.1
FORW.767.1
REV.709.2
REV.711.7
chr1:87 451 800
314M of 531M

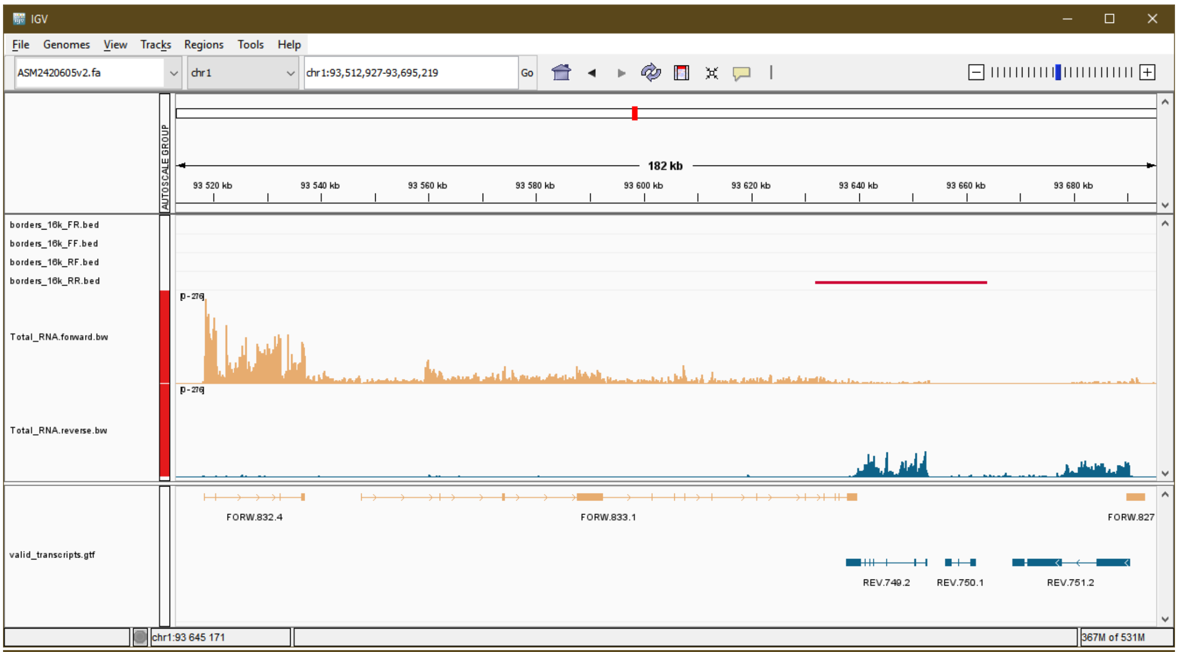
IGV
File Genomes View Tracks Regions Tools Help
ASM2420605v2.fa
chr1
chr1:93,512,927-93,695,219
Go
182 kb
93 520 kb
93 540 kb
93 560 kb
93 580 kb
93 600 kb
93 620 kb
93 640 kb
93 660 kb
93 680 kb
AUTOSCALE GROUP
borders_16k_FR.bed
borders_16k_FF.bed
borders_16k_RF.bed
borders_16k_RR.bed
Total_RNA.forward.bw
Total_RNA.reverse.bw
valid_transcripts.gtf
FORW.832.4
FORW.833.1
FORW.827
REV.749.2
REV.750.1
REV.751.2
chr1:93 645 171
367M of 531M

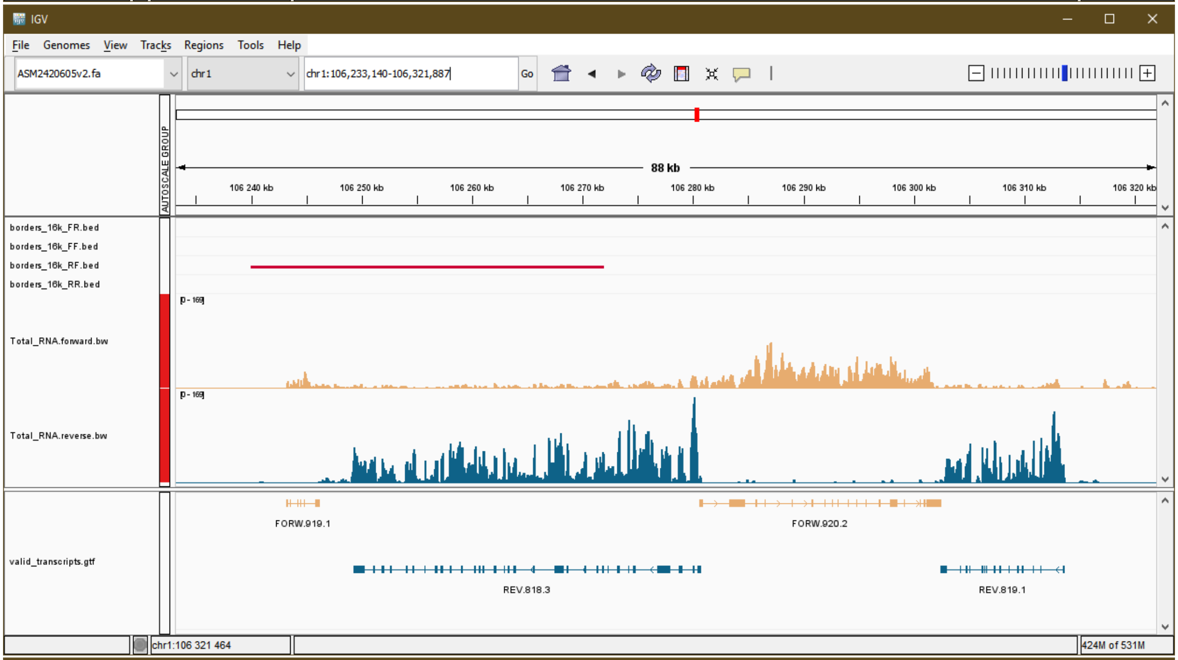
IGV
File Genomes View Tracks Regions Tools Help
ASM2420605v2.fa
chr1
chr1:106,233,140-106,321,887
Go
88 kb
106 240 kb
106 250 kb
106 260 kb
106 270 kb
106 280 kb
106 290 kb
106 300 kb
106 310 kb
106 320 kb
AUTOSCALE GROUP
borders_16k_FR.bed
borders_16k_FF.bed
borders_16k_RF.bed
borders_16k_RR.bed
Total_RNA.forward.bw
Total_RNA.reverse.bw
valid_transcripts.gtf
FORW.919.1
FORW.920.2
REV.818.3
REV.819.1
chr1:106 321 464
424M of 531M

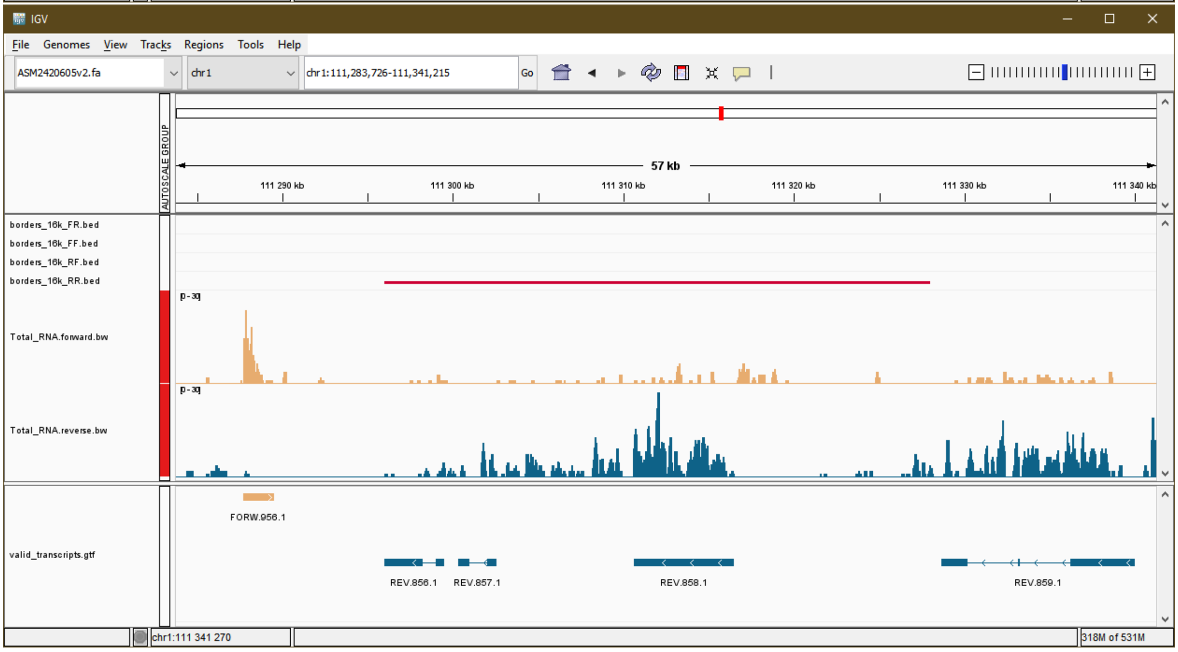
IGV
File Genomes View Tracks Regions Tools Help
ASM2420605v2.fa
chr1
chr1:111,283,726-111,341,215
Go
57 kb
111 290 kb
111 300 kb
111 310 kb
111 320 kb
111 330 kb
111 340 kb
AUTOSCALE GROUP
borders_16k_FR.bed
borders_16k_FF.bed
borders_16k_RF.bed
borders_16k_RR.bed
Total_RNA.forward.bw
Total_RNA.reverse.bw
valid_transcripts.gtf
FORW.956.1
REV.856.1
REV.857.1
REV.858.1
REV.859.1
chr1:111 341 270
318M of 531M

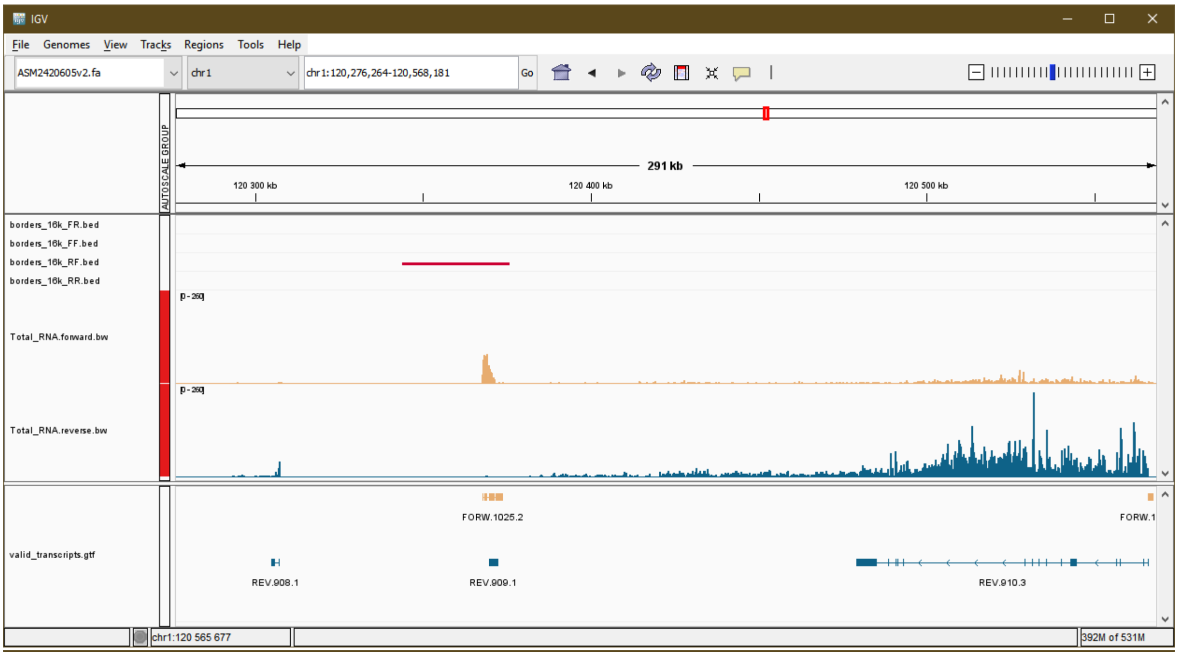
IGV
File Genomes View Tracks Regions Tools Help
ASM2420605v2.fa
chr1
chr1:120,276,264-120,568,181
Go
AUTOSCALE GROUP
291 kb
120 300 kb
120 400 kb
120 500 kb
borders_16k_FR.bed
borders_16k_FF.bed
borders_16k_RF.bed
borders_16k_RR.bed
Total_RNA.forward.bw
Total_RNA.reverse.bw
valid_transcripts.gtf
FORW.1025.2
FORW.1
REV.908.1
REV.909.1
REV.910.3
chr1:120 565 677
392M of 531M

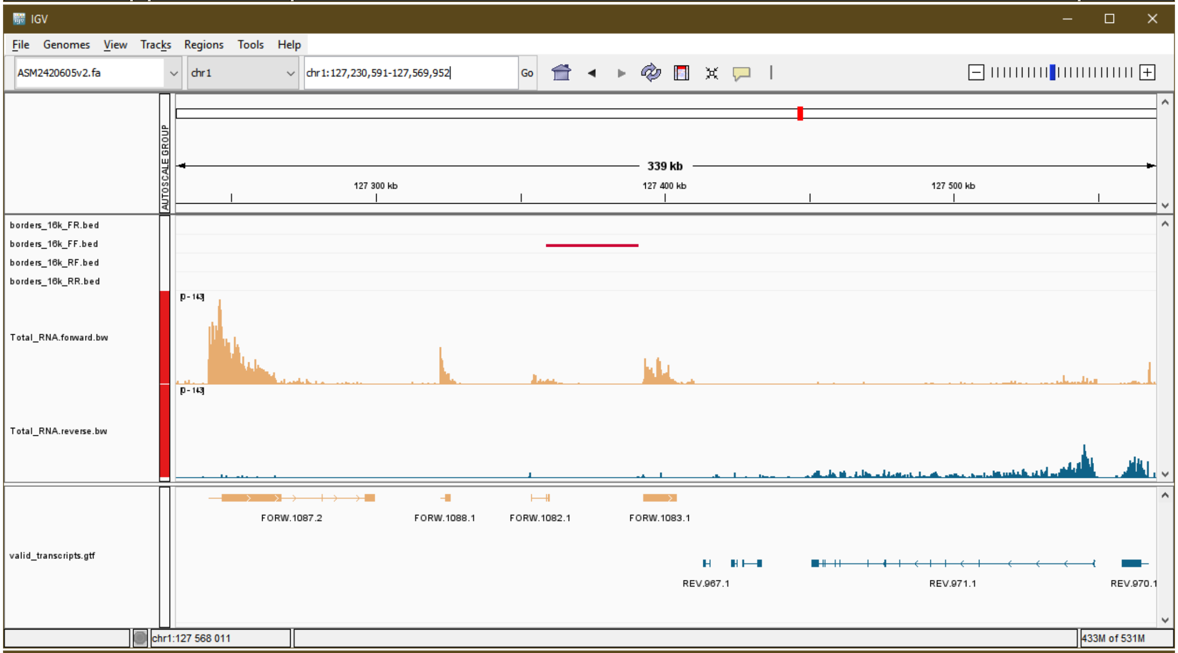
IGV
File Genomes View Tracks Regions Tools Help
ASM2420605v2.fa
chr1
chr1:127,230,591-127,569,952
Go
AUTOSCALE GROUP
339 kb
127 300 kb
127 400 kb
127 500 kb
borders_16k_FR.bed
borders_16k_FF.bed
borders_16k_RF.bed
borders_16k_RR.bed
Total_RNA.forward.bw
Total_RNA.reverse.bw
valid_transcripts.gtf
FORW.1087.2
FORW.1088.1
FORW.1082.1
FORW.1083.1
REV.967.1
REV.971.1
REV.970.1
chr1:127 568 011
433M of 531M

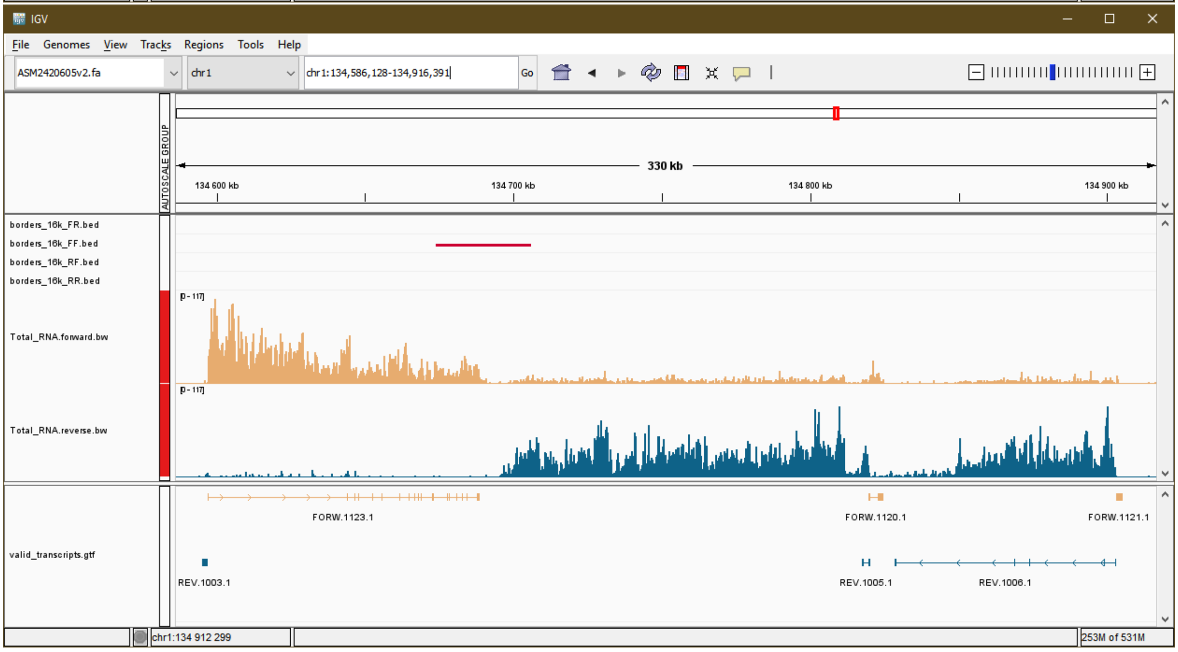
IGV
File Genomes View Tracks Regions Tools Help
ASM2420605v2.fa
chr1
chr1:134,586,128-134,916,391
Go
AUTOSCALE GROUP
330 kb
134 600 kb
134 700 kb
134 800 kb
134 900 kb
borders_16k_FR.bed
borders_16k_FF.bed
borders_16k_RF.bed
borders_16k_RR.bed
Total_RNA.forward.bw
Total_RNA.reverse.bw
valid_transcripts.gtf
FORW.1123.1
FORW.1120.1
FORW.1121.1
REV.1003.1
REV.1005.1
REV.1006.1
chr1:134 912 299
253M of 531M

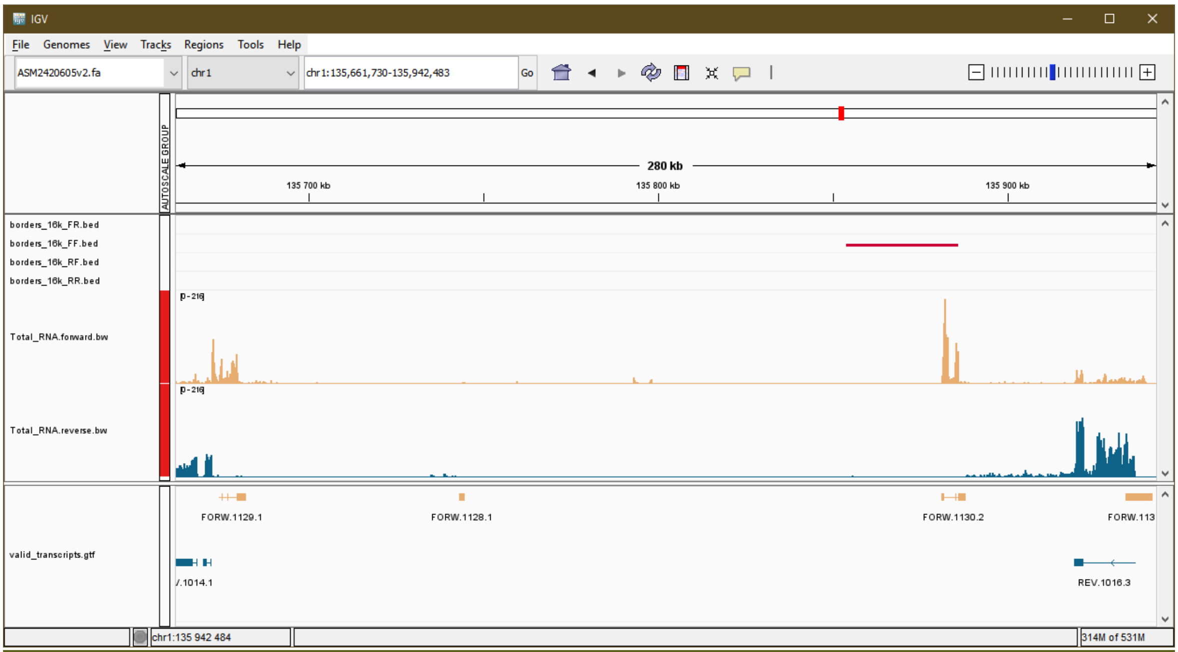
IGV
File Genomes View Tracks Regions Tools Help
ASM2420605v2.fa
chr1
chr1:135,661,730-135,942,483
Go
AUTOSCALE GROUP
280 kb
135 700 kb
135 800 kb
135 900 kb
borders_16k_FR.bed
borders_16k_FF.bed
borders_16k_RF.bed
borders_16k_RR.bed
Total_RNA.forward.bw
Total_RNA.reverse.bw
valid_transcripts.gtf
FORW.1129.1
FORW.1128.1
FORW.1130.2
FORW.113
/.1014.1
REV.1016.3
chr1:135 942 484
314M of 531M

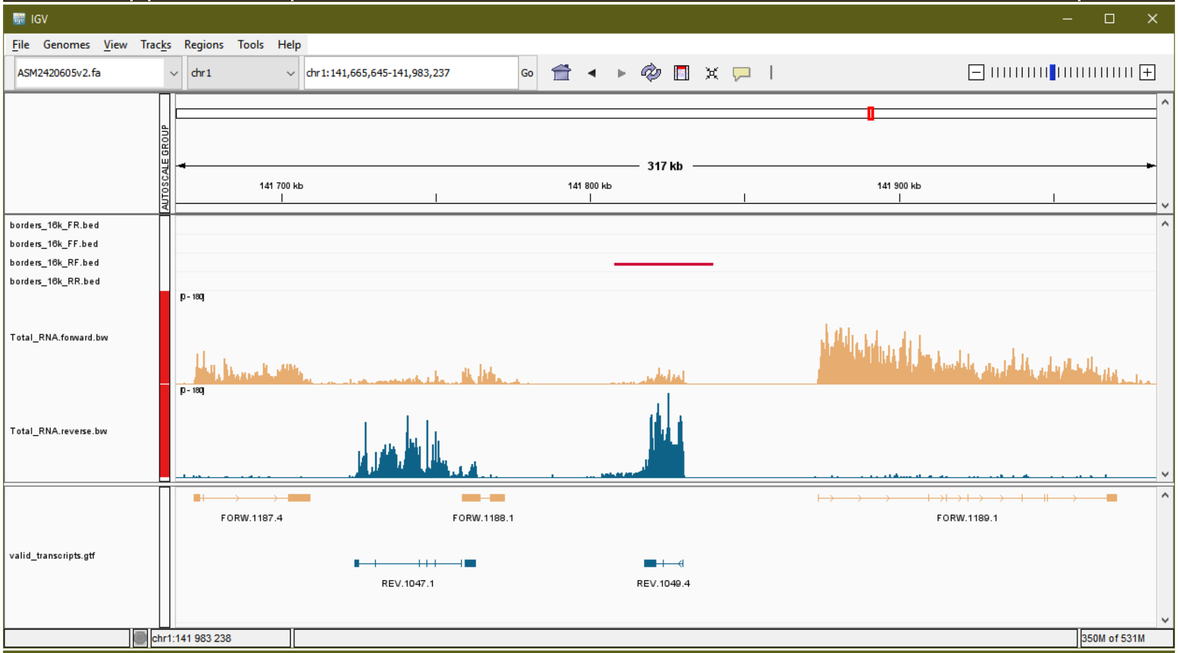
IGV
File Genomes View Tracks Regions Tools Help
ASM2420605v2.fa
chr1
chr1:141,665,645-141,983,237
Go
AUTOSCALE GROUP
317 kb
141 700 kb
141 800 kb
141 900 kb
borders_16k_FR.bed
borders_16k_FF.bed
borders_16k_RF.bed
borders_16k_RR.bed
Total_RNA.forward.bw
Total_RNA.reverse.bw
valid_transcripts.gtf
FORW.1187.4
FORW.1188.1
FORW.1189.1
REV.1047.1
REV.1049.4
chr1:141 983 238
350M of 531M

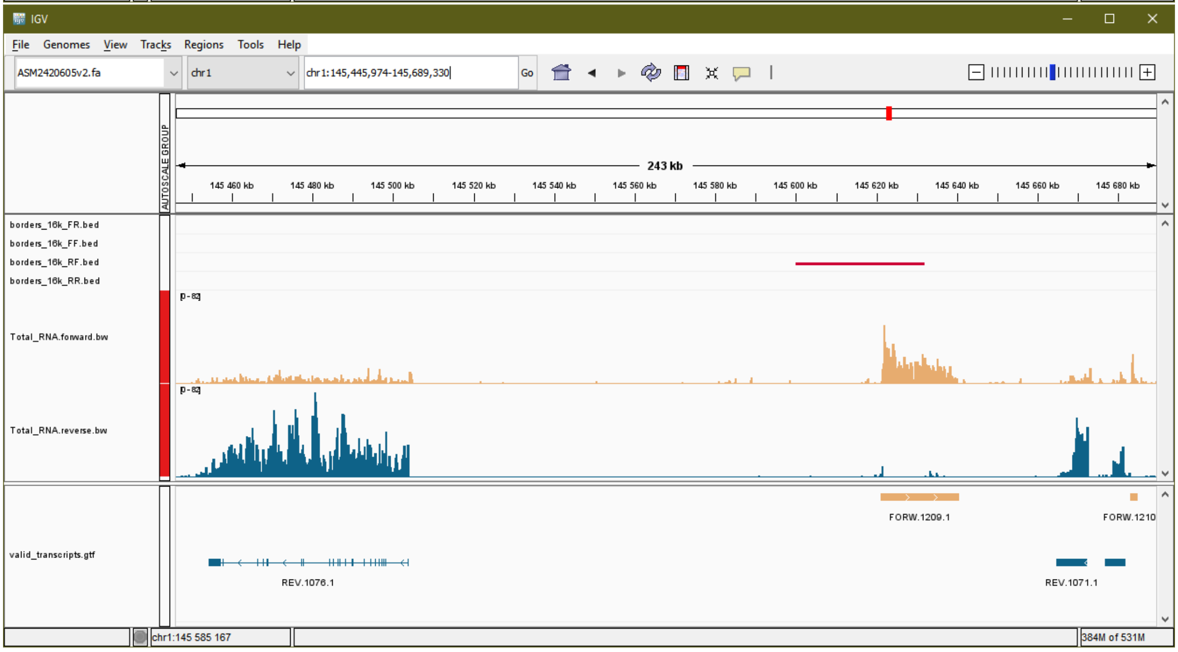
IGV
File Genomes View Tracks Regions Tools Help
ASM2420605v2.fa
chr1
chr1:145,445,974-145,689,330
Go
AUTOSCALE GROUP
243 kb
145 460 kb
145 480 kb
145 500 kb
145 520 kb
145 540 kb
145 560 kb
145 580 kb
145 600 kb
145 620 kb
145 640 kb
145 660 kb
145 680 kb
borders_16k_FR.bed
borders_16k_FF.bed
borders_16k_RF.bed
borders_16k_RR.bed
Total_RNA.forward.bw
Total_RNA.reverse.bw
valid_transcripts.gtf
FORW.1209.1
FORW.1210
REV.1076.1
REV.1071.1
chr1:145 585 167
384M of 531M

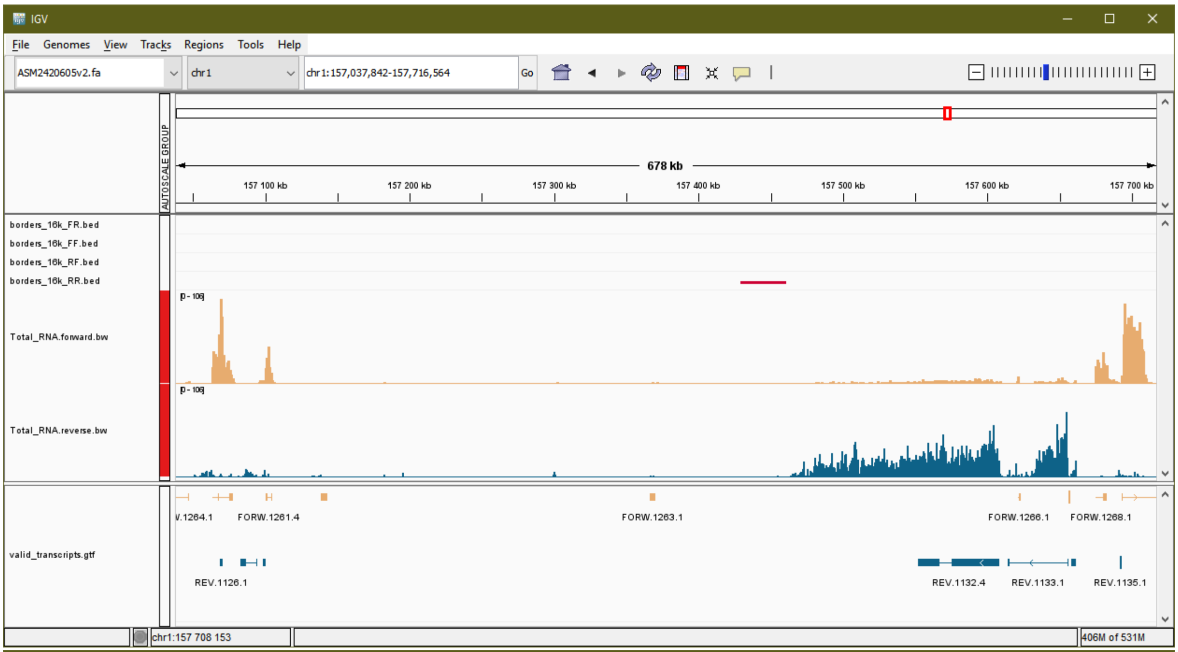
IGV
File Genomes View Tracks Regions Tools Help
ASM2420605v2.fa
chr1
chr1:157,037,842-157,716,564
Go
678 kb
157 100 kb
157 200 kb
157 300 kb
157 400 kb
157 500 kb
157 600 kb
157 700 kb
AUTOSCALE GROUP
borders_16k_FR.bed
borders_16k_FF.bed
borders_16k_RF.bed
borders_16k_RR.bed
Total_RNA.forward.bw
Total_RNA.reverse.bw
[0 - 106]
valid_transcripts.gtf
V.1264.1
FORW.1261.4
FORW.1263.1
FORW.1266.1
FORW.1268.1
REV.1126.1
REV.1132.4
REV.1133.1
REV.1135.1
chr1:157 708 153
406M of 531M

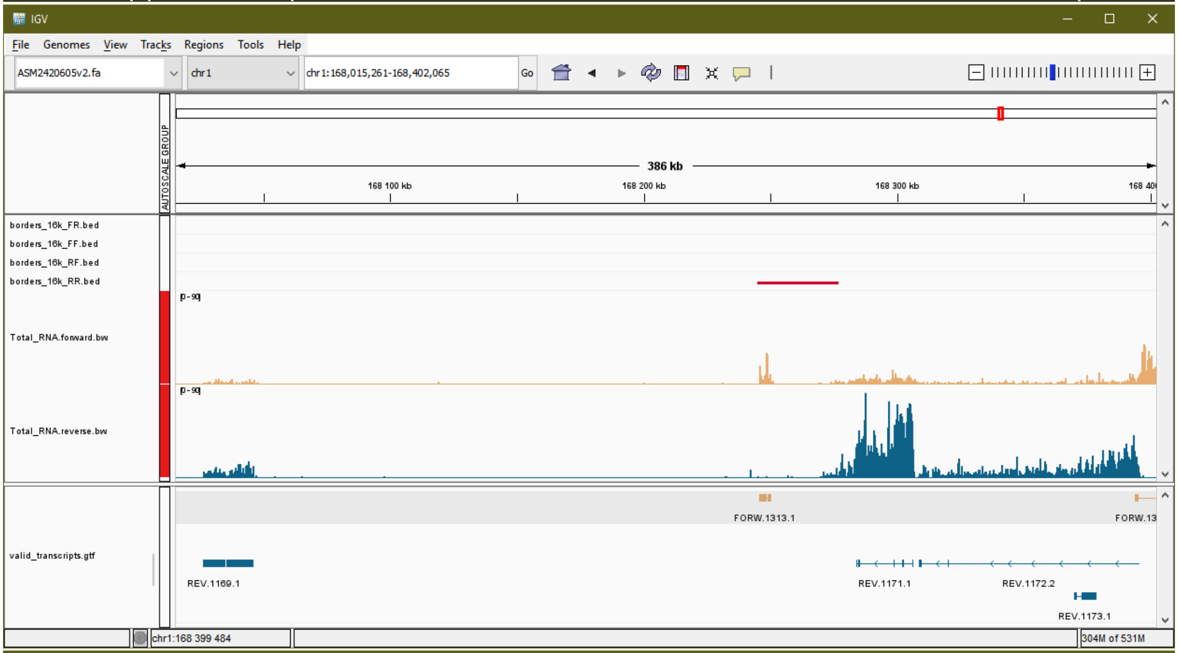
IGV
File Genomes View Tracks Regions Tools Help
ASM2420605v2.fa
chr1
chr1:168,015,261-168,402,065
Go
386 kb
168 100 kb
168 200 kb
168 300 kb
AUTOSCALE GROUP
borders_16k_FR.bed
borders_16k_FF.bed
borders_16k_RF.bed
borders_16k_RR.bed
Total_RNA.forward.bw
Total_RNA.reverse.bw
[0 - 90]
valid_transcripts.gtf
FORW.1313.1
REV.1169.1
REV.1171.1
REV.1172.2
REV.1173.1
chr1:168 399 484
304M of 531M

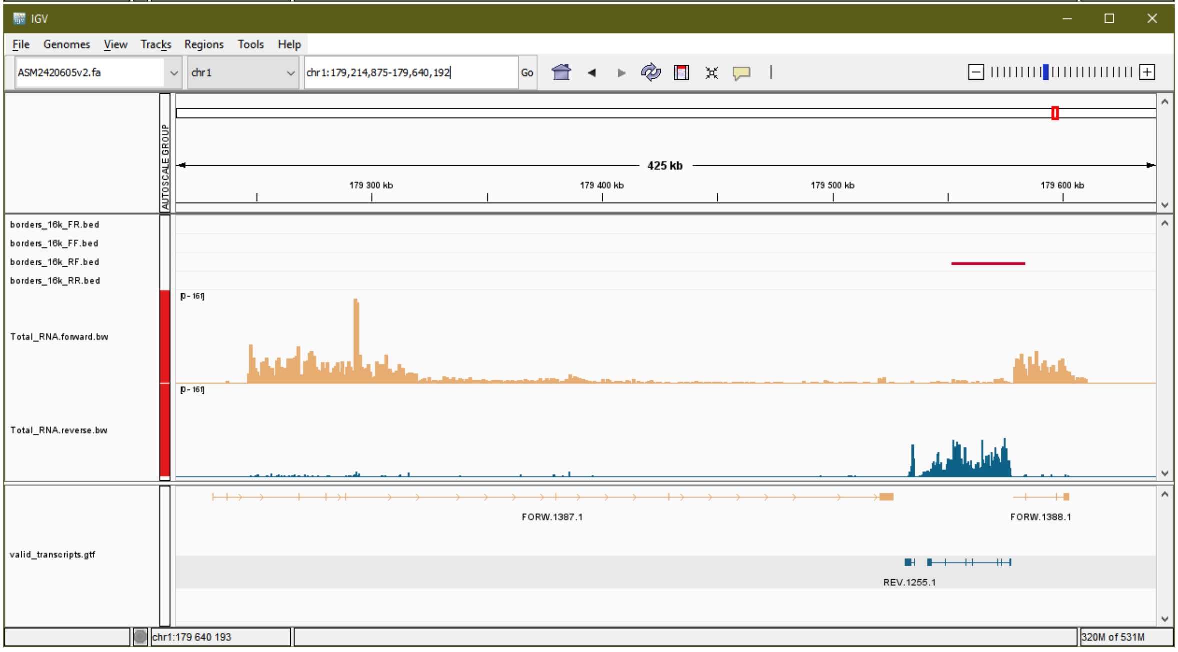
IGV
File Genomes View Tracks Regions Tools Help
ASM2420605v2.fa
chr1
chr1:179,214,875-179,640,192
Go
425 kb
179 300 kb
179 400 kb
179 500 kb
179 600 kb
AUTOSCALE GROUP
borders_16k_FR.bed
borders_16k_FF.bed
borders_16k_RF.bed
borders_16k_RR.bed
Total_RNA.forward.bw
Total_RNA.reverse.bw
[0 - 161]
valid_transcripts.gtf
FORW.1387.1
FORW.1388.1
REV.1255.1
chr1:179 640 193
320M of 531M

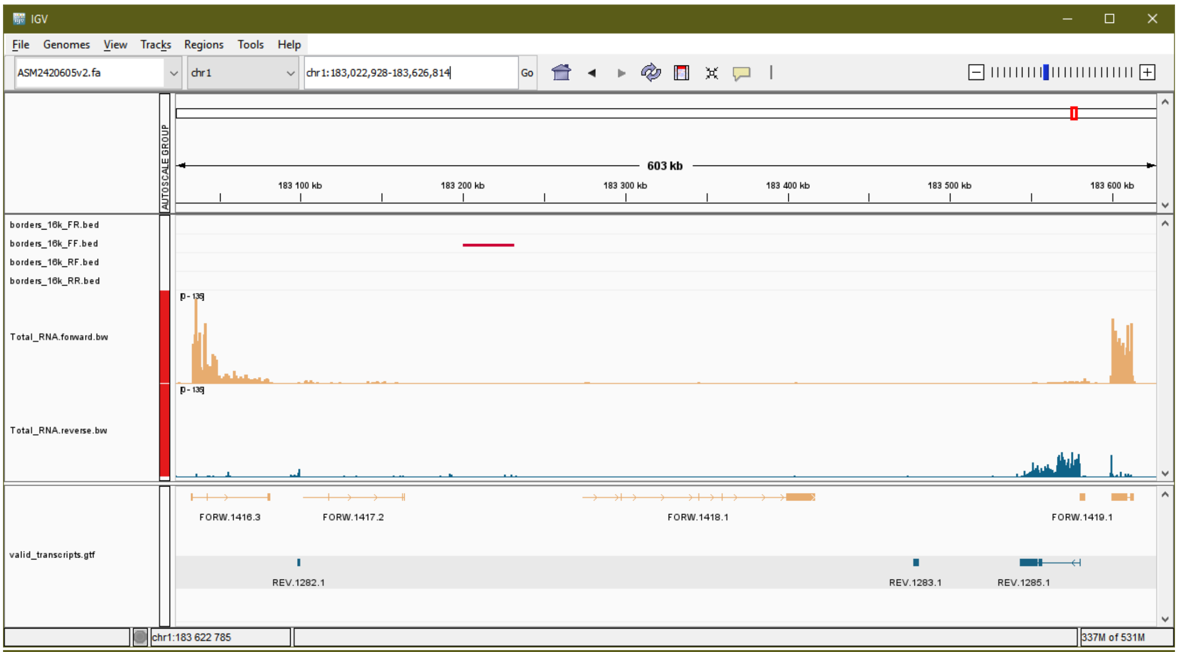
IGV
File Genomes View Tracks Regions Tools Help
ASM2420605v2.fa
chr1
chr1:183,022,928-183,626,814
Go
AUTOSCALE GROUP
603 kb
183 100 kb
183 200 kb
183 300 kb
183 400 kb
183 500 kb
183 600 kb
borders_16k_FR.bed
borders_16k_FF.bed
borders_16k_RF.bed
borders_16k_RR.bed
Total_RNA.forward.bw
Total_RNA.reverse.bw
valid_transcripts.gtf
FORW.1416.3
FORW.1417.2
FORW.1418.1
FORW.1419.1
REV.1282.1
REV.1283.1
REV.1285.1
chr1:183 622 785
337M of 531M

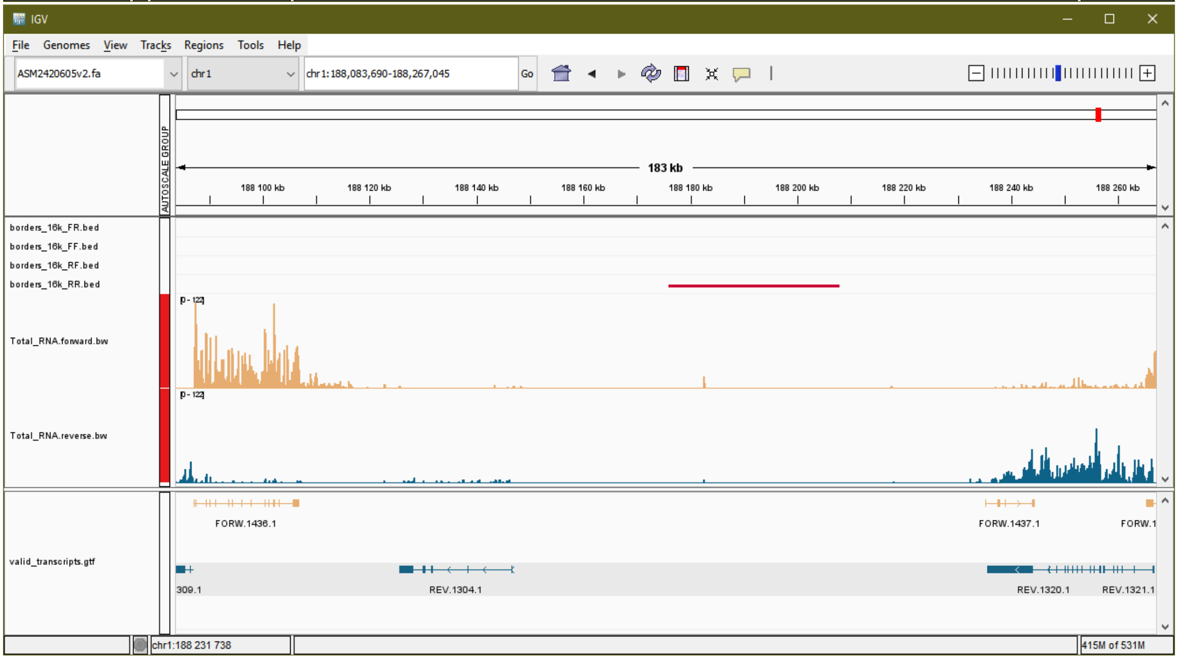
IGV
File Genomes View Tracks Regions Tools Help
ASM2420605v2.fa
chr1
chr1:188,083,690-188,267,045
Go
AUTOSCALE GROUP
183 kb
188 100 kb
188 120 kb
188 140 kb
188 160 kb
188 180 kb
188 200 kb
188 220 kb
188 240 kb
188 260 kb
borders_16k_FR.bed
borders_16k_FF.bed
borders_16k_RF.bed
borders_16k_RR.bed
Total_RNA.forward.bw
Total_RNA.reverse.bw
valid_transcripts.gtf
FORW.1436.1
FORW.1437.1
FORW.1
309.1
REV.1304.1
REV.1320.1
REV.1321.1
chr1:188 231 738
415M of 531M

**List of Supplementary Tables**

Table S1. Single nucleus Hi-C libraries quantitative characteristics for the data quality control.

Table S2. Sizes of isolated oocyte nuclei.

Table S3. The list of BAC clones containing fragments of chicken genomic DNA that were used for FISH mapping.

Table S4. Transcription activity of genes for the CTCF and different cohesin and condensin subunits.

**List of Online Supplementary Materials**

Online S1. 3D structure of the lampbrush chromosome 1 loci.

Online S2. Flattened structure of the lampbrush chromosome 1 loci.

Online S3. 3D structure of the post-lampbrush chromosome 1 loci.

Online S4. Flattened structure of the post-lampbrush chromosome 1 loci.
